# Longitudinal Impacts of Sex-Steroid Hormones on Cerebellar Structure in Adulthood are Sex Differentiated

**DOI:** 10.64898/2026.10.01.754568

**Authors:** Rushali H. Patel, Thamires N.C. Magalhães, Jessica A. Bernard

**Affiliations:** Department of Psychological and Brain Sciences, Texas A&M University, College Station, Texas, United States of America; Division of Interventional Psychiatry, The University of Texas Southwestern Medical Center, Dallas, Texas, United States of America; Texas A&M Institute for Neuroscience, Texas A&M University, College Station, Texas, United States of America

**Keywords:** sex-differences, sex-steroid hormones, cerebellum, aging, longitudinal

## Abstract

Sex-steroid hormones influence brain structure across aging, but their relationship to the cerebellum, a region implicated in cognition and vulnerable to age-related decline, remains poorly characterized, particularly longitudinally and across both sexes. We examined whether 17β-estradiol, progesterone, and testosterone relate to cerebellar volume across two timepoints, and whether these relationships differ by sex. In a mixed-sex sample of aging adults (Baseline N = 137; Year 1 N = 93), free salivary hormones and cerebellar lobular volumes were quantified. Linear regression tested cross-sectional, predictive, and change-score hormone-volume associations, and formal hormone-by-sex interactions, in *a priori* primary and secondary cerebellar lobules of interest. Hormone-cerebellum relationships were consistently sex-differentiated. Estradiol change related to Crus I volume change in opposite directions in males and females, a dissociation extending across bilateral Crus I and Lobule VI, and Right Crus II. In females, greater increases in testosterone were associated with greater decline in Left Crus I and Right Lobule V volumes. In males, testosterone showed no significant associations in primary analyses, however, increases in estradiol were associated with larger volumes. Progesterone showed a male-specific cross-sectional association with Crus II. Several relationships ran opposite to the direction predicted by estradiol’s established neurotrophic role. Sex-steroid influences on the aging cerebellum are sex-differentiated, regionally specific, and may not be predicted exclusively by cortical or female-focused models. These findings establish the cerebellum as a potential site of sex-specific hormonal influence on brain aging and also further highlight the complexity of the impact of sex-steroid hormones on the aging brain.

## Introduction

The human brain undergoes substantial changes with age. Post-mortem and in vivo imaging studies have shown that the human brain atrophies and experiences ventricular enlargement with advancing age (1, 2). However, these changes are not uniform across individuals. Sex may, in part, shape these age-related trajectories. Males display substantially larger overall brain volumes, white matter volume, and cortical surface areas, whereas females exhibit thicker cerebral cortices (3, 4). Cross-sectionally, males and females show differing unadjusted gray matter volumes in different regions of the brain with females exhibiting larger gray matter volume in the frontal areas than males and males exhibiting larger gray matter volumes in the motor and subcortical structures such as the hippocampus and amygdala (5). Longitudinally, older males showed earlier and steeper age-related declines in total brain, gray matter, and white matter, and across a greater number of brain regions; however, older females showed decline in fewer regions with limited cortical changes along the superior temporal sulcus and greater ventricular expansion (6, 7). However, findings remain mixed, with some evidence supporting greater volumetric vulnerability for females as well (8). Nonetheless, a disproportionate burden of age-related diseases, specifically Alzheimer’s disease (AD), is borne by females (9), an asymmetry that sex differences in healthy cortical aging appear insufficient to explain, pointing toward mechanisms beyond structural decline alone (7). Since females and males have distinct hormonal trajectories across adulthood (10), these differences could be further explored as a potential consequence of hormone-driven sex-specific aging.

Gonadal hormones, such as estrogen, progesterone, and testosterone, influence brain structure and function throughout adulthood (4, 11, 12), and may represent a mechanism through which sex shapes age-related brain change. Estrogen, specifically 17-ß-estradiol (referred to as estradiol hereafter) has been found to influence synaptic density (13, 14) and dendritic spine growth (15) in animal models. Regional gray matter volumes, particularly within the prefrontal cortex and the medial temporal lobe have been associated with estradiol in humans, with postmenopausal females exhibiting lower gray matter volumes in several cortical structures and subcortical structures like hippocampus, amygdala, and hypothalamus (16–18), and with exogenous estrogen manipulations showing mixed results in improving certain volumetric outcomes in females (19, 20). In females, progesterone fluctuations have been shown to modulate tissue volume (21–23) and white matter microstructural integrity (24, 25). In males, testosterone undergoes local conversion into estradiol via the aromatase enzyme (26, 27) which may contribute toward neuroprotection. Additionally, levels of total and free testosterone have been associated with different structural outcomes dependent on sex and age (28, 29). Consistent with this, the loss of ovarian hormones at menopause has been implicated in females’ elevated risk of AD (12, 30), and cumulative sex-hormone exposure interacts with the apolipoprotein E ε4 allele, the strongest common genetic risk factor for late-onset AD, in shaping brain aging (31).

The existing body of work is extensively focused on the cerebrum and relies largely on cross-sectional designs, leaving unresolved how hormonal decline shapes other hormone-sensitive regions within individuals over time. The cerebellum, by contrast, has been largely overlooked in this context, historically regarded as a motor structure though it is also implicated in cognition and vulnerable to aging (32–35). The cerebellum is also implicated in AD, showing altered synaptic processing in mice (36) and regional atrophy across disease progression (37, 38). Studies in animal models have shown that the cerebellum expresses estrogen receptors (39), progesterone receptors (40), and androgen receptors (41), and synthesizes neurosteroids locally (42). Specifically, estradiol and progesterone, found to be produced *de novo* in Purkinje cells, promote dendritic growth, spinogenesis, and synaptogenesis (43, 44), while androgen receptors in Purkinje cells were modulated by circulating testosterone (41). This cerebellar hormone signaling appears to be sexually dimorphic, with sex differences in Purkinje cell estrogen-receptor-dependent markers (45) and in progesterone-receptor regulation (40). This receptor architecture and local steroidogenesis make the cerebellum a plausible site at which different hormonal environments may give rise to sex differences in cerebellar structure and, as hormonal levels decline with age, to sex-specific patterns of cerebellar aging.

In humans, the cerebellum shows sex differences in both structure and its trajectory across aging. Males show larger cerebellar volumes than females (5, 46), a difference already evident in young adulthood (28), and sex differences in adjusted volume have since been observed across most anterior and posterior lobules, with linear age associations in both sexes (47). Longitudinal work has also identified sex differences over time in specific lobules, including Lobule VI and Crus II (48). Endogenous sex-steroid levels have also been directly linked to cerebellar organization in adults, with higher estradiol relating to greater connectivity in cognitive subregions such as Crus I and Crus II, testosterone associated with lobular cerebellar volume, and hormone connectivity relationships differing between males and females (49). Within female adulthood, exploratory work suggests regional cerebellar differences by reproductive stage, and reproductive stage has been associated with the connectivity of lobular cerebellar networks, implicating the hormonal decline of the menopausal transition (47, 50). These relationships are functionally constrained by cerebellar topography. The anterior lobules, including Lobule V, support sensorimotor processing, whereas the posterior regions, particularly Crus I and Crus II, are interconnected with prefrontal and parietal association cortices and have been observed to engage during language, working memory, and executive tasks, with Lobule VI spanning both domains (35, 51). Most of this evidence relies on cross-sectional data, with some longitudinal work documenting sex differences in cerebellar aging (48). However, longitudinal studies examining the contribution of sex-steroid hormones to these sex differences are sparse, and where hormones have been measured directly, their associations with cerebellar structure have been inconsistent (52). Whether the sex-differential cerebellar trajectories observed to date are shaped by declining sex-steroid levels remains unknown.

These cerebellar changes carry cognitive consequences. Regional cerebellar volume relates to cognitive performance in young and older adults, with age-related differences concentrated in cognitive lobules (53), and the cerebellum has been proposed as an underappreciated contributor to cognitive aging (54). Age-related cerebellar decline is also spatially heterogenous across lobules, tracking cognitive resilience in later life (55). Longitudinal cerebellar change tracks cognitive change independent of hippocampal volume (56), and this relevance extends to AD. Cerebellar atrophy progresses from vermal and anterior regions at the mild cognitive impairment stage to Crus I at dementia (37), and cerebellar volume tracks cognition even at the preclinical stage (57). Given that women bear a disproportionate burden of AD (58) that sex differences in cortical aging appear insufficient to explain (7), sex-specific cerebellar aging may be one contributing pathway. Critically, brain-cognition relationships themselves differ between older males and females (59), and sex hormones may contribute, as lower testosterone relates to poorer cognition in females (60). Whether these hormone-cerebellar relationships relate to cognitive performance, and whether this differs by sex, remains untested.

The present study addresses these gaps by examining whether free sex-steroid hormones (17β-estradiol, progesterone, and testosterone) relate to cerebellar structure longitudinally, and whether these relationships differ between males and females. Given that estradiol, progesterone, and testosterone each exert neurotrophic and neuroprotective actions, and that the cerebellum expresses receptors for all three, we hypothesized that changes in hormone levels would be associated with changes in cerebellar volume, and that hormonal decline over the follow-up interval would accompany volumetric decline. Because these hormones follow divergent trajectories in males and females across midlife and older adulthood, we expected the magnitude and these associations to differ by sex, with estradiol and progesterone relating most strongly to cerebellar structure in females, in whom they decline steeply across the menopausal transition, and testosterone relating to both sexes, given its predominance as a free sex steroid in both males and in peri- and post-menopausal females (61, 62).

We also expected these relationships to be regionally specific. Since Crus I and Crus II are densely interconnected with prefrontal and parietal association cortices, support higher-order cognition, and also show a steep age-related decline of any cerebellar region, we predicted that hormone-volume associations would be most pronounced in these subregions. Lobules V and VI, which span sensorimotor and transitional functions, were also examined as secondary regions to determine whether any hormone-cerebellum relationships were specific to just the cognitive regions of the cerebellum or if they extended across its functional divisions. Post-hoc, we examined whether hormone-related differences in cerebellar volume were associated with cognitive performance, and whether these associations differed by sex.

## Methods

### Participants

The data for this study are part of a larger investigation into longitudinal changes in the cerebellum across adulthood. At each of two timepoints, participants completed cognitive testing and provided saliva samples for hormone quantification, followed by a magnetic resonance imaging (MRI) session. Our group has previously reported on data drawn from this larger study (49, 63).

Participants were enrolled on a rolling basis. Baseline assessments were conducted between January 2020 and September 2022, and Year 1 follow-up assessments between February 2021 and July 2023. Because baseline data collection began at the onset of the COVID-19 pandemic, the interval between saliva collection and imaging, and between baseline and Year 1, varied across participants owing to pandemic-related scheduling disruptions. Participants returned for the Year 1 follow-up after a median of 366 days for saliva collection (range: 345-750 days) and 409 days for structural imaging (range: 173-759 days).

Exclusion criteria included a history of neurological disease, stroke, or diagnosed psychiatric illness, contraindications for the MRI environment, and use of hormone therapy or hormonal contraceptives; female participants with a hysterectomy within the preceding ten years were also excluded, so that analyses reflected normative endocrine aging. All procedures were approved by the Institutional Review Board at Texas A&M University, and all participants provided written informed consent.

At baseline, the sample comprised 137 participants (73 female, 64 male). Reproductive stage was determined at both timepoints using Stages of Reproductive Aging Workshop +10 (STRAW+10) criteria (64), following the classification applied in prior work from our group (47, 50); participants for whom the available information was insufficient to assign a stage were categorized as unknown (*n* = 3) and excluded from reproductive stage analyses. No participant included in the present analyses transitioned reproductive stage across the follow-up interval. Of the 137 baseline participants, 93 (49 female, 44 male) had usable structural data at Year 1 and were included in the longitudinal analyses. The mean scan-to-scan interval was 1.14 years (*SD* = 0.28; range 0.47–2.08), and the mean saliva-to-saliva interval was 1.07 years (*SD* = 0.18; range 0.95–2.05). Of these, 88 had complete hormone data across all three assays; the remainder were missing one or more hormone values owing to insufficient sample volume or assay-related issues. Since hormone assays and imaging measures were subject to independent missingness, the number of participants included in each analysis model varied with that particular model’s data requirements, and the analytic *N* is reported alongside each result. All participants were right-handed. Sample characteristics are also summarized in Table 1.

**Table 1:** Demographic and reproductive characteristics of the sample at Baseline and Year 1.

| Characteristic | Baseline<br>(*n* = 137) | Year 1<br>(*n* = 93) |
| --- | --- | --- |
| *N* | 137 | 93 |
| Age, *M* ± *SD* (range) | 56.7 ± 13.3 (35–86) | 59.0 ± 13.1 (36–87) |
| Education (years), *M* ± *SD* (range) | 17.1 ± 3.0 (12–28) | 17.3 ± 3.1 (12–28) |
| **Race, *n* (%)** |  |  |
| White | 117 (85.4) | 84 (90.3) |
| Asian | 10 (7.3) | 4 (4.3) |
| Black or African American | 3 (2.2) | 1 (1.1) |
| More than one race | 6 (4.4) | 3 (3.2) |
| American Indian/Alaska Native | 1 (0.7) | 1 (1.1) |
| **Ethnicity, *n* (%)** |  |  |
| Hispanic or Latino | 16 (11.7) | 11 (11.8) |
| Not Hispanic or Latino | 118 (86.1) | 79 (84.9) |
| Unknown / Not reported | 3 (2.2) | 3 (3.2) |
| **Sex, *n* (%)** |  |  |
| Male | 64 (46.7) | 44 (47.3) |
| Female | 73 (53.3) | 49 (52.7) |
| **Reproductive stage (of females), *n* (%)** |  |  |
| Reproductive | 18 (24.7) | 11 (22.4) |
| Early Postmenopausal | 14 (19.2) | 10 (20.4) |
| Late Postmenopausal | 37 (50.7) | 27 (55.1) |
| Hysterectomy | 1 (1.4) | 1 (2.0) |
| Unknown | 3 (4.1) | 0 (0.0) |
| **MoCA, *M* ± *SD* (range)** |  |  |
| Full sample | 26.7 ± 2.1 (20–30) | 27.2 ± 2.3 (20–30) |
| Male | 26.3 ± 2.1 (20–30) | 26.7 ± 2.6 (20–30) |
| Female | 27.1 ± 2.1 (22–30) | 27.6 ± 2.0 (22–30) |

*Note.* *M* = mean; *SD* = standard deviation; MoCA = Montreal Cognitive Assessment; Unknown = female participants whose reproductive stage could not be determined from the available data. Reproductive stage percentages are computed relative to the number of female participants at each timepoint. Age at baseline and at Year 1 reflect participant age at each respective assessment.

Eleven participants were identified as hormonal outliers using a criterion of 3 standard deviations (*SD*) applied within sex and timepoint. These participants were retained in all primary analyses. Our rationale for this is that extreme but valid hormone values are still characteristic of the population under study rather than exclusively measurement errors, and that their exclusion would compromise generalizability. All primary analyses were additionally repeated with these participants removed; findings surviving this procedure are described as robust.

### Hormone quantification

Saliva samples were collected following the methods detailed in our recent work (49) at each timepoint using the passive drool technique into pre-labeled cryovials (Salimetrics SalivaLab, Carlsbad, CA), with participants providing approximately 1 mL per sample. Samples were stored at −80°C until assay. Salimetrics quantified free salivary 17β-estradiol, progesterone, and testosterone levels, with assay sensitivities of 0.1 pg/mL for 17β-estradiol (65), 0.458 pg/mL for testosterone (66), and 5.0 pg/mL for progesterone (67). For samples in which a replicate fell below the assay sensitivity and was reported as non-detectable, that replicate was imputed as the sensitivity value divided by √2 (68) before averaging replicates into the sample level. This applied to 17β-estradiol and testosterone; no progesterone replicate fell below detection, so no imputation was required.

### Image acquisition

Structural MRI was acquired on a Siemens Magnetom Verio 3.0T scanner with a 32-channel head coil. A high-resolution T1-weighted 3D MPRAGE sequence was collected (TR = 2400 ms; voxel size = 0.8 mm³; multiband acceleration factor = 2), along with a high-resolution T2-weighted scan (TR = 3200 ms; voxel size = 0.8 mm³). Acquisition protocols were adapted from the Human Connectome Project (69) to support data sharing and reproducibility. The full acquisition protocol for the complete scanning session has been described previously (49); the present analyses used only the structural T1- and T2-weighted images.

### Image processing and cerebellar segmentation

T1-weighted images were converted from DICOM to NIfTI format and processed using FastSurfer (70, 71) together with FreeSurfer version 7.4.1 for cortical and subcortical volumetric segmentation. Cerebellar lobular volumes were obtained using CerebNet (72), a fully automated, validated deep-learning tool for cerebellar sub-segmentation integrated within the FastSurfer pipeline. Bilateral volumes were extracted for each cerebellar region of interest (ROI), following the approach used in our recent work (49).

### Cognitive measures

Three cognitive measures were examined in relation to cerebellar volume. Global cognitive status was assessed with the Montreal Cognitive Assessment (MoCA; 73), a widely used screening instrument spanning attention, executive function, memory, language, visuospatial ability, and orientation. Processing speed was indexed by the Digit Symbol Substitution task from the Wechsler Adults Intelligence Scale, 4^th^ Edition (WAIS-IV–Coding; 74), in which participants translate symbols to digits according to a key within a fixed time limit, with the number of correct substitutions serving as the score. Working memory was assessed with a composite derived from the forward, backward, and sequencing Digit Span conditions (74, 75) each standardized (*z*-scored) across the full sample within each timepoint and averaged to yield a working-memory composite. All cognitive measures were administered at both timepoints; baseline values were used in the analyses reported here.

### Statistical Analyses

All analyses were conducted in R (version 4.4.1; R Core Team) using RStudio (version 2024.09.0+375). Sex-steroid hormone levels (17β-estradiol, progesterone, testosterone) were examined in relation to cerebellar volume using linear regression. Analyses were conducted in the full sample and separately by sex. All volumetric models included estimated total intracranial volume, age, years of education, and (in full-sample analyses) sex as covariates; sex was necessarily omitted in sex-stratified models. Longitudinal prediction models were controlled for baseline volume. These predictive models tested whether baseline hormone level was associated with Year 1 volume; the primary longitudinal analysis was the change-score model described below. All findings reported below derive from the full sample with outliers retained; sensitivity analyses excluding outlying values are summarized within each subsection, with complete results for all analyses, including limbic, cortical, and reproductive-stage findings, reported in Supplementary Materials (see Tables S3-S8, S16-S19).

Consistent with our longitudinal hypotheses, our primary analyses tested whether change in hormone from baseline to Year 1 was associated with change in cerebellar volume, and whether these change relationships differed by sex. Change was quantified as the simple difference between Year 1 and baseline values. Crus I and Crus II (Left and Right) were designated *a priori* primary regions of interest on anatomical and functional grounds (35, 49, 51), and were evaluated at *p* < .05 without correction for multiple comparisons; this targeted threshold reflects their pre-specification rather than data-driven selection. Lobules V and VI (left and right) were examined as secondary cerebellar regions to determine whether any relationships extended beyond the cognitive cerebellum. Limbic regions were examined as secondary regions, and remaining regions as exploratory. Secondary and exploratory regions were corrected using the Benjamini-Hochberg false discovery rate (FDR) procedure (76) applied within region family and within analysis type.

To formally test whether hormone-volume change relationships differed by sex, models including a hormone × sex interaction term were fit, evaluated at *p* < .05 uncorrected for primary regions and FDR-corrected for secondary regions. Sex was coded with females as the reference group, such that interaction terms represent the difference in the hormone-volume slope for males relative to females. To contextualize the change-score findings, we additionally examined cross-sectional hormone-volume associations at baseline and Year 1, and whether baseline hormone level predicted Year 1 volume (controlling baseline volume). These analyses inform interpretation of the longitudinal results but were not themselves hypothesis-driven; in these analyses, the same correction structure was applied, with Crus I and Crus II evaluated uncorrected as *a priori* regions and all other regions FDR-corrected within family. Sensitivity analyses additionally modeled Year 1 volume as a function of hormone change and baseline hormone level, controlling for baseline volume, to address the possibility that change-score associations reflected regression to the mean. These are reported in Supplementary Materials (see Tables S10-S11). To confirm variation in follow-up interval did not account for the observed relationships, all change-score analyses were repeated with the between-session intervals (scan-to-scan and saliva-to-saliva) included as a covariate, and cross-sectional analyses were repeated with the within-session interval between the scan and saliva collection dates as a covariate. Correction structure mirrored that of the primary analyses. Additionally, robustness checks for the estradiol × sex dissociation, alternative volume normalization and global cerebellar control were conducted (see Table S9). Results are reported in Supplementary Materials (see Tables S12-S15).

To examine whether hormone-volume relationships related to cognitive performance, we tested hormone × cognition interactions on volume, using MoCA, processing speed (Digit Symbol), and the working-memory composite. These analyses were post-hoc and FDR-corrected within region family (primary cerebellar; secondary cerebellar; and global volumes, comprising total gray matter, cortical, and cerebral white matter volume to test whether any hormone x cognition associations were specific to the cerebellum or extended to the global brain structure). Exploratory analyses stratifying the female sample by reproductive stage were also conducted; given the small stage subgroup sizes, these are reported in Supplementary Materials (see Tables S16-S19) and interpreted as preliminary.

## Results

Descriptive statistics for hormone levels and cerebellar volumes, by sex and timepoint, are presented in Table S1. The analyses comprised cross-sectional associations at baseline and Year 1, baseline hormone predicting Year 1 volume, and change-score associations, together with formal tests of whether change relationships differed by sex.

### Estradiol

#### Cross-sectional and prediction analyses

At baseline, higher estradiol was associated with smaller Right Crus I volume in females (*b* = −468.2, *p* = .021). At Year 1, this negative association appeared bilaterally in females, in Right Crus I (*b* = −698.7, *p* = .035) and Left Crus I (*b* = −590.4, *p* = .035), and in the full sample, in Right Crus I (*b* = −468.0, *p* = .032) and Left Crus I (*b* = −463.6, *p* = .023). Baseline estradiol did not predict Year 1 Crus I or Crus II volume in any group, though it predicted larger Year 1 Right Lobule VI in females (*b* = 117.3, *p*_FDR = .038; Table S2).

#### Change scores

Estradiol change related to Left Crus I change in opposite directions by sex. In females, an increase in estradiol accompanied a reduced Left Crus I volume (*b* = −116.9, *p* = .037), whereas in males, an increase in estradiol accompanied a larger Left Crus I volume (*b* = 189.9, *p* = .020; Figure 1). Increases in estradiol were also associated with larger Right Lobule VI volume in males (*b* = 146.1, *p*_FDR = .017; Table 2).

**Figure 1.**
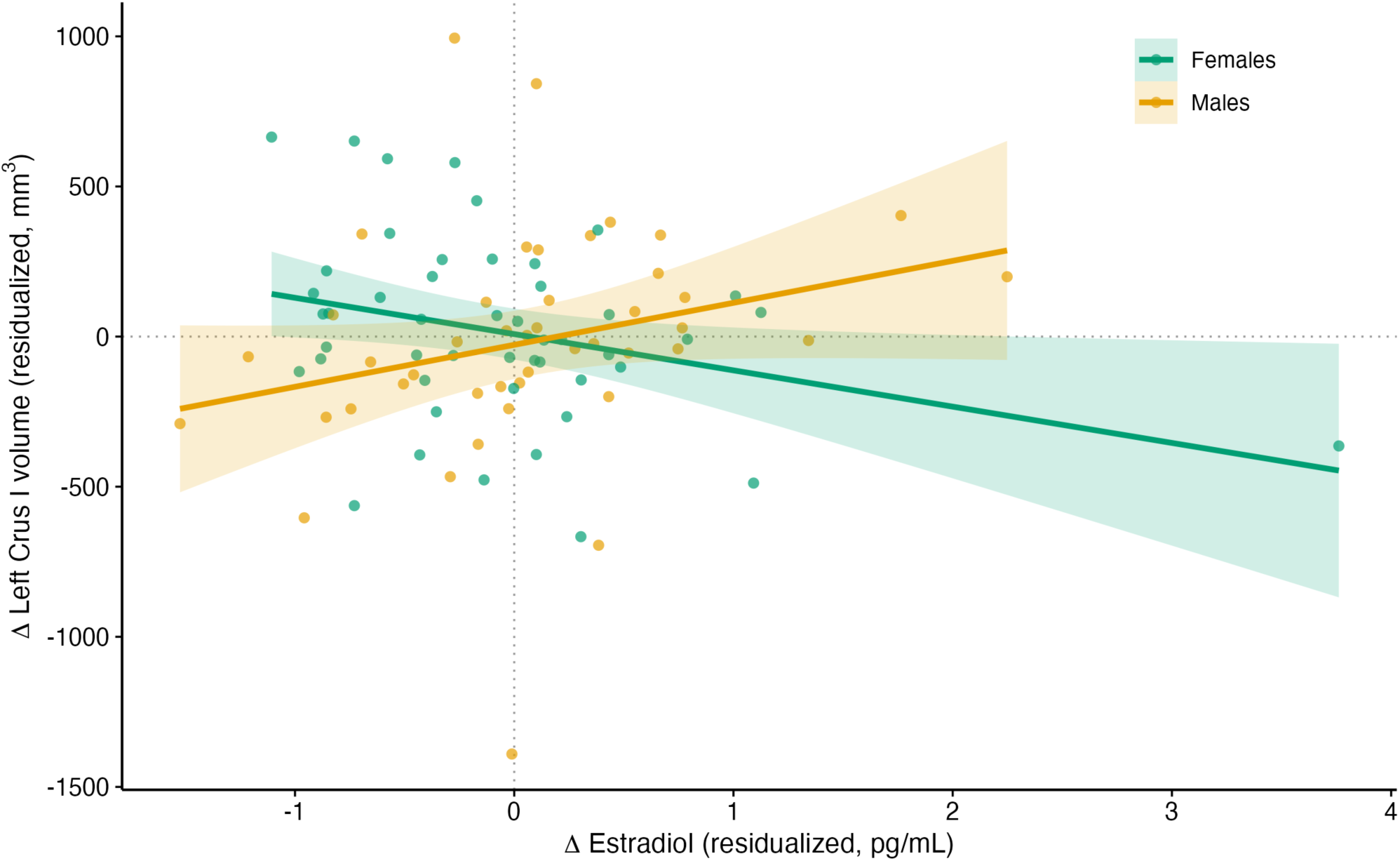
Estradiol-Left Crus I Change-Score Association by Sex *Note.* Change in Left Crus I volume as a function of estradiol change, shown separately for females (teal) and males (orange). Females displayed a negative relationship, where an increase in estradiol was associated with a reduction in volume. Males showed an opposite relationship, with an increase in estradiol associated with an increase in volume. Points represent individual participants; shaded bands indicate 95% confidence intervals. Both axes are residualized for age, education, and estimated total intracranial volume.

**Table 2:**
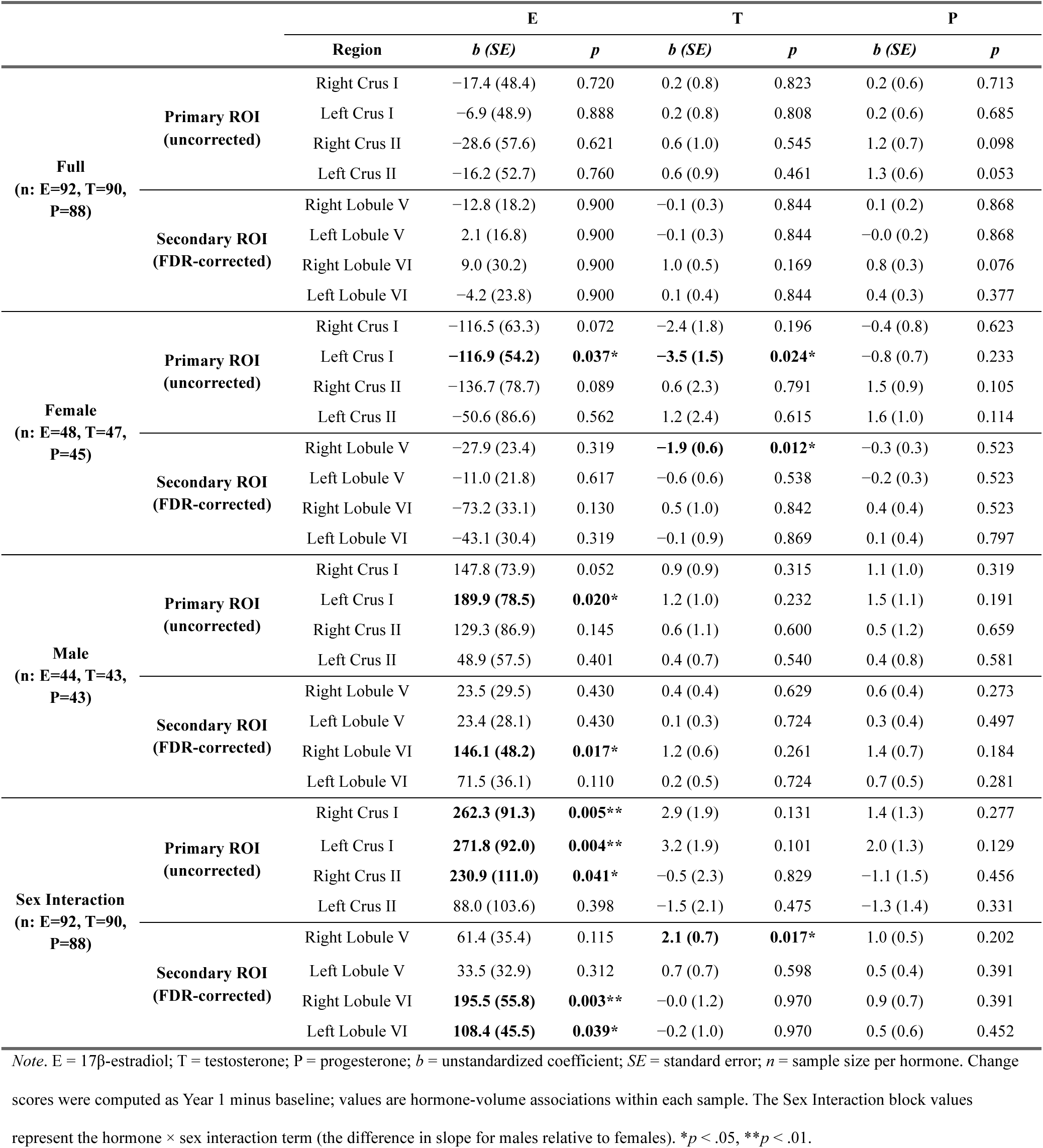
Hormone-cerebellar change-score associations with sex interactions.

#### Sex interactions

The estradiol × sex interaction was significant for volumetric change in Left Crus I (*b* = 271.8, *p* = .004), Right Crus I (*b* = 262.3, *p* = .005), and Right Crus II (*b* = 230.9, *p* = .041). It extended to bilateral Lobule VI, in the right (*b* = 195.5, *p*_FDR = .003) and left (*b* = 108.4, *p*_FDR = .039) hemispheres.

#### Sensitivity analyses

The male change-score effects and all estradiol × sex interactions held with outlying values excluded, and the interaction additionally extended to Right Lobule V. The female cross-sectional and change-score associations were no longer significant when outliers were excluded. All primary and secondary estradiol effects also held when the between-session interval was included as a covariate; the female cross-sectional associations, though no longer significant by outlier exclusion, were unchanged by adjustment for the interval between the scan and saliva collection dates.

### Testosterone

#### Cross-sectional and prediction analyses

Testosterone showed no significant cross-sectional or predictive associations with cerebellar volume.

#### Change scores

In females, increases in testosterone accompanied a reduced Left Crus I volume (*b* = −3.5, *p* = .024). Among secondary cerebellar regions, increases in testosterone accompanied a reduced Right Lobule V volume in females (*b* = −1.9, *p*_FDR = .012; Figure 2). No change-score associations were significant in males.

**Figure 2.**
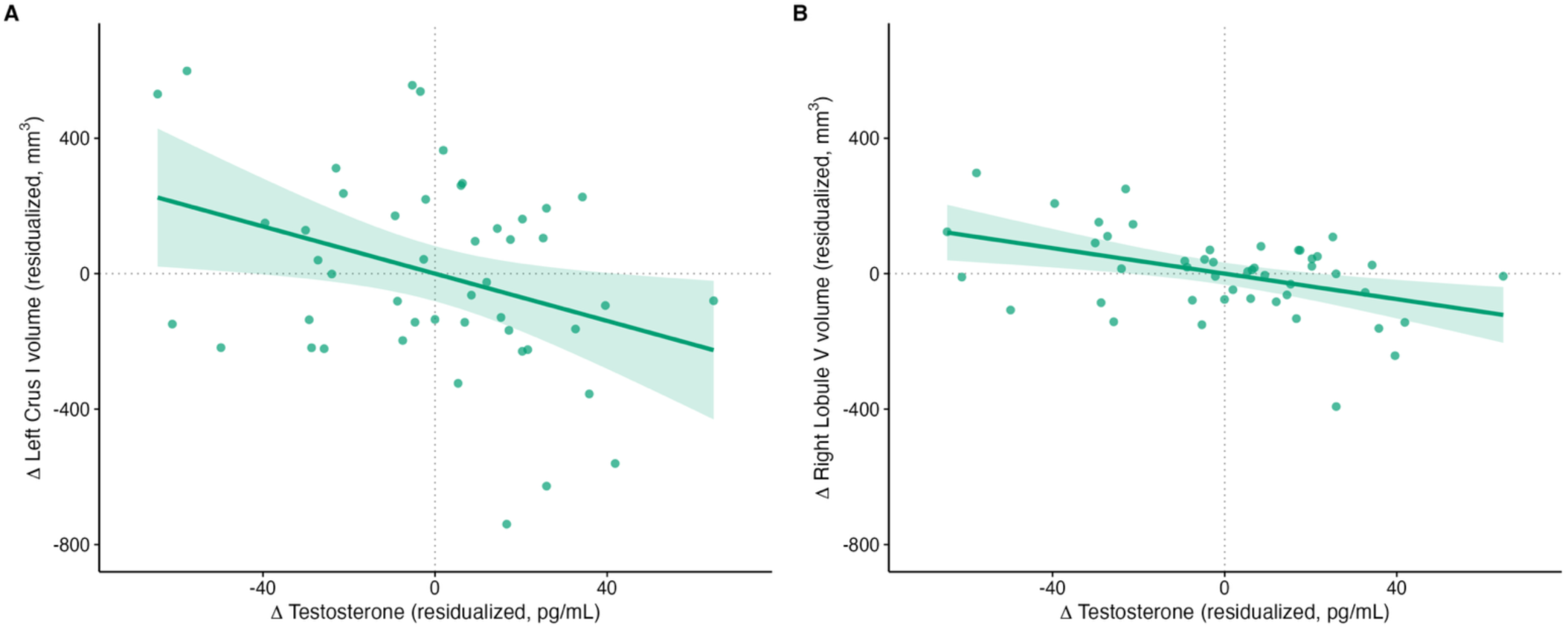
Testosterone Change Associations With Left Crus I and Right Lobule V in Females *Note.* Change in (A) Left Crus I and (B) Right Lobule V volume as a function of testosterone change in female participants (teal). In both regions, an increase in testosterone was associated with a reduction in volume. Points represent individual female participants; shaded bands indicate 95% confidence intervals. Axes are shared across panels, and both axes are residualized for age, education, and estimated total intracranial volume.

#### Sex interactions

The testosterone × sex interaction was significant for Right Lobule V change (*b* = 2.1, *p*_FDR = .017). The interaction did not reach significance for Left Crus I, indicating that the Left Crus I association, while significant in females, was not formally sex-differentiated.

#### Sensitivity analyses

Both female change-score associations, and the Right Lobule V sex interaction, strengthened when outlying values were excluded, and all held when the between-session interval was included as a covariate. Sensitivity analyses additionally revealed male-specific associations were not present in the full sample. At baseline, higher testosterone was associated with larger Right Crus II (*b* = 7.5, *p* = .029), and baseline testosterone predicted smaller Year 1 Right Crus I (*b* = −3.6, *p* = .004).

### Progesterone

#### Cross-sectional and prediction analyses

Higher baseline progesterone was associated with smaller Left Crus II volume in males (*b* = −7.7, *p* = .025; Figure S1). In the full sample, baseline progesterone was negatively associated with Right Crus I (*b* = −4.5, *p* = .023). At Year 1, progesterone was negatively associated with bilateral Crus I in the full sample, in Right Crus I (*b* = −5.0, *p* = .014) and Left Crus I (*b* = −4.2, *p* = .027). Baseline progesterone did not predict Year 1 volume.

#### Change scores

Change in progesterone was not associated with cerebellar volumetric change.

#### Sex interactions

In the full sample, the progesterone × sex interaction was significant for bilateral Crus II, in Left Crus II (*b* = −8.1, *p* = .020) and Right Crus II (*b* = −7.2, *p* = .049) at baseline. Together with the male-specific cross-sectional association, this indicates that the progesterone-Crus II relationship differed by sex, being driven by males.

#### Sensitivity analyses

The male Left Crus II association and the Left Crus II sex interaction strengthened when outlying values were excluded, and the cross-sectional progesterone associations were unchanged when the within-session interval between the scan and saliva collection dates was added as a covariate. Sensitivity analyses additionally revealed isolated associations not present in the full sample, in male bilateral Crus I prediction, and female Left Crus I change.

### Post-hoc cognitive analyses

Given that cerebellar structure relates to cognitive performance, we tested whether hormone-cerebellar relationships were moderated by cognition. Baseline estradiol interacted with MoCA to predict Year 1 Left Lobule V volume in the full sample (*b* = 32.5, *p*_FDR = .032) and, more strongly, in males (*b* = 63.3, *p*_FDR = .026); at higher MoCA scores, the estradiol-Lobule V association became more positive (Figure 3). At the global level, estradiol also interacted with MoCA to predict Year 1 total gray matter (*b* = −2406.1, *p*_FDR = .006) and cortical volume in the full sample (*b* = −2180.2, *p*_FDR = .006). In females, this global effect sustained in total gray matter (b = −3064.2, *p_FDR* = .008) and in cortical volume (*b* = −2820.1, *p*_FDR = .006), in the negative direction. Testosterone change interacted with MoCA to predict Right Lobule V change in females (*b* = −0.96, *p*_FDR = .015), such that the negative testosterone-Lobule V relationship was more pronounced at higher MoCA scores. Processing speed and the working-memory composite yielded no significant cerebellar interactions.

**Figure 3.**
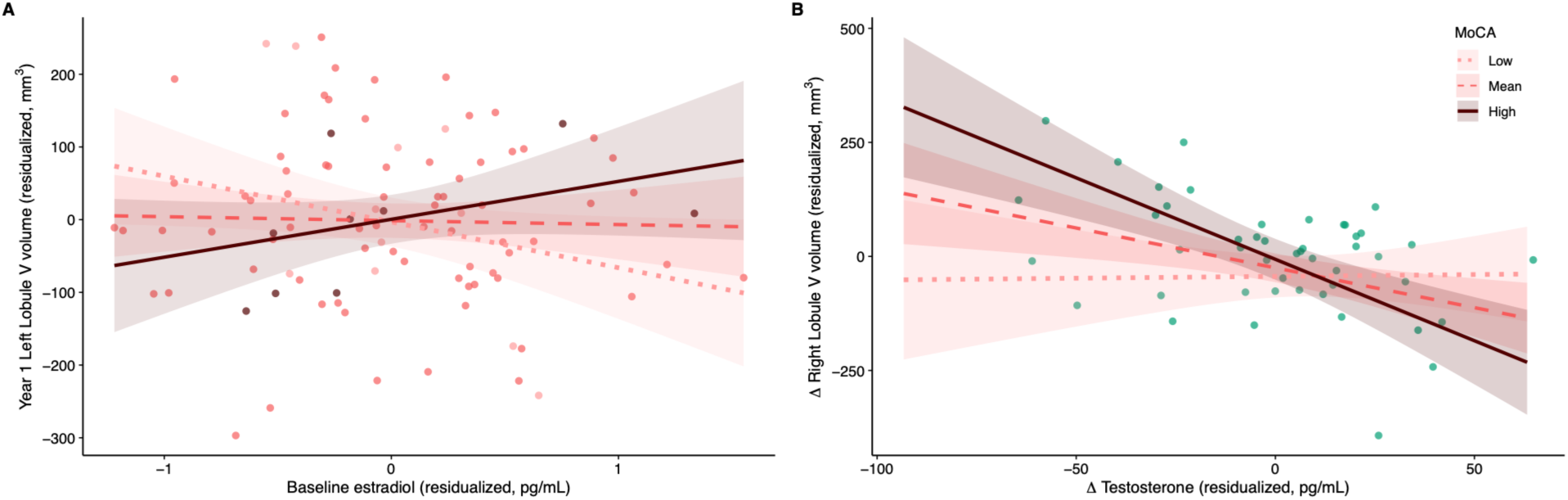
Hormone-Cerebellar Associations Moderated by MoCA *Note.* Simple slopes of the hormone-volume associations at low, mean, and high MoCA (mean ± 1 *SD*). Hormones and volumes were residualized for the model covariates. Line shades denote low to high MoCA (light to dark). Shaded bands indicate 95% confidence intervals. (A) Baseline estradiol predicting Year 1 Left Lobule V volume in the full sample; at higher MoCA scores, the estradiol-volume relationship became more positive (MoCA levels: low = 24.9, mean = 26.9, high = 29.0); adjusted for baseline volume, age, sex, education, and estimated total intracranial volume. Points represent individual participants, shaded by MoCA level (light to dark for low to high). (B) In females, the negative association between testosterone change and change in Right Lobule V volume was more pronounced at higher MoCA scores (MoCA levels: low = 25.5, mean = 27.4, high = 29.3). Points represent individual female participants (teal); adjusted for age, education, and estimated total intracranial volume.

#### Sensitivity analyses

The estradiol × MoCA effects on Left Lobule V and on global volumes held when outlying values were excluded. A hormone-independent association between MoCA and Left Lobule VI, and an estradiol × processing-speed association with Right Crus II change, emerged only in outlier-excluded analyses.

### Exploratory analyses

We conducted exploratory sex-stratified and reproductive-stage analyses, and examined exploratory regions including the vermis and global volumes. Reproductive-stage subgroups were in several cases too small for stable model estimation, for example *n* = 11 in the reproductive group, and these analyses are not reported. Among adequately powered exploratory analyses, notable patterns included a strong negative association between progesterone and bilateral Lobule V in early postmenopausal females at baseline (Figure S2), and a negative association between progesterone and Vermis X volume in females at Year 1 (Figure S3). Exploratory limbic (hippocampus, amygdala, and thalamus) and global analyses further indicated that estradiol was negatively associated with left hippocampus and right amygdala volumes at baseline in males and positively associated with cerebral white matter volume at baseline in females. These findings are preliminary and require confirmation in adequately powered samples; results for the exploratory analyses are reported in the Supplementary Materials (see Tables S3-S8, S16-S19).

## Discussion

This study examined whether sex-steroid hormones relate to cerebellar structure longitudinally, and whether these relationships differ between males and females in a mixed-sex sample of aging adults. Across estradiol, testosterone, and progesterone, and across cross-sectional, predictive, and change-based analyses, hormone-cerebellum relationships were region-specific, sex-dependent, and in several cases opposite in direction to what a literature grounded in cross-sectional, cerebral, and female-focused work would posit. The effects concentrated in Crus I, Crus II, and Lobules V and VI, the cerebellar regions most engaged in higher-order function and vulnerable to age-related decline, while the Lobule-IX control region showed no associations. Additional findings for global cortical and subcortical areas are highlighted in the Supplementary Materials (see Tables S3, S5, and S7). In the sections that follow, we consider each hormone in turn before drawing together the sex-differentiated pattern that distinguishes these findings, and its implications for how the aging cerebellum is understood.

Table 3 summarizes the significant change-score associations and hormone × sex interactions discussed below.

**Table 3:** Summary of significant change-score associations and sex interactions.

| Hormone | Region | Change-score association | Sex interaction |
| --- | --- | --- | --- |
| E | Left Crus I | Females: $-116.9$ ($p = .037$);<br>Males: $189.9$ ($p = .020$) | Significant ($b = 271.8$, $p = .004$) |
| E | Right Crus I | — | Significant ($b = 262.3$, $p = .005$) |
| E | Right Crus II | — | Significant ($b = 230.9$, $p = .041$) |
| E | Right Lobule VI | Males: $146.1$ ($p_{\text{FDR}} = .017$) | Significant ($b = 195.5$, $p_{\text{FDR}} = .003$) |
| E | Left Lobule VI | — | Significant ($b = 108.4$, $p_{\text{FDR}} = .039$) |
| T | Left Crus I | Females: $-3.5$ ($p = .024$) | Not significant |
| T | Right Lobule V | Females: $-1.9$ ($p_{\text{FDR}} = .012$) | Significant ($b = 2.1$, $p_{\text{FDR}} = .017$) |
| P | — | No significant change-score<br>associations | — |

*Note.* E = $17\beta$ -estradiol; T = testosterone; P = progesterone; $b$ = unstandardized coefficient;
Dashes indicate no significant effect of that type in that region. Primary cerebellar regions (Crus I, Crus II) were evaluated at $p < .05$ uncorrected per the a priori hypothesis; secondary regions (Lobules V, VI) were FDR-corrected ( $p_{\text{FDR}}$ reported). Progesterone additionally showed a significant sex interaction cross-sectionally at baseline in bilateral Crus II (Left $b = -8.1$ , $p = .020$ ; Right $b = -7.2$ , $p = .049$ ), driven by males; as a cross-sectional effect it falls outside the change-score/interaction scope of this table.

### Estradiol

The estradiol findings diverge from the hypothesis. Estradiol’s neurotrophic and neuroprotective actions (77), together with its steep decline across the menopausal transition (31, 64), predict that lower estradiol would accompany reduced cerebellar volume in females, and thus a positive association between estradiol and volume. However, the observed relationship was the opposite. Higher estradiol was associated with smaller Right Crus I volume in females at baseline, an association that appeared bilaterally at Year 1. To our knowledge, a direct association between estradiol and Crus I volume has not been previously reported, and the negative direction in the present study does not reconcile with the existing neurotrophic account of estradiol’s role in the female brain.

The observed estradiol change points toward a mechanism that potentially differs from the current consensus in aging females. Estradiol is characterized by an increasing variability and eventual low, unstable levels rather than a smooth declining trajectory across the menopausal transition (64). An increase in estradiol in a peri- or post-menopausal female may therefore signal hormonal instability which may then reflect a broader estradiol-specific endocrine dysregulation associated with the volumetric change observed in the present study. This, however, is not evidence that estradiol is detrimental and rather there is a possibility that the negative association between estradiol and cerebellar volumes in females signifies a process distinct from the prevailing neurotrophic framing. The cross-sectional finding highlighted in the study, however, counsels caution in its interpretation since the effect did not survive outlier removal in sensitivity checks. Given the decision to retain outliers for all the analyses in the present study due to the inherent variability in hormonal levels in females, the results still hold a nuance worth exploring further.

The more robust estradiol signal emerged in the longitudinal and sex-interaction analyses (Table 3). In these analyses, estradiol change was related to Left Crus I volume change in opposite directions in males and females: in females, increases in estradiol were associated with a reduced Left Crus I volume, whereas in males, increases in estradiol were associated with a greater Left Crus I volume. This male effect was sustained across sensitivity analyses. The sex interaction spanned bilateral Crus I, Right Crus II, and extended to Lobule VI among secondary regions. This positive association of estradiol increase with greater Left Crus I volume diverges from the common framing of estradiol as a predominantly female hormone, and calls for a mechanism specific to the male endocrine context.

Aromatization is one such mechanism. Despite low levels of circulating estradiol in males, a substantial fraction of the estradiol acting within the male brain is synthesized locally from testosterone by aromatase (26, 27). This enzyme is highly expressed in cerebellar tissue along with estrogen receptors (39, 78). Measured estradiol in males may therefore reflect this local conversion, and higher estradiol may index greater aromatase activity in cerebellar tissue equipped to respond to the resulting estradiol (44). This interpretation may connect two findings that are otherwise puzzling in isolation. It offers a potential explanation for why estradiol is the hormone coupled to male cerebellar structure and not testosterone, and possibly why the male testosterone change analyses in the present study returned no significant results. This does not render testosterone irrelevant to the male cerebellum, but it may indicate that its structural effect is likely carried by the estrogenic product of aromatization rather than by testosterone directly.

This poses a potentially direct and clinically crucial implication. If the impact of testosterone on the cerebellar structure is through the aromatized estradiol, then current interventions aimed at raising testosterone while simultaneously suppressing its conversion to estradiol may not align with the association observed here. This is relevant to testosterone supplementation, which is increasingly prescribed in aging males (79), and to the use of aromatase inhibitors alongside it. Aromatization of testosterone has been shown to be required for some of its physiological effects in aging males, including maintenance of bone mineral density (80), establishing that testosterone’s benefits can depend on its conversion to estradiol rather than on testosterone itself. The present findings, however, do not answer the question of whether the neurostructural association observed here depends on aromatization in a way that suppressing this conversion during supplementation would have a negative effect. Future controlled studies measuring both hormones could potentially resolve this tension and offer a better understanding for safer and optimal supplementation routines.

### Testosterone

Testosterone showed no cross-sectional or predictive associations with cerebellar volume, and its relationships emerged instead in the change-score analyses, and specifically in females. Increases in testosterone accompanied decreases in Left Crus I volume, and, among secondary cerebellar regions, decreases in Right Lobule V volume. Both female associations were among the most stable in the study, strengthening rather than weakening when outlying values were excluded, and the Lobule V relationship was further supported by a significant sex interaction. In males, testosterone change was unrelated to cerebellar volume throughout.

The negative direction of these associations diverges from a typical neurotrophic account, which postulates that higher testosterone would be expected to accompany a greater volume preservation (81). Instead, an increase in testosterone, in the current sample, was associated with a volume reduction in females, suggesting that testosterone’s relationship to the female cerebellum may depend on the context in which the androgen is elevated. In females, testosterone remains as the predominant free sex steroid across peri- and postmenopausal stages due to the steady decline in estradiol (62), and this testosterone-dominant environment may align more toward an androgenized hormonal profile. The negative association observed here may then reflect an atypical hormonal environment when androgen excess occurs, and likely different from an environment typically expected to be neuroprotective. However, this does not automatically mean that testosterone has a broadly detrimental effect on the cerebellum. This finding may instead be better understood as identifying the specific vulnerability that elevated androgen poses on the female cerebellum, particularly across mid- and later-life.

A combination of the interpretation highlighted above and the absence of any testosterone relationship in males is informative when presented alongside the estradiol findings observed in same sex. The process of aromatization of testosterone into estradiol may be a driving factor of the structural effect of testosterone availability on the cerebellum. Testosterone, and its conversion to estrogen in the brain, thus potentially displays different relationships to cerebellar structure in the two sexes with testosterone significantly affecting volume in females and estradiol being significant for cerebellar structure in males, a dissociation that may be obscured by considering either hormone alone. Because of the marked changes in estradiol and progesterone associated with the menopausal transition in females, these hormones (along with follicle stimulating hormone) have been of particular interest (16, 82), while testosterone has been less of a focus. Our results suggest here that the broader hormonal milieu, including androgens is a key consideration in the context of the cerebellum.

These findings also motivate targeted investigation of conditions characterized by sustained androgen elevation in females. Higher testosterone change was associated with cerebellar volume reduction which raises the question of whether hyperandrogenic conditions, most notably polyendocrine metabolic ovarian syndrome (PMOS; formerly polycystic ovary syndrome;, 83), are accompanied by accelerated cerebellar aging. Structural neuroimaging in PMOS has largely concentrated on cortical, limbic, and white-matter measures (84), and the studies that have quantified the cerebellum have treated it as a single ROI within whole-brain analyses, reporting null cross-sectional differences in reproductive-age females (85, 86). Since the associations observed in the present study were regionally specific, especially concentrated in Crus I and Lobule V rather than distributed across the entire cerebellum, analyzing the cerebellum as whole structure could potentially obscure region-specific effects. However, the present study is not equipped to interpret whether the altered subregional cerebellar structure in females is a result of androgen excess as it examined hormonal variation rather than a chronic androgenic hormonal profile. Future studies, taking the regional-specificity of cerebellar lobules into account, in hyperandrogenic females could directly examine whether the direction identified here extends to sustained androgenic profile. There would be further benefit to the inclusion of longitudinal approaches in those with PMOS to determine the degree to which exposure to a chronic androgenic profile impacts cerebellar structure across adulthood and into aging.

### Progesterone

Progesterone was associated with cerebellar volume cross-sectionally but not longitudinally. Higher baseline progesterone was associated with smaller Left Crus II volume in males, an association that strengthened when outlying values were excluded, and with smaller Right Crus I volume in the full sample, extending bilaterally at Year 1. A significant progesterone by sex interaction in bilateral Crus II indicated that the Crus II relationship differed by sex, driven by males. Progesterone did not predict future volume, and progesterone change was unrelated to volumetric change in any group.

This pattern was the least anticipated of the three hormones. Progesterone, like estradiol, declines across the menopausal transition (64), and the present study was framed around the expectation that its decline would relate to cerebellar structure in females longitudinally. Instead, the robust progesterone association was cross-sectional and in males, and the anticipated female longitudinal relationships were absent. These findings cannot, however, be interpreted as a total absence of progesterone influence on the cerebellar structure given the limitations of salivary quantification of the hormone.

Salivary sampling was chosen because it indexes the free, biologically active fraction of circulating hormone (typically 1-15% of the total) and enables non-invasive, repeated assessment suited to longitudinal designs. For estradiol and, in males, testosterone, salivary measures correspond well with serum (65, 66), supporting the estradiol and male testosterone findings reported here. Salivary progesterone quantifies only the small free fraction of circulating progesterone, and its correspondence with serum concentrations is more variable. Manufacturer validation reports adequate serum correspondence under optimal conditions (67), whereas a large multi-center validation study, using the same immunoassay, found poor correlation with serum (87), which weakened further at low concentrations and in conditions characteristic of the menopausal transition and postmenopausal stage (88). These conditions are represented in the current sample of females as well, and they may have attenuated the sensitivity to progesterone-volume relationships generally. The longitudinal null observed in females therefore may reflect the limits of the measured analyte in this population instead of an absence of the effect of progesterone on the cerebellar structure altogether. Salivary progesterone is also not recommended for males owing to characteristically low levels (67), warranting caution in interpreting the male progesterone-Crus II cross-sectional association.

This warrants a consideration of exploring a molecule which carries the central effects of progesterone in the brain. Progesterone is largely a precursor to allopregnanolone (ALLO), the neuroactive metabolite through which many of its actions in the brain are exerted, and which acts as a positive modulator of gamma-aminobutyric type A receptors with established neuroprotective effects (89). Salivary progesterone does not measure serum progesterone concentrations or ALLO, so a study measuring salivary progesterone may be less likely to detect cerebellar relationships that may in fact be mediated by the metabolite rather than the free fraction of the precursor in saliva. Future work using serum progesterone and ALLO could test whether any cerebellar relationships emerge when the neuroactive component of the hormone is measured directly.

### Sex differences in hormone-cerebellum relationships

Sex interactions were significant for two of the three hormones in multiple cerebellar subregions (Table 3). For estradiol, the change-score interaction spanned bilateral Crus I and Right Crus II, and extended to bilateral Lobule VI. For testosterone, the sex interaction was significant for change in Right Lobule V volume. Additionally, the estradiol interaction reflected a difference in magnitude and direction such that estradiol change in males was positively associated with change in Crus I volume, but negatively associated in females.

Moreover, sex differences in cortical hormone effects are typically framed in terms of estrogen’s neuroprotective actions which are presumed to operate in the same direction across sexes and possibly to a greater effect in females (77). However, an opposing relationship, as observed in the present study, is difficult to accommodate within a model in which sex modulates the strength of a shared hormonal mechanism. This potentially indicates that the mechanisms themselves may be sex-differentiated. The aging cerebellum is thus a structure whose endocrine regulation may be organized differently by sex, and this is a distinction that cortically derived and sex-aggregated models alone may not predict adequately.

Prior analyses of the same cohort examined sex and age effects on cerebellar volume directly, in which no sex-by-age interaction emerged, and sex differences were confined to anterior regions (49). Consistent with this regional pattern, longitudinal work has reported sex differences in cerebellar aging concentrated in specific lobules, including Lobule VI and Crus II (48), the same regions where the present hormone × sex interactions emerged. The sex-differentiation identified here shows how sex-steroid hormones impact cerebellar volume differentially between males and females. Population-level structural comparisons may show no sex differences while the underlying hormone-structure relationships differ by sex and this may underscore the value of individual variability in hormonal profiles when characterizing brain aging (77).

Furthermore, the negative association between testosterone change and Left Crus I volume change was significant in females and non-significant in males. However, the sex interaction did not reach significance, indicating that the pattern is specific to females and cannot be claimed as a formal difference between sexes. Similarly, the cross-sectional relationship between progesterone and Crus II volume was specific to males without a significant interaction. These effects are present in one sex and not reliably detected in the other, and they are reported as such. Formal interactions were regionally specific rather than spread across the cerebellum where they emerged. The effects were concentrated in Crus I and Lobule VI for estradiol, and in Lobule V for testosterone, consistent with the regional specificity evident throughout the study (Figure 4). This suggests that sex-differentiated hormonal regulation of cerebellar structure may itself be organized along functionally specific regions within the cerebellum.

**Figure 4.**
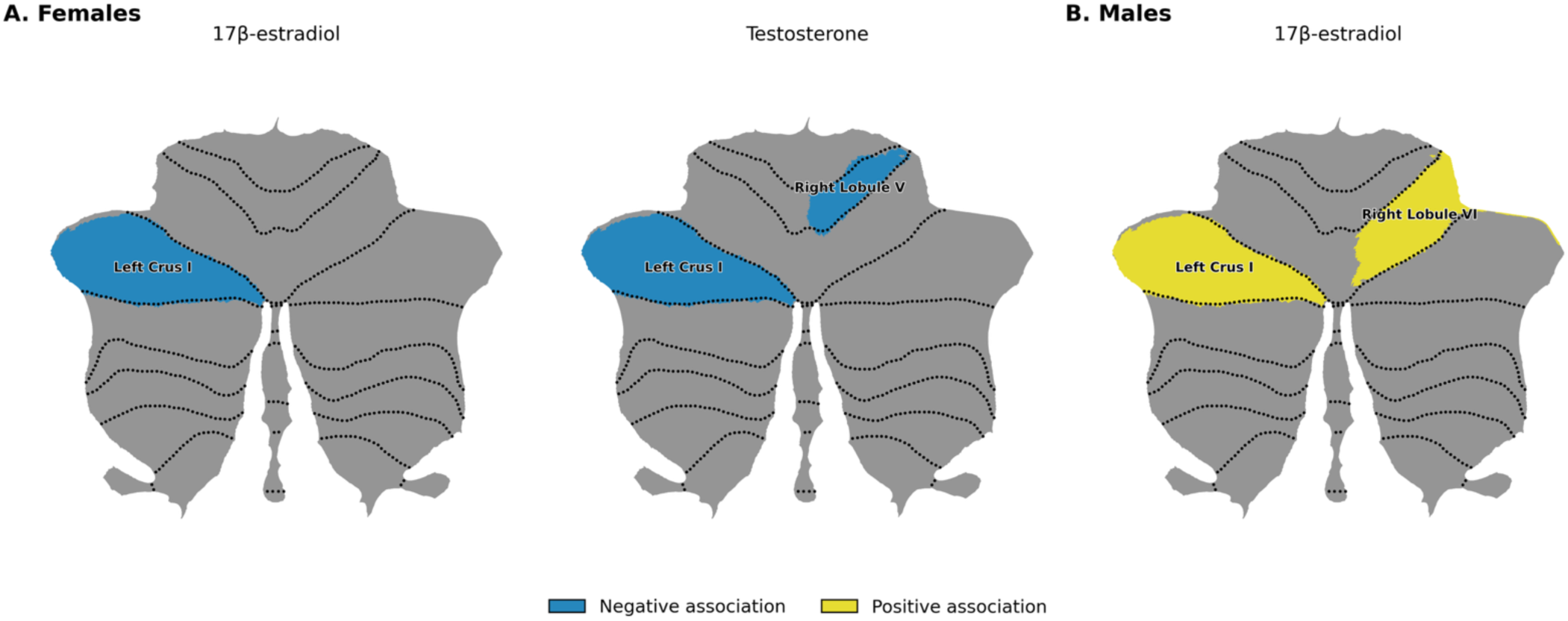
Change-Score Hormone-Cerebellar Volume Associations by Sex *Note.* Shaded regions indicate significant hormone-volume change-score associations; color denotes association direction (blue = negative, yellow = positive). (A) In females, greater estradiol change was negatively associated with change in Left Crus I volume, and greater testosterone change was negatively associated with change in Left Crus I and Right Lobule V volume. (B) In males, greater estradiol change was positively associated with change in Left Crus I and Right Lobule VI volume; male testosterone showed no significant change-score associations and is not illustrated. Primary regions (Crus I) were evaluated uncorrected; secondary regions (Lobules V, VI) were FDR-corrected. Regions are displayed on the SUIT flatmap (90), using the probabilistic lobular atlas (91); segmentation used CerebNet (see Methods).

### Cognitive relevance of hormone-cerebellum relationships

The hormone-cerebellum relationships identified here were, in post-hoc analyses moderated by cognitive status, and these moderations concentrated in Lobule V (35, 51). Baseline estradiol interacted with global cognition to predict Year 1 Lobule V volume, such that the estradiol-Lobule V relationship became more positive with higher MoCA scores in both the full sample and males. Testosterone operated on a different temporal axis but converged on the same region as estradiol. Interaction between change in testosterone levels and global cognition was associated with Lobule V volume change in females, with the negative testosterone-Lobule V relationship more pronounced at higher MoCA scores. These cognition-dependent associations of Lobule V with both estradiol and testosterone, that appeared in both sexes, suggests that this region may be a site at which hormonal and cognitive influences on the cerebellar structure converge, rather than one at which hormone and cognition act independently. A parallel estradiol-by-cognition association with global gray matter and cortical volume, in the opposite direction, indicates that this moderation is likely not confined to the cerebellum. Given that these analyses were post-hoc and are reported only as observations that may be tested directly, the results do suggest that the cognitive consequences of hormone-cerebellum associations warrant further investigation.

### Exploratory and reproductive-stage analyses

Exploratory analyses including regions beyond the primary cerebellar regions were consistent with the study’s broader pattern of regionally specific, sex-dependent hormone associations. In females, progesterone was negatively associated with Vermis X volume at Year 1 and with Lobule V volume in early postmenopausal females, while estradiol was positively associated with cerebral white matter volume at baseline in females. Notably, the progesterone associations concentrated in the vermis and Lobule V, midline, and anterior regions distinct from the Crus subregions implicated for estradiol and testosterone in the primary analyses, suggesting that progesterone may relate to a different cerebellar territory than the other two hormones. This regional segregation is consistent with evidence from animal studies that progesterone acts locally within the vermis to shape Purkinje cell dendritic development, an effect demonstrated by direct progesterone administration and blocked by progesterone-receptor antagonism in the posterior vermis (43). Whether the concentration of progesterone associations in vermal and anterior regions reflects a corresponding gradient in progesterone receptor density across cerebellar subregions remains to be established. Nonetheless, this regional pattern observed in the present study is consistent with evidence that sex differences in cerebellar aging are themselves lobule-specific (48), and that exploring hormone-specific effects on these patterns is warranted. However, these analyses were underpowered, and reproductive-stage stratification in particular produced subgroups too small for adequate interpretation of the results. They are therefore not interpreted further here and are reported in the Supplementary Materials (see Tables S16-S19) to potentially serve as a foundation for hypothesis generation in larger cohorts.

Finally, the present study has several limitations. The study included two timepoints, which supports the estimation of change but not the impact of declining or accelerating hormonal trajectories, and the interval between assessments varied across participants owing to pandemic-related scheduling. Sensitivity analyses incorporating this interval as a covariate confirmed that the change-score findings were not attributable to variations in either the interval between scans or saliva collection (see Supplementary Materials; Tables S12, S13).

Sex hormones were quantified from saliva, which captures the free rather than the total circulating fraction. As discussed above, this particularly limits interpretation of the progesterone findings. Furthermore, the male subsample was smaller than the female, and sex-stratified and reproductive-stage subgroups were in some cases small, constraining power for stratified and stage-specific analyses. Despite these limitations, the present study provides among the first longitudinal evidence on sex-steroid-cerebellum relationships across both sexes, a question that remains largely unexamined.

Beyond the hormone-specific directions noted above, several broader avenues follow from these findings. The opposite-direction estradiol relationships in males and females call for direct measurement of the mechanisms proposed to underlie them, including brain aromatase activity and its change with age, which would test whether the male cerebellar association is indeed carried by locally synthesized estradiol. Additionally, the balance between estradiol and testosterone may be a more informative index of hormonal influence on neural structure. The testosterone-to-estradiol ratio differs by sex in cerebellar tissue (92) and has been linked to cognitive performance in postmenopausal females (93). Given the cerebellum’s involvement in cognition, examining this ratio rather than each hormone in isolation, may therefore clarify sex-specific hormonal influences on cerebellar structure. The concentration of hormone-cognition moderations in Lobule V invites targeted examination of whether hormone-related cerebellar structure mediates cognitive change, a question that present post-hoc analyses can only raise.

Relatedly, whether the sex-differentiated cerebellar-cognitive relationships identified here bear on the sex differences in risk for neurodegenerative disease that motivated this work remains an open question. The cerebellum’s involvement in the progression of AD (37, 57), together with the disproportionate prevalence of AD in women (9, 58), makes the sex-specific hormonal impact on the cognitive cerebellum a candidate pathway worth investigating directly, though the present normative sample and screening-level cognitive measures cannot address it.

Here, the sex-differentiated and regionally specific character of these relationships indicates that future work on the influence of sex-steroid hormones on brain aging may benefit from subregional measures instead of treating the cerebellum as a structure without regional specificity. Additionally, analytic approaches that utilize hormone-structure relationships within each sex rather than only using sex as a covariate may be beneficial.

## Conclusion

The findings in the present study provide insight into the impact of sex-steroid hormones on the cerebellar structure in aging in a sex-differentiated, regionally specific, and directionally complex manner. As important as models derived exclusively from cortical and female-focused work are, understanding the unique effects of hormones on the cerebellum is equally important. Taken together, the opposite directions observed in the estradiol relationships in males and females, the testosterone associations specific to females, and the male cross-sectional progesterone effect in the present study indicate that the aging cerebellum is also sensitive to endocrine function and the effects differ by sex. Several of the relationships observed in this study were opposite to the direction predicted by the existing neurotrophic framing, particularly the negative association between estradiol and cerebellar volume in females and the preservation effect of estradiol observed in males, which positions them as potential empirical realities that need to be accounted for in future investigations. Our findings highlight the need for recognizing the cerebellum as a site of sex-specific hormonal influence on brain aging and identifying the mechanisms that may drive the sex-dependent effects in these relationships. The findings represent potential future directions to enhance the understanding of how sex-steroid hormones could shape the aging brain.

## Acknowledgements

This work was supported by the National Institute on Aging of the National Institutes of Health under award number R01AG064010 to J.A.B. The content is solely the responsibility of the authors and does not necessarily represent the official views of the National Institutes of Health.

## Notes

### Competing Interest Statement

The authors have declared no competing interest.

## References

1. S. M. Resnick, D. L. Pham, M. A. Kraut, A. B. Zonderman, C. Davatzikos, Longitudinal Magnetic Resonance Imaging Studies of Older Adults: A Shrinking Brain. J. Neurosci. 23, 3295–3301 (2003).

2. I. Skampardoni, et al., Coupled cross-sectional and longitudinal non-negative matrix factorization reveals dominant brain aging trajectories in 48,949 individuals. Nat. Commun. 17, 5709 (2026).

3. S. J. Ritchie, et al., Sex Differences in the Adult Human Brain: Evidence from 5216 UK Biobank Participants. Cereb. Cortex 28, 2959–2975 (2018).

4. S. N. Vucic, B. Georges, S. Frangou, N. Sadeghi, T. White, Sex differences in global metrics of brain size across the lifespan. Front. Neurosci. 20, 1646144 (2026).

5. M. Lotze, et al., Novel findings from 2,838 Adult Brains on Sex Differences in Gray Matter Brain Volume. Sci. Rep. 9, 1671 (2019).

6. N. M. Armstrong, et al., Sex differences in brain aging and predictors of neurodegeneration in cognitively healthy older adults. Neurobiol. Aging 81, 146–156 (2019).

7. A. Ravndal, et al., Sex differences in healthy brain aging are unlikely to explain higher Alzheimer’s disease prevalence in women. Proc. Natl. Acad. Sci. 122, e2510486122 (2025).

8. F. Crivello, N. Tzourio-Mazoyer, C. Tzourio, B. Mazoyer, Longitudinal Assessment of Global and Regional Rate of Grey Matter Atrophy in 1,172 Healthy Older Adults: Modulation by Sex and Age. PLoS ONE 9, e114478 (2014).

9. 2026 Alzheimer’s disease facts and figures. Alzheimers Dement. 22, e71345 (2026).

10. R. F. Buckley, et al., Menopause Status Moderates Sex Differences in Tau Burden: A Framingham PET Study. Ann. Neurol. 92, 11–22 (2022).

11. S. Vadakkadath Meethal, C. S. Atwood, Alzheimer’s disease: the impact of age-related changes in reproductive hormones: The role of hypothalamic-pituitary-gonadal hormones in the normal structure and functioning of the brain. CMLS Cell. Mol. Life Sci. 62, 257–270 (2005).

12. S. Jett, et al., Endogenous and Exogenous Estrogen Exposures: How Women’s Reproductive Health Can Drive Brain Aging and Inform Alzheimer’s Prevention. Front. Aging Neurosci. 14, 831807 (2022).

13. M. M. Khan, K. M. Dhandapani, Q. Zhang, D. W. Brann, Estrogen regulation of spine density and excitatory synapses in rat prefrontal and somatosensory cerebral cortex. Steroids 78, 614–623 (2013).

14. Y. Zhu, et al., Protective Effect of 17β-Estradiol Upon Hippocampal Spine Density and Cognitive Function in an Animal Model of Vascular Dementia. Sci. Rep. 7, 42660 (2017).

15. J. Hao, et al., Estrogen Alters Spine Number and Morphology in Prefrontal Cortex of Aged Female Rhesus Monkeys. J. Neurosci. 26, 2571–2578 (2006).

16. L. Mosconi, et al., Menopause impacts human brain structure, connectivity, energy metabolism, and amyloid-beta deposition. Sci. Rep. 11, 10867 (2021).

17. R. G. Zsido, et al., Ultra-high-field 7T MRI reveals changes in human medial temporal lobe volume in female adults during menstrual cycle. Nat. Ment. Health 1, 761–771 (2023).

18. G.-W. Kim, K. Park, Y.-H. Kim, G.-W. Jeong, Altered brain morphology and functional connectivity in postmenopausal women: automatic segmentation of whole-brain and thalamic subnuclei and resting-state fMRI. Aging 16, 4965–4979 (2024).

19. K. I. Erickson, M. W. Voss, R. S. Prakash, L. Chaddock, A. F. Kramer, A cross-sectional study of hormone treatment and hippocampal volume in postmenopausal women: Evidence for a limited window of opportunity. Neuropsychology 24, 68–76 (2010).

20. A. Ambikairajah, et al., Investigating the synergistic effects of hormone replacement therapy, APOLIPOPROTEIN E and age on brain health in the UK Biobank. Hum. Brain Mapp. 45, e26612 (2024).

21. C. M. Taylor, et al., Progesterone shapes medial temporal lobe volume across the human menstrual cycle. NeuroImage 220, 117125 (2020).

22. C. Heller, et al., Hormonal milieu influences whole-brain structural dynamics across the menstrual cycle using dense sampling in multiple individuals. Nat. Neurosci. 28, 2588– 2600 (2025).

23. S. Nehls, E. Losse, M. Armin, U. Habel, N. Chechko, Progesterone-associated adjustments in brain structure during menstruation and the periovulatory phase—an MRI study. eBioMedicine 125, 106184 (2026).

24. L. Pritschet, et al., Neuroanatomical changes observed over the course of a human pregnancy. Nat. Neurosci. 27, 2253–2260 (2024).

25. Y. Niu, et al., Longitudinal investigation of neurobiological changes across pregnancy. *Commun*. Biol. 8, 82 (2025).

26. C. Roselli, M. Liu, P. Hurn, Brain Aromatization: Classic Roles and New Perspectives. Semin. Reprod. Med. 27, 207–217 (2009).

27. A. Biegon, et al., Aromatase Imaging with [ *N* -Methyl-^11^ C]Vorozole PET in Healthy Men and Women. J. Nucl. Med. 56, 580–585 (2015).

28. B. Pletzer, Sex Hormones and Gender Role Relate to Gray Matter Volumes in Sexually Dimorphic Brain Areas. Front. Neurosci. 13, 592 (2019).

29. M. Nerattini, et al., Sex-specific associations of serum testosterone with gray matter volume and cerebral blood flow in midlife individuals at risk for Alzheimer’s disease. PLOS ONE 20, e0317303 (2025).

30. C. Barth, A. Crestol, A.-M. G. De Lange, L. A. M. Galea, Sex steroids and the female brain across the lifespan: insights into risk of depression and Alzheimer’s disease. Lancet Diabetes Endocrinol. 11, 926–941 (2023).

31. A. G. De Lange, et al., Women’s brain aging: Effects of sex-hormone exposure, pregnancies, and genetic risk for Alzheimer’s disease. Hum. Brain Mapp. 41, 5141–5150 (2020).

32. A. Arleo, et al., Consensus Paper: Cerebellum and Ageing. The Cerebellum 23, 802–832 (2023).

33. R. L. Buckner, The Cerebellum and Cognitive Function: 25 Years of Insight from Anatomy and Neuroimaging. Neuron 80, 807–815 (2013).

34. J. D. Schmahmann, J. C. Sherman, The cerebellar cognitive affective syndrome. (1998).

35. C. J. Stoodley, E. M. Valera, J. D. Schmahmann, Functional topography of the cerebellum for motor and cognitive tasks: An fMRI study. NeuroImage 59, 1560–1570 (2012).

36. E. Hoxha, et al., The Emerging Role of Altered Cerebellar Synaptic Processing in Alzheimer’s Disease. Front. Aging Neurosci. 10, 396 (2018).

37. S. Toniolo, et al., Patterns of Cerebellar Gray Matter Atrophy Across Alzheimer’s Disease Progression. Front. Cell. Neurosci. 12, 430 (2018).

38. S. J. Colloby, J. T. O’Brien, J.-P. Taylor, Patterns of cerebellar volume loss in dementia with Lewy bodies and Alzheimer’s disease: A VBM-DARTEL study. Psychiatry Res. Neuroimaging 223, 187–191 (2014).

39. R. H. Price Jr., R. J. Handa, Expression of estrogen receptor-beta protein and mRNA in the cerebellum of the rat. Neurosci. Lett. 288, 115–118 (2000).

40. C. Guerra-Araiza, A. Coyoy-Salgado, I. Camacho-Arroyo, Sex differences in the regulation of progesterone receptor isoforms expression in the rat brain. Brain Res. Bull. 59, 105–109 (2002).

41. M. Perez-Pouchoulen, et al., Androgen receptors in Purkinje neurons are modulated by systemic testosterone and sexual training in a region-specific manner in the male rat. Physiol. Behav. 156, 191–198 (2016).

42. K. Tsutsui, S. Haraguchi, Neuroprotective actions of cerebellar and pineal allopregnanolone on Purkinje cells. FASEB BioAdvances 2, 149–159 (2020).

43. H. Sakamoto, K. Ukena, K. Tsutsui, Effects of Progesterone Synthesized *De Novo* in the Developing Purkinje Cell on Its Dendritic Growth and Synaptogenesis. J. Neurosci. 21, 6221–6232 (2001).

44. H. Sakamoto, Y. Mezaki, H. Shikimi, K. Ukena, K. Tsutsui, Dendritic Growth and Spine Formation in Response to Estrogen in the Developing Purkinje Cell. Endocrinology 144, 4466–4477 (2003).

45. J. M. Abel, D. M. Witt, E. F. Rissman, Sex Differences in the Cerebellum and Frontal Cortex: Roles of Estrogen Receptor Alpha and Sex Chromosome Genes. Neuroendocrinology 93, 230–240 (2011).

46. N. Raz, F. Gunning-Dixon, D. Head, A. Williamson, J. D. Acker, Age and Sex Differences in the Cerebellum and the Ventral Pons: A Prospective MR Study of Healthy Adults. AJNR Am. J. Neuroradiol. 22, 1161–1167 (2001).

47. T. H. Hicks, H. K. Ballard, H. Sang, J. A. Bernard, Age–volume associations in cerebellar lobules by sex and reproductive stage. Brain Struct. Funct. 227, 2439–2455 (2022).

48. S. Han, Y. An, A. Carass, J. L. Prince, S. M. Resnick, Longitudinal analysis of regional cerebellum volumes during normal aging. NeuroImage 220, 117062 (2020).

49. T. N. C. Magalhães, et al., The Influence of Sex and Sex-Steroid Hormones on Cerebellar Structure and Functional Connectivity Across Adulthood. Hum. Brain Mapp. 46, e70356 (2025).

50. H. K. Ballard, T. B. Jackson, T. H. Hicks, J. A. Bernard, The association of reproductive stage with lobular cerebellar network connectivity across female adulthood. Neurobiol. Aging 117, 139–150 (2022).

51. C. J. Stoodley, J. Schmahmann, Functional topography in the human cerebellum: A meta-analysis of neuroimaging studies. NeuroImage 44, 489–501 (2009).

52. T. H. Hicks, et al., Network segregation in aging females and evaluation of the impact of sex steroid hormones. Front. Hum. Neurosci. 17, 1059091 (2023).

53. J. A. Bernard, R. D. Seidler, Relationships Between Regional Cerebellar Volume and Sensorimotor and Cognitive Function in Young and Older Adults. The Cerebellum 12, 721–737 (2013).

54. J. A. Bernard, Don’t forget the little brain: A framework for incorporating the cerebellum into the understanding of cognitive aging. Neurosci. Biobehav. Rev. 137, 104639 (2022).

55. F. d’Oleire Uquillas, et al., Cerebellar aging is spatially heterogeneous and supports cognitive resilience in later life. Nat. Neurosci. (2026). 10.1038/s41593-026-02289-x.

56. C. P. Cooper, et al., Associations of baseline and longitudinal change in cerebellum volume with age-related changes in verbal learning and memory. NeuroImage 272, 120048 (2023).

57. Y. H. Um, et al., Longitudinal analysis of cerebellar volume changes across the Alzheimer’s disease spectrum: implications for cognitive function and neuropsychiatric symptoms. Alzheimers Dement. 21, e099275 (2025).

58. L. Castro-Aldrete, et al., Alzheimer disease seen through the lens of sex and gender. Nat. Rev. Neurol. 21, 235–249 (2025).

59. C. Jockwitz, C. Krämer, P. Dellani, S. Caspers, Differential predictability of cognitive profiles from brain structure in older males and females. GeroScience 46, 1713–1730 (2023).

60. M. A. Dratva, et al., Low testosterone levels relate to poorer cognitive function in women in an APOE-ε4-dependant manner. Biol. Sex Differ. 15, 45 (2024).

61. M. F. R. Sowers, et al., Testosterone, sex hormone-binding globulin and free androgen index among adult women: chronological and ovarian aging. Hum. Reprod. 24, 2276–2285 (2009).

62. B. G. Keevil, et al., Distribution of Salivary Testosterone in Men and Women in a British General Population-Based Sample: The Third National Survey of Sexual Attitudes and Lifestyles (Natsal-3). J. Endocr. Soc. 1, 14–25 (2017).

63. T. H. Hicks, et al., Functional and structural cerebellar-behavior relationships in aging. Brain Struct. Funct. 230, 10 (2024).

64. S. D. Harlow, et al., Executive summary of the Stages of Reproductive Aging Workshop + 10: addressing the unfinished agenda of staging reproductive aging. Menopause 19, 387–395 (2012).

65. Salimetrics SalivaLab, Salimetrics Salivary Estradiol Assay Kit (Cat. No. 1-3702). (2022). Available at: https://salimetrics.com/analyte/salivary-estradiol/ [Accessed 24 July 2026].

66. Salimetrics SalivaLab, Salimetrics Salivary Testosterone Assay Kit (Cat. No. 1-2402). (2022). Available at: https://salimetrics.com/analyte/salivary-testosterone/ [Accessed 24 July 2026].

67. Salimetrics SalivaLab, Salimetrics Salivary Progesterone Assay Kit (Cat. No. 1-1502). (2022). Available at: https://salimetrics.com/analyte/salivary-progesterone/ [Accessed 24 July 2026].

68. R. W. Hornung, L. D. Reed, Estimation of Average Concentration in the Presence of Nondetectable Values. Appl. Occup. Environ. Hyg. 5, 46–51 (1990).

69. M. P. Harms, et al., Extending the Human Connectome Project across ages: Imaging protocols for the Lifespan Development and Aging projects. NeuroImage 183, 972–984 (2018).

70. L. Henschel, et al., FastSurfer - A fast and accurate deep learning based neuroimaging pipeline. NeuroImage 219, 117012 (2020).

71. L. Henschel, D. Kügler, M. Reuter, FastSurferVINN: Building resolution-independence into deep learning segmentation methods—A solution for HighRes brain MRI. NeuroImage 251, 118933 (2022).

72. J. Faber, et al., CerebNet: A fast and reliable deep-learning pipeline for detailed cerebellum sub-segmentation. NeuroImage 264, 119703 (2022).

73. Z. S. Nasreddine, et al., The Montreal Cognitive Assessment, MoCA: A Brief Screening Tool For Mild Cognitive Impairment. J. Am. Geriatr. Soc. 53, 695–699 (2005).

74. D. Wechsler, Wechsler Adult Intelligence Scale – Fourth Edition: Administration and scoring manual, 4th Ed. (Pearson Assessment, 2008).

75. N. Benson, D. M. Hulac, J. H. Kranzler, Independent examination of the Wechsler Adult Intelligence Scale—Fourth Edition (WAIS-IV): What does the WAIS-IV measure? Psychol. Assess. 22, 121–130 (2010).

76. Y. Benjamini, Y. Hochberg, Controlling the False Discovery Rate: A Practical and Powerful Approach to Multiple Testing. J. R. Stat. Soc. Ser. B Stat. Methodol. 57, 289–300 (1995).

77. G. E. Gillies, S. McArthur, Estrogen Actions in the Brain and the Basis for Differential Action in Men and Women: A Case for Sex-Specific Medicines. Pharmacol. Rev. 62, 155– 198 (2010).

78. V. L. Hedges, T. J. Ebner, R. L. Meisel, P. G. Mermelstein, The cerebellum as a target for estrogen action. Front. Neuroendocrinol. 33, 403–411 (2012).

79. S. Bhasin, Testosterone replacement in aging men: an evidence-based patient-centric perspective. J. Clin. Invest. 131, e146607 (2021).

80. J. P. Dias, et al., Effects of aromatase inhibition vs. testosterone in older men with low testosterone: randomized-controlled trial. Andrology 4, 33–40 (2016).

81. S. D. Moffat, Effects of Testosterone on Cognitive and Brain Aging in Elderly Men. Ann. N. Y. Acad. Sci. 1055, 80–92 (2005).

82. L. L. Gravelsins, L. A. M. Galea, One size does not fit all: how type of menopause and hormone therapy matters for brain health. Br. J. Psychiatry 226, 369–382 (2025).

83. H. J. Teede, et al., Polyendocrine metabolic ovarian syndrome, the new name for polycystic ovary syndrome: a multistep global consensus process. The Lancet 407, 2329–2339 (2026).

84. B. Ozgen Saydam, B. O. Yildiz, Polycystic Ovary Syndrome and Brain: An Update on Structural and Functional Studies. J. Clin. Endocrinol. Metab. 106, e430–e441 (2021).

85. M.-J. Chen, et al., Influence of dehydroepiandrosterone sulphate levels on the slower age-related decline in grey matter in younger women with polycystic ovary syndrome. Brain Commun. 7, fcaf052 (2024).

86. N. Baspinar, S. Salk, A. Ersoz, Automated MRI-based brain volumetric analysis in women with polycystic ovary syndrome. BMC Med. Imaging 26, 262 (2026).

87. D. Sakkas, et al., A multi-centre international study of salivary hormone oestradiol and progesterone measurements in ART monitoring. Reprod. Biomed. Online 42, 421–428 (2021).

88. E. Ferrer, et al., The use of saliva and blood progesterone to profile the menstrual cycles of youth professional football players. Front. Sports Act. Living 6, 1430158 (2024).

89. R. Guennoun, Progesterone in the Brain: Hormone, Neurosteroid and Neuroprotectant. Int. J. Mol. Sci. 21, 5271 (2020).

90. J. Diedrichsen, E. Zotow, Surface-Based Display of Volume-Averaged Cerebellar Imaging Data. PLOS ONE 10, e0133402 (2015).

91. J. Diedrichsen, J. H. Balsters, J. Flavell, E. Cussans, N. Ramnani, A probabilistic MR atlas of the human cerebellum. (2009).

92. D. Caruso, et al., Comparison of plasma and cerebrospinal fluid levels of neuroactive steroids with their brain, spinal cord and peripheral nerve levels in male and female rats. Psychoneuroendocrinology 38, 2278–2290 (2013).

93. J. Ryan, et al., Hormone levels and cognitive function in postmenopausal midlife women. Neurobiol. Aging 33, 1138–1147.e2 (2012).

