## Supplementary Information for "Longitudinal Impacts of Sex-Steroid Hormones on Cerebellar Structure in Adulthood are Sex Differentiated"

**Affiliations:**

**This PDF file includes:**

Figures S1 to S3

Tables S1 to S19

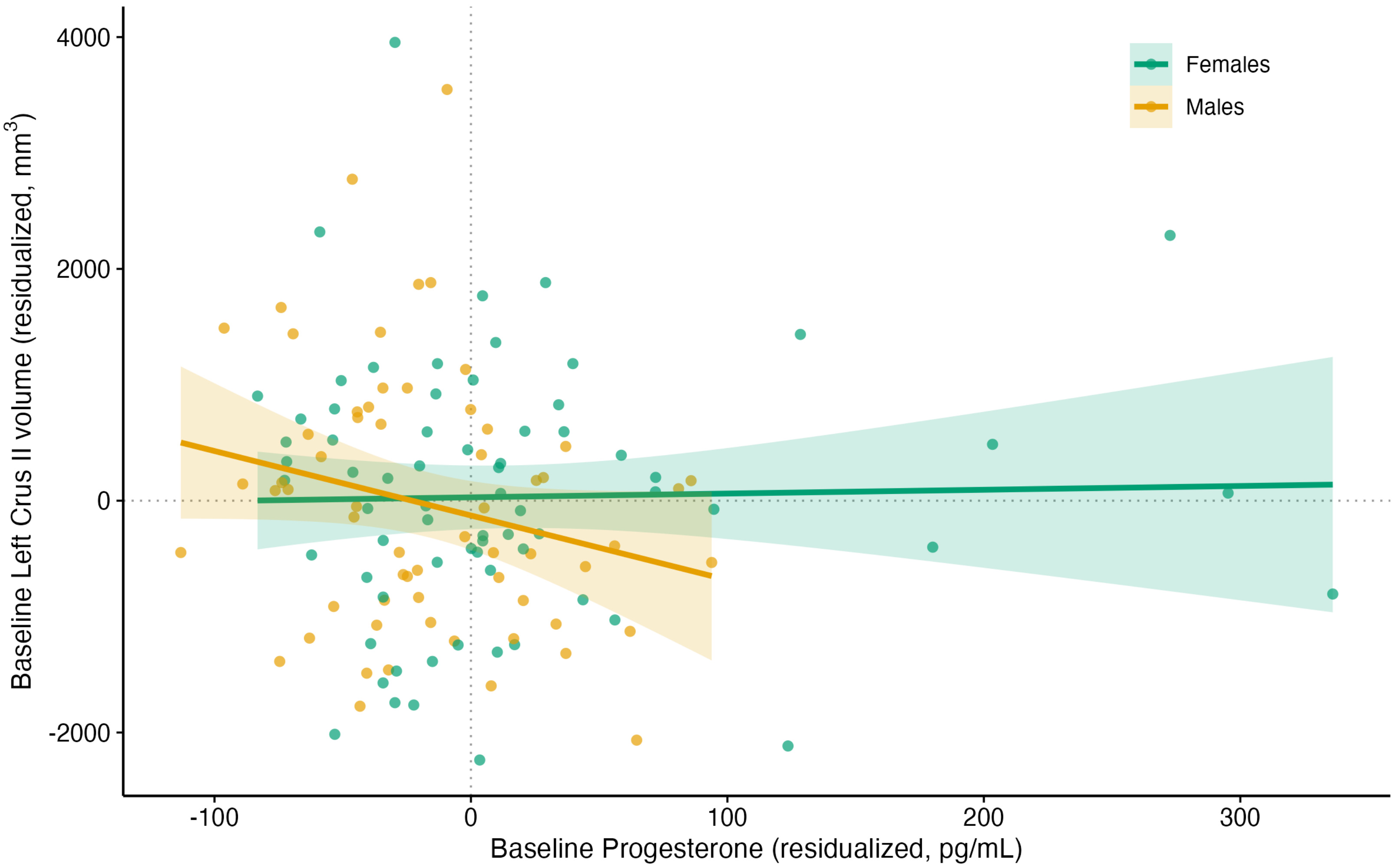

**Figure S1.** Progesterone and Left Crus II volume by sex at baseline.

*Note.* Residualized cross-sectional association between baseline progesterone and baseline Left Crus II volume, shown for both sexes (outliers retained). The association was significant in males ($n$ = 62; $b$ = -7.68, $p$ = .025) and non-significant in females ($n$ = 70; $b$ = -0.01, $p$ = .996); the female slope is shown for contrast, illustrating the male-specific pattern underlying the significant progesterone × sex interaction. Points are individual participants; shaded bands are 95% confidence intervals. Both axes are residualized for age, education, and estimated total intracranial volume.

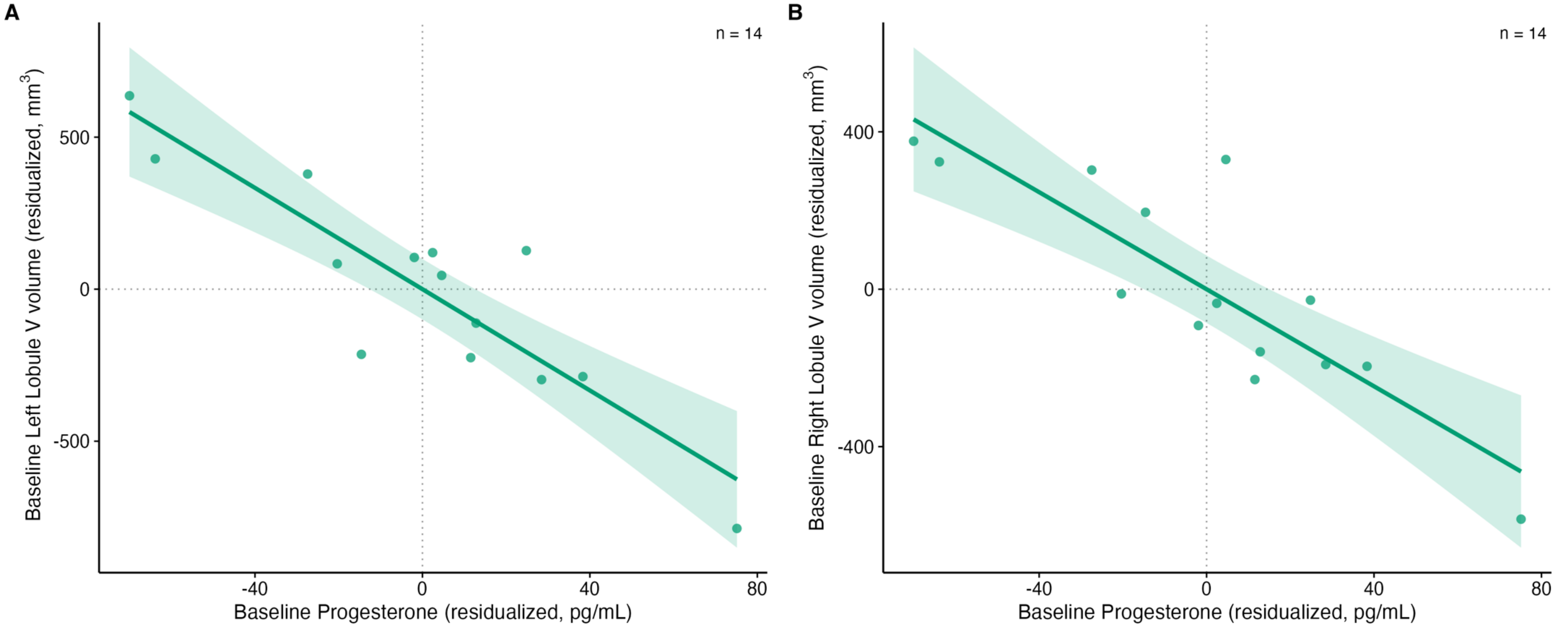

**Figure S2.** Progesterone and bilateral Lobule V volume in early-postmenopausal females at baseline.

*Note.* Residualized cross-sectional associations between baseline progesterone and baseline (A) Left and (B) Right Lobule V volume in early-postmenopausal females ($n$ = 14). Greater progesterone levels were associated with smaller Lobule V volumes bilaterally (Left $b$ = -8.33, $p$_FDR < .001; Right $b$ = -6.17, $p$_FDR = .001). Points are individual participants; shaded bands are 95% confidence intervals. Both axes are residualized for age, education, and estimated total intracranial volume. Given the small subgroup, these findings are exploratory and interpreted as preliminary.

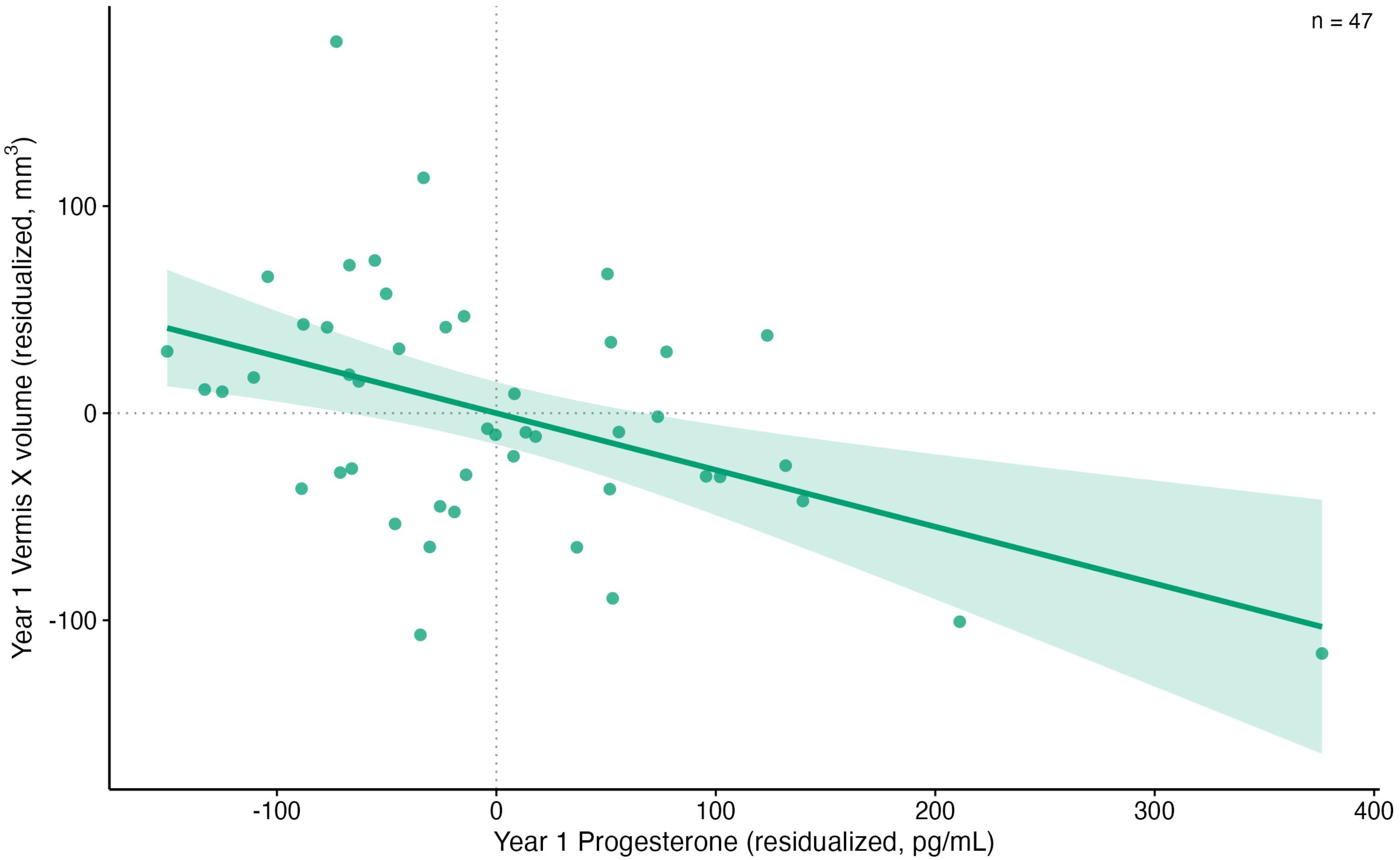

**Figure S3.** Progesterone and Vermis X volume in females at Year 1.

*Note.* Residualized cross-sectional association between Year 1 progesterone and Year 1 Vermis X volume in females (*n* = 47; outliers retained). Greater progesterone was associated with smaller Vermis X volume (*b* = -0.27, *p*_FDR = .016). Points are individual participants; shaded bands are 95% confidence intervals. Both axes are residualized for age, education, and estimated total intracranial volume.

**Table S1.** Hormone levels and raw cerebellar volumes by sex and timepoint.

| | Baseline | | Year 1 | |
|---|---|---|---|---|
| | Females | Males | Females | Males |
| **Hormones (pg/mL), *M* ± *SD*** | | | | |
| Estradiol | 1.3 ± 0.9 | 1.0 ± 0.6 | 1.3 ± 0.8 | 1.3 ± 0.8 |
| Testosterone | 75.3 ± 36.2 | 154.3 ± 69.8 | 78.4 ± 34.0 | 159.4 ± 47.2 |
| Progesterone | 84.9 ± 94.7 | 59.6 ± 46.8 | 106.8 ± 106.8 | 98.5 ± 64.4 |
| **Primary ROI volume (mm³), *M* ± *SD*** | | | | |
| Right Crus I | 10908 ± 1559 | 11734 ± 1873 | 10926 ± 1616 | 11971 ± 1764 |
| Left Crus I | 10542 ± 1392 | 11301 ± 1799 | 10549 ± 1368 | 11515 ± 1702 |
| Right Crus II | 6866 ± 1203 | 7531 ± 1236 | 6980 ± 1166 | 7610 ± 1261 |
| Left Crus II | 6727 ± 1158 | 6898 ± 1134 | 6713 ± 1160 | 6887 ± 1244 |
| **Secondary ROI volume (mm³), *M* ± *SD*** | | | | |
| Right Lobule V | 2744 ± 383 | 3160 ± 422 | 2738 ± 416 | 3237 ± 422 |
| Left Lobule V | 2863 ± 441 | 3230 ± 447 | 2884 ± 474 | 3262 ± 403 |
| Right Lobule VI | 6928 ± 963 | 7159 ± 987 | 7019 ± 975 | 7195 ± 975 |
| Left Lobule VI | 6859 ± 963 | 7095 ± 1078 | 6910 ± 1020 | 7145 ± 1079 |

*Note*. *M* = mean; *SD* = standard deviation. Hormone levels reflect free salivary levels (pg/mL); volumes are raw and uncorrected for intracranial volume (mm³).

**Table S2.** Hormone-cerebellar volume associations: Baseline, Year 1, and Baseline-predicting Year 1 models.

| | | | Baseline | | | | | | Year 1 | | | | | | Baseline→Year 1 | | | | | |
|---|---|---|---|---|---|---|---|---|---|---|---|---|---|---|---|---|---|---|---|---|
| | | | E | | T | | P | | E | | T | | P | | E | | T | | P | |
| | | **Region** | ***b* (*SE*)** | ***p*** | ***b* (*SE*)** | ***p*** | ***b* (*SE*)** | ***p*** | ***b* (*SE*)** | ***p*** | ***b* (*SE*)** | ***p*** | ***b* (*SE*)** | ***p*** | ***b* (*SE*)** | ***p*** | ***b* (*SE*)** | ***p*** | ***b* (*SE*)** | ***p*** |
| **Full Sample** | | | *n* = 136 | | *n* = 135 | | *n* = 132 | | *n* = 92 | | *n* = 91 | | *n* = 90 | | *n* = 93 | | *n* = 92 | | *n* = 91 | |
| | **Primary ROI (uncorrected)** | Right Crus I | −310.6 (176.6) | 0.081 | 1.0 (2.6) | 0.682 | **−4.5 (2.0)** | **0.023*** | **−468.0 (214.9)** | **0.032*** | −6.3 (4.4) | 0.150 | **−5.0 (2.0)** | **0.014*** | 83.8 (62.2) | 0.181 | 0.1 (0.7) | 0.896 | 0.2 (0.5) | 0.661 |
| | | Left Crus I | −218.6 (172.0) | 0.206 | 3.1 (2.5) | 0.213 | −3.4 (1.9) | 0.077 | **−463.6 (200.7)** | **0.023*** | −2.1 (4.1) | 0.607 | **−4.2 (1.9)** | **0.027*** | 3.3 (61.2) | 0.957 | −0.3 (0.7) | 0.682 | −0.8 (0.5) | 0.121 |
| | | Right Crus II | −189.3 (138.6) | 0.174 | 3.4 (2.0) | 0.084 | 0.4 (1.6) | 0.782 | −102.9 (168.7) | 0.544 | 2.5 (3.4) | 0.464 | 1.4 (1.6) | 0.383 | 43.0 (72.0) | 0.552 | −0.5 (0.8) | 0.548 | −0.6 (0.6) | 0.359 |
| | | Left Crus II | −214.0 (130.9) | 0.105 | 1.5 (1.9) | 0.441 | −1.1 (1.5) | 0.476 | −67.4 (167.3) | 0.688 | 2.6 (3.3) | 0.447 | 0.3 (1.6) | 0.829 | 84.7 (66.6) | 0.206 | −0.5 (0.7) | 0.472 | 0.2 (0.6) | 0.663 |
| | **Secondary ROI (FDR-corrected)** | Right Lobule V | −16.2 (42.6) | 0.704 | 1.0 (0.6) | 0.338 | −0.1 (0.5) | 0.816 | −64.9 (51.9) | 0.430 | −0.3 (1.0) | 0.810 | 0.0 (0.5) | 0.980 | 12.3 (23.9) | 0.610 | −0.1 (0.3) | 0.747 | −0.2 (0.2) | 0.600 |
| | | Left Lobule V | −20.1 (46.5) | 0.704 | 0.9 (0.7) | 0.338 | 0.2 (0.5) | 0.816 | 7.3 (54.1) | 0.892 | −1.6 (1.1) | 0.584 | −0.4 (0.5) | 0.675 | −11.2 (21.8) | 0.610 | −0.2 (0.2) | 0.572 | −0.2 (0.2) | 0.600 |
| | | Right Lobule VI | −147.9 (98.7) | 0.273 | −0.2 (1.4) | 0.915 | −1.9 (1.1) | 0.174 | −28.2 (121.0) | 0.892 | −0.8 (2.4) | 0.810 | −0.8 (1.2) | 0.675 | 81.5 (38.5) | 0.109 | −0.3 (0.4) | 0.572 | −0.1 (0.3) | 0.697 |
| | | Left Lobule VI | −202.0 (102.3) | 0.202 | −0.6 (1.5) | 0.885 | −2.1 (1.2) | 0.174 | −214.7 (126.2) | 0.370 | −1.9 (2.5) | 0.810 | −1.4 (1.2) | 0.675 | 58.9 (30.2) | 0.109 | 0.4 (0.3) | 0.572 | −0.1 (0.3) | 0.697 |
| **Females** | | | *n* = 72 | | *n* = 72 | | *n* = 70 | | *n* = 48 | | *n* = 48 | | *n* = 47 | | *n* = 49 | | *n* = 48 | | *n* = 47 | |
| | **Primary ROI (uncorrected)** | Right Crus I | **−468.2 (198.6)** | **0.021*** | −7.0 (4.9) | 0.158 | −3.6 (2.2) | 0.106 | **−698.7 (320.3)** | **0.035*** | −8.0 (7.2) | 0.272 | −4.1 (2.4) | 0.090 | 144.9 (83.0) | 0.088 | 2.1 (1.3) | 0.110 | 0.7 (0.6) | 0.283 |
| | | Left Crus I | −349.4 (180.5) | 0.057 | −3.3 (4.4) | 0.453 | −2.6 (2.0) | 0.189 | **−590.4 (270.8)** | **0.035*** | −1.7 (6.2) | 0.781 | −3.1 (2.0) | 0.127 | 14.5 (74.7) | 0.847 | 0.2 (1.2) | 0.836 | −0.2 (0.6) | 0.770 |
| | | Right Crus II | −183.6 (159.1) | 0.253 | 1.6 (3.9) | 0.673 | 1.2 (1.8) | 0.490 | −66.0 (241.7) | 0.786 | −0.3 (5.2) | 0.951 | 1.0 (1.8) | 0.563 | 104.6 (98.8) | 0.295 | −1.0 (1.6) | 0.508 | −0.4 (0.8) | 0.627 |
| | | Left Crus II | −187.7 (156.8) | 0.236 | −1.3 (3.9) | 0.738 | 0.0 (1.8) | 0.996 | 57.3 (246.3) | 0.817 | 1.4 (5.3) | 0.796 | −0.3 (1.8) | 0.891 | 113.8 (105.0) | 0.285 | −0.8 (1.6) | 0.645 | 0.8 (0.8) | 0.303 |

| | | | | | | | | | | | | | | | | | | | | |
|---|---|---|---|---|---|---|---|---|---|---|---|---|---|---|---|---|---|---|---|---|
| | **Secondary ROI (FDR-corrected)** | Right Lobule V | −80.0 (44.9) | 0.228 | 0.8 (1.1) | 0.697 | −0.5 (0.5) | 0.471 | −73.5 (69.7) | 0.578 | −0.2 (1.5) | 0.992 | −0.3 (0.5) | 0.591 | 32.4 (31.5) | 0.414 | 0.6 (0.5) | 0.463 | −0.1 (0.2) | 0.952 |
| | | Left Lobule V | −69.1 (49.9) | 0.228 | −0.5 (1.2) | 0.697 | −0.4 (0.6) | 0.535 | 43.7 (78.0) | 0.578 | −0.9 (1.7) | 0.992 | −0.5 (0.6) | 0.502 | 1.5 (29.8) | 0.961 | −0.3 (0.4) | 0.528 | −0.2 (0.2) | 0.952 |
| | | Right Lobule VI | −127.9 (115.3) | 0.271 | −1.8 (2.9) | 0.697 | −2.6 (1.2) | 0.082 | −108.3 (188.8) | 0.578 | 0.0 (4.1) | 0.992 | −1.6 (1.4) | 0.502 | **117.3 (43.2)** | **0.038*** | 0.5 (0.7) | 0.528 | 0.0 (0.3) | 0.952 |
| | | Left Lobule VI | −172.9 (116.7) | 0.228 | −1.1 (2.9) | 0.697 | −2.7 (1.2) | 0.082 | −314.4 (193.3) | 0.444 | −1.7 (4.3) | 0.992 | −2.7 (1.4) | 0.238 | 41.6 (40.5) | 0.414 | 0.8 (0.6) | 0.463 | 0.0 (0.3) | 0.952 |
| **Males** | | | *n* = 64 | | *n* = 63 | | *n* = 62 | | *n* = 44 | | *n* = 43 | | *n* = 43 | | *n* = 44 | | *n* = 44 | | *n* = 44 | |
| | **Primary ROI (uncorrected)** | Right Crus I | 29.9 (331.3) | 0.928 | 2.8 (2.9) | 0.343 | −5.1 (4.6) | 0.271 | −155.9 (297.0) | 0.603 | −6.5 (5.4) | 0.233 | −7.1 (3.9) | 0.075 | 56.8 (88.3) | 0.524 | −0.5 (0.8) | 0.501 | −0.4 (1.2) | 0.719 |
| | | Left Crus I | 56.2 (354.9) | 0.875 | 4.8 (3.1) | 0.129 | −5.4 (4.9) | 0.274 | −277.7 (312.8) | 0.380 | −3.1 (5.8) | 0.596 | −7.3 (4.1) | 0.086 | 21.9 (92.4) | 0.814 | −0.5 (0.8) | 0.503 | −1.4 (1.3) | 0.272 |
| | | Right Crus II | −239.8 (273.4) | 0.384 | 4.5 (2.4) | 0.060 | −5.2 (3.7) | 0.167 | −181.8 (251.5) | 0.474 | 5.4 (4.6) | 0.249 | 1.4 (3.4) | 0.686 | −28.9 (109.5) | 0.793 | −0.3 (0.9) | 0.745 | −1.1 (1.5) | 0.479 |
| | | Left Crus II | −314.4 (247.2) | 0.208 | 2.7 (2.2) | 0.219 | **−7.7 (3.3)** | **0.025*** | −196.5 (242.9) | 0.423 | 4.2 (4.5) | 0.354 | 1.1 (3.3) | 0.751 | 53.1 (72.1) | 0.466 | −0.6 (0.6) | 0.300 | −0.4 (1.0) | 0.725 |
| | **Secondary ROI (FDR-corrected)** | Right Lobule V | 110.6 (79.4) | 0.322 | 1.1 (0.7) | 0.283 | 1.1 (1.1) | 0.709 | −12.7 (70.6) | 0.858 | 0.0 (1.3) | 0.972 | 0.1 (1.0) | 0.910 | −11.7 (37.7) | 0.758 | −0.4 (0.3) | 0.502 | −0.9 (0.5) | 0.330 |
| | | Left Lobule V | 68.4 (90.0) | 0.450 | 1.5 (0.8) | 0.212 | 1.0 (1.2) | 0.709 | −14.0 (72.6) | 0.858 | −1.4 (1.3) | 0.842 | −0.9 (1.0) | 0.748 | −23.7 (35.4) | 0.687 | −0.1 (0.3) | 0.662 | −0.4 (0.5) | 0.621 |
| | | Right Lobule VI | −226.4 (191.2) | 0.322 | 0.5 (1.7) | 0.765 | −1.7 (2.7) | 0.709 | 69.6 (158.2) | 0.858 | −1.3 (2.9) | 0.883 | 1.2 (2.1) | 0.765 | 43.7 (66.5) | 0.687 | −0.8 (0.6) | 0.502 | 0.4 (0.9) | 0.684 |
| | | Left Lobule VI | −304.3 (201.9) | 0.322 | −0.6 (1.8) | 0.765 | −0.8 (2.9) | 0.781 | −73.7 (170.0) | 0.858 | −2.5 (3.1) | 0.842 | 2.3 (2.3) | 0.748 | 85.7 (43.6) | 0.227 | 0.2 (0.4) | 0.662 | 0.5 (0.6) | 0.621 |

*Note*. E = 17β-estradiol; T = testosterone; P = progesterone; *b* = unstandardized coefficient; *SE* = standard error; *n* = analytic sample within each sample and timepoint; *n* values vary due to hormone-specific missing data. Primary regions (Crus I, Crus II) were evaluated uncorrected; secondary regions (Lobule V, VI) were FDR-corrected within family. *$p < .05$.

**Table S3.** Hormone-volume associations: Baseline, Year 1, and Baseline-predicting Year 1 models for exploratory regions, with outliers.

| | Baseline | | | | | | Year 1 | | | | | | Baseline → Year 1 | | | | | |
|---|---|---|---|---|---|---|---|---|---|---|---|---|---|---|---|---|---|---|
| | E | | T | | P | | E | | T | | P | | E | | T | | P | |
| Region | *b (SE)* | *p* | *b (SE)* | *p* | *b (SE)* | *p* | *b (SE)* | *p* | *b (SE)* | *p* | *b (SE)* | *p* | *b (SE)* | *p* | *b (SE)* | *p* | *b (SE)* | *p* |
| Left Lobule IX | 14.4 (55.8) | 0.917 | 0.1 (0.8) | 1.000 | 0.8 (0.6) | 0.387 | 26.4 (64.7) | 0.760 | 1.6 (1.3) | 0.664 | 0.5 (0.6) | 0.573 | -6.2 (21.2) | 0.887 | -0.2 (0.2) | 0.631 | -0.1 (0.2) | 0.945 |
| Right Lobule IX | 13.0 (58.5) | 0.917 | 0.7 (0.8) | 0.799 | 0.9 (0.7) | 0.387 | 38.9 (66.1) | 0.760 | 2.1 (1.3) | 0.664 | 0.7 (0.6) | 0.573 | -0.3 (20.4) | 0.987 | -0.2 (0.2) | 0.631 | 0.0 (0.2) | 0.945 |
| Vermis VI | -22.1 (26.4) | 0.697 | 0.4 (0.4) | 0.799 | -0.4 (0.3) | 0.387 | -15.0 (31.8) | 0.760 | 0.2 (0.6) | 0.763 | -0.2 (0.3) | 0.573 | -10.6 (9.8) | 0.565 | -0.1 (0.1) | 0.670 | 0.0 (0.1) | 0.945 |
| Vermis VII | -19.1 (13.7) | 0.552 | -0.1 (0.2) | 1.000 | -0.1 (0.2) | 0.619 | -26.9 (15.4) | 0.421 | -0.5 (0.3) | 0.664 | -0.1 (0.1) | 0.573 | 6.5 (6.1) | 0.565 | 0.0 (0.1) | 0.633 | 0.1 (0.1) | 0.494 |
| Vermis VIII | -21.3 (28.0) | 0.697 | 0.3 (0.4) | 0.799 | -0.1 (0.3) | 0.670 | -40.3 (34.7) | 0.574 | 0.2 (0.7) | 0.763 | -0.3 (0.3) | 0.573 | -11.5 (12.0) | 0.565 | -0.1 (0.1) | 0.670 | -0.2 (0.1) | 0.494 |
| Vermis IX | -16.2 (11.3) | 0.552 | 0.0 (0.2) | 1.000 | -0.1 (0.1) | 0.396 | -14.6 (13.7) | 0.574 | -0.2 (0.3) | 0.664 | -0.1 (0.1) | 0.573 | -2.0 (7.0) | 0.887 | 0.0 (0.1) | 0.633 | 0.0 (0.1) | 0.993 |
| Vermis X | -7.1 (6.3) | 0.660 | 0.0 (0.1) | 1.000 | -0.1 (0.1) | 0.387 | -1.1 (8.4) | 0.894 | -0.1 (0.2) | 0.664 | -0.1 (0.1) | 0.573 | -6.1 (3.4) | 0.565 | -0.1 (0.0) | 0.565 | 0.0 (0.0) | 0.602 |
| Total Gray Matter | -2095.9 (3015.1) | 0.697 | -26.4 (42.9) | 0.899 | -55.1 (33.9) | 0.387 | -6740.7 (3428.1) | 0.421 | -67.1 (69.9) | 0.664 | -1.9 (33.3) | 0.954 | 1896.9 (1485.2) | 0.565 | 20.2 (16.2) | 0.565 | 13.3 (12.5) | 0.602 |
| Cortical Volume | -65.1 (2639.7) | 0.980 | -48.5 (37.2) | 0.799 | -38.0 (29.5) | 0.387 | -4262.8 (3164.3) | 0.574 | -52.3 (63.8) | 0.664 | 12.5 (30.5) | 0.759 | 1839.9 (1346.0) | 0.565 | 20.6 (14.9) | 0.565 | 17.2 (11.3) | 0.494 |
| Cerebral White Matter | 7342.5 (3188.4) | 0.229 | 51.8 (46.2) | 0.799 | -15.6 (36.6) | 0.670 | -2020.7 (3892.9) | 0.760 | -45.2 (77.2) | 0.699 | -35.1 (36.7) | 0.573 | -288.4 (1126.2) | 0.887 | 14.8 (12.1) | 0.565 | -3.8 (9.4) | 0.945 |
| Left Hippocampus | -18.3 (40.1) | 0.860 | -0.3 (0.6) | 0.814 | 0.0 (0.5) | 0.985 | -100.3 (45.0) | 0.085 | -0.5 (0.9) | 0.835 | 0.1 (0.4) | 0.810 | 2.9 (13.1) | 0.956 | 0.2 (0.1) | 0.660 | -0.1 (0.1) | 0.917 |
| Right Hippocampus | 7.8 (41.2) | 0.860 | 0.1 (0.6) | 0.815 | 0.5 (0.5) | 0.432 | -51.6 (47.5) | 0.421 | -1.0 (0.9) | 0.835 | 0.2 (0.4) | 0.757 | -7.2 (13.6) | 0.956 | 0.0 (0.1) | 0.985 | -0.2 (0.1) | 0.481 |
| Left Amygdala | -2.9 (16.5) | 0.860 | -0.1 (0.2) | 0.815 | -0.4 (0.2) | 0.311 | **-53.7 (19.0)** | **0.035*** | -0.3 (0.4) | 0.835 | -0.2 (0.2) | 0.757 | 5.2 (10.9) | 0.956 | 0.1 (0.1) | 0.660 | 0.0 (0.1) | 0.917 |
| Right Amygdala | -8.7 (17.8) | 0.860 | -0.3 (0.3) | 0.598 | -0.3 (0.2) | 0.432 | -40.0 (20.8) | 0.115 | 0.0 (0.4) | 0.934 | -0.1 (0.2) | 0.757 | 10.0 (9.3) | 0.956 | 0.1 (0.1) | 0.660 | 0.0 (0.1) | 0.917 |
| Left Thalamus | 69.9 (66.8) | 0.860 | 0.6 (0.9) | 0.814 | 0.2 (0.8) | 0.949 | 48.1 (79.9) | 0.657 | 0.7 (1.6) | 0.835 | 0.4 (0.8) | 0.757 | -1.5 (27.8) | 0.956 | -0.1 (0.3) | 0.852 | 0.0 (0.2) | 0.917 |
| Right Thalamus | 119.4 | 0.322 | 1.6 (0.9) | 0.416 | 0.7 (0.7) | 0.432 | 32.3 (72.5) | 0.657 | 0.6 (1.5) | 0.835 | 0.7 (0.7) | 0.757 | -6.3 (32.7) | 0.956 | 0.3 (0.4) | 0.660 | -0.1 (0.3) | 0.917 |

| | E | | T | | P | | E | | T | | P | | E | | T | | P | |
|---|---|---|---|---|---|---|---|---|---|---|---|---|---|---|---|---|---|---|
| | Baseline | | | | | | Year 1 | | | | | | Baseline → Year 1 | | | | | |
| **Region** | *b (SE)* | *p* | *b (SE)* | *p* | *b (SE)* | *p* | *b (SE)* | *p* | *b (SE)* | *p* | *b (SE)* | *p* | *b (SE)* | *p* | *b (SE)* | *p* | *b (SE)* | *p* |
| Left Lobule IX | 14.4 (55.8) | 0.917 | 0.1 (0.8) | 1.000 | 0.8 (0.6) | 0.387 | 26.4 (64.7) | 0.760 | 1.6 (1.3) | 0.664 | 0.5 (0.6) | 0.573 | -6.2 (21.2) | 0.887 | -0.2 (0.2) | 0.631 | -0.1 (0.2) | 0.945 |
| Right Lobule IX | 13.0 (58.5) | 0.917 | 0.7 (0.8) | 0.799 | 0.9 (0.7) | 0.387 | 38.9 (66.1) | 0.760 | 2.1 (1.3) | 0.664 | 0.7 (0.6) | 0.573 | -0.3 (20.4) | 0.987 | -0.2 (0.2) | 0.631 | 0.0 (0.2) | 0.945 |
| Vermis VI | -22.1 (26.4) | 0.697 | 0.4 (0.4) | 0.799 | -0.4 (0.3) | 0.387 | -15.0 (31.8) | 0.760 | 0.2 (0.6) | 0.763 | -0.2 (0.3) | 0.573 | -10.6 (9.8) | 0.565 | -0.1 (0.1) | 0.670 | 0.0 (0.1) | 0.945 |
| Vermis VII | -19.1 (13.7) | 0.552 | -0.1 (0.2) | 1.000 | -0.1 (0.2) | 0.619 | -26.9 (15.4) | 0.421 | -0.5 (0.3) | 0.664 | -0.1 (0.1) | 0.573 | 6.5 (6.1) | 0.565 | 0.0 (0.1) | 0.633 | 0.1 (0.1) | 0.494 |
| Vermis VIII | -21.3 (28.0) | 0.697 | 0.3 (0.4) | 0.799 | -0.1 (0.3) | 0.670 | -40.3 (34.7) | 0.574 | 0.2 (0.7) | 0.763 | -0.3 (0.3) | 0.573 | -11.5 (12.0) | 0.565 | -0.1 (0.1) | 0.670 | -0.2 (0.1) | 0.494 |
| Vermis IX | -16.2 (11.3) | 0.552 | 0.0 (0.2) | 1.000 | -0.1 (0.1) | 0.396 | -14.6 (13.7) | 0.574 | -0.2 (0.3) | 0.664 | -0.1 (0.1) | 0.573 | -2.0 (7.0) | 0.887 | 0.0 (0.1) | 0.633 | 0.0 (0.1) | 0.993 |
| Vermis X | -7.1 (6.3) | 0.660 | 0.0 (0.1) | 1.000 | -0.1 (0.1) | 0.387 | -1.1 (8.4) | 0.894 | -0.1 (0.2) | 0.664 | -0.1 (0.1) | 0.573 | -6.1 (3.4) | 0.565 | -0.1 (0.0) | 0.565 | 0.0 (0.0) | 0.602 |
| Total Gray Matter | -2095.9 (3015.1) | 0.697 | -26.4 (42.9) | 0.899 | -55.1 (33.9) | 0.387 | -6740.7 (3428.1) | 0.421 | -67.1 (69.9) | 0.664 | -1.9 (33.3) | 0.954 | 1896.9 (1485.2) | 0.565 | 20.2 (16.2) | 0.565 | 13.3 (12.5) | 0.602 |
| Cortical Volume | -65.1 (2639.7) | 0.980 | -48.5 (37.2) | 0.799 | -38.0 (29.5) | 0.387 | -4262.8 (3164.3) | 0.574 | -52.3 (63.8) | 0.664 | 12.5 (30.5) | 0.759 | 1839.9 (1346.0) | 0.565 | 20.6 (14.9) | 0.565 | 17.2 (11.3) | 0.494 |
| Cerebral White Matter | 7342.5 (3188.4) | 0.229 | 51.8 (46.2) | 0.799 | -15.6 (36.6) | 0.670 | -2020.7 (3892.9) | 0.760 | -45.2 (77.2) | 0.699 | -35.1 (36.7) | 0.573 | -288.4 (1126.2) | 0.887 | 14.8 (12.1) | 0.565 | -3.8 (9.4) | 0.945 |
| Left Hippocampus | -18.3 (40.1) (61.4) | 0.860 | -0.3 (0.6) | 0.814 | 0.0 (0.5) | 0.985 | -100.3 (45.0) | 0.085 | -0.5 (0.9) | 0.835 | 0.1 (0.4) | 0.810 | 2.9 (13.1) | 0.956 | 0.2 (0.1) | 0.660 | -0.1 (0.1) | 0.917 |

*Note.* Full sample (sex included as a covariate). E = 17β-estradiol; T = testosterone; P = progesterone; b = unstandardized coefficient; *SE* = standard error. FDR correction was applied within analysis type: Lobule IX, vermis subregions, and global volumes were corrected together as one family; limbic regions (hippocampus, amygdala, thalamus) were corrected as a separate family. Prediction models control for baseline volume. *$p$_FDR < .05.

**Table S4.** Hormone-volume associations: Baseline, Year 1, and Baseline-predicting Year 1 models for exploratory regions, without outliers.

| | **Baseline** | | | | | | **Year 1** | | | | | | **Baseline → Year 1** | | | | | |
|---|---|---|---|---|---|---|---|---|---|---|---|---|---|---|---|---|---|---|
| | **E** | | **T** | | **P** | | **E** | | **T** | | **P** | | **E** | | **T** | | **P** | |
| **Region** | *b (SE)* | *p* | *b (SE)* | *p* | *b (SE)* | *p* | *b (SE)* | *p* | *b (SE)* | *p* | *b (SE)* | *p* | *b (SE)* | *P* | *b (SE)* | *p* | *b (SE)* | *p* |
| Left Lobule IX | 52.5 (78.2) | 0.553 | -0.4 (1.0) | 0.950 | 1.5 (0.8) | 0.368 | 236.1 (102.9) | 0.186 | 1.6 (1.4) | 0.503 | 0.9 (0.9) | 0.567 | -15.8 (23.8) | 0.585 | -0.4 (0.3) | 0.493 | -0.2 (0.3) | 0.857 |
| Right Lobule IX | 50.5 (82.8) | 0.553 | 0.5 (1.1) | 0.950 | 1.6 (0.9) | 0.368 | 226.7 (106.9) | 0.186 | 2.1 (1.4) | 0.503 | 1.3 (0.9) | 0.509 | -13.9 (22.1) | 0.585 | -0.4 (0.3) | 0.493 | -0.1 (0.2) | 0.857 |
| Vermis VI | -44.3 (36.5) | 0.553 | 0.5 (0.5) | 0.950 | -0.3 (0.4) | 0.744 | -0.3 (50.9) | 0.998 | -0.1 (0.7) | 0.821 | -0.3 (0.4) | 0.597 | -16.2 (10.7) | 0.550 | -0.2 (0.2) | 0.493 | -0.1 (0.1) | 0.857 |
| Vermis VII | -16.7 (18.8) | 0.553 | 0.0 (0.3) | 0.950 | -0.1 (0.2) | 0.744 | -25.0 (25.0) | 0.752 | -0.5 (0.3) | 0.503 | -0.3 (0.2) | 0.509 | 5.4 (6.9) | 0.585 | -0.1 (0.1) | 0.493 | 0.0 (0.1) | 0.901 |
| Vermis VIII | -39.7 (39.5) | 0.553 | 0.6 (0.5) | 0.950 | 0.2 (0.4) | 0.770 | 0.2 (56.8) | 0.998 | 0.5 (0.7) | 0.589 | -0.3 (0.5) | 0.597 | -13.7 (13.0) | 0.585 | 0.0 (0.2) | 0.939 | -0.2 (0.1) | 0.779 |
| Vermis IX | -30.0 (15.9) | 0.310 | 0.1 (0.2) | 0.950 | -0.2 (0.2) | 0.744 | -20.9 (22.2) | 0.752 | -0.2 (0.3) | 0.503 | -0.3 (0.2) | 0.509 | -4.7 (7.2) | 0.585 | 0.0 (0.1) | 0.939 | 0.0 (0.1) | 0.901 |
| Vermis X | -17.8 (8.6) | 0.310 | 0.0 (0.1) | 0.950 | -0.1 (0.1) | 0.744 | 0.9 (13.7) | 0.998 | -0.2 (0.2) | 0.503 | -0.1 (0.1) | 0.509 | -7.5 (3.4) | 0.316 | 0.0 (0.1) | 0.493 | 0.0 (0.0) | 0.857 |
| Total Gray Matter | -4490.0 (4185.2) | 0.553 | -20.5 (55.5) | 0.950 | -28.9 (44.8) | 0.744 | -5012.7 (5630.2) | 0.752 | -75.2 (74.1) | 0.503 | 11.5 (46.2) | 0.804 | 1576.9 (1610.9) | 0.585 | 23.1 (23.3) | 0.493 | 15.9 (17.4) | 0.857 |
| Cortical Volume | -2204.4 (3704.3) | 0.553 | -53.8 (48.6) | 0.950 | -15.5 (39.3) | 0.770 | -3609.9 (5288.8) | 0.828 | -62.5 (69.5) | 0.503 | 29.3 (43.1) | 0.597 | 2082.4 (1486.0) | 0.550 | 22.1 (21.9) | 0.493 | 23.1 (16.1) | 0.779 |
| Cerebral White Matter | 3043.6 (4375.0) | 0.553 | 6.3 (58.2) | 0.950 | 13.0 (46.6) | 0.781 | -2999.9 (6200.5) | 0.900 | -91.5 (80.2) | 0.503 | 32.8 (50.0) | 0.597 | -691.4 (1261.3) | 0.585 | 11.3 (18.1) | 0.666 | 1.6 (13.7) | 0.905 |
| Left Hippocampus | -62.7 (55.3) | 0.635 | 0.2 (0.7) | 0.888 | -0.1 (0.6) | 0.864 | -40.1 (74.3) | 0.709 | -0.6 (1.0) | 0.816 | 0.0 (0.6) | 0.943 | -2.2 (14.3) | 0.880 | 0.1 (0.2) | 0.943 | 0.0 (0.2) | 0.912 |
| Right Hippocampus | -36.7 (58.1) | 0.635 | 0.4 (0.8) | 0.888 | 0.4 (0.6) | 0.615 | -23.4 (78.7) | 0.767 | -1.4 (1.0) | 0.816 | -0.1 (0.6) | 0.943 | -10.9 (14.8) | 0.695 | 0.0 (0.2) | 0.943 | -0.2 (0.2) | 0.755 |
| Left Amygdala | -24.2 (22.4) | 0.635 | 0.0 (0.3) | 0.888 | -0.5 (0.2) | 0.179 | -36.2 (30.9) | 0.489 | -0.2 (0.4) | 0.816 | -0.2 (0.3) | 0.943 | 3.7 (12.1) | 0.880 | 0.3 (0.2) | 0.314 | 0.1 (0.1) | 0.912 |
| Right Amygdala | -18.5 (24.0) | 0.635 | 0.1 (0.3) | 0.888 | -0.3 (0.3) | 0.341 | -20.6 (32.5) | 0.709 | 0.1 (0.4) | 0.816 | 0.0 (0.3) | 0.943 | 7.3 (9.9) | 0.695 | 0.2 (0.1) | 0.401 | 0.1 (0.1) | 0.912 |
| Left Thalamus | 8.0 (95.2) | 0.933 | 0.9 (1.3) | 0.888 | 1.3 (1.0) | 0.341 | 158.0 (130.9) | 0.489 | 0.8 (1.7) | 0.816 | 2.1 (1.1) | 0.156 | -27.7 (31.1) | 0.695 | -0.6 (0.4) | 0.401 | 0.1 (0.3) | 0.912 |
| Right Thalamus | 67.9 (86.9) | 0.635 | 1.8 (1.1) | 0.700 | 1.7 (0.9) | 0.179 | 145.7 (117.0) | 0.489 | 0.4 (1.6) | 0.816 | 2.3 (0.9) | 0.089 | -30.4 (35.9) | 0.695 | 0.0 (0.5) | 0.943 | 0.0 (0.4) | 0.912 |

*Note.* Full sample with outlying values removed (11 subjects, 3*SD* within sex and timepoint). Sex included as a covariate. E = 17β-estradiol; T = testosterone; P = progesterone; *b* = unstandardized coefficient; *SE* = standard error. FDR correction was applied within analysis type: Lobule IX, vermis subregions, and global volumes were corrected together as one family; limbic regions (hippocampus, amygdala, thalamus) were corrected as a separate family. Prediction models control for baseline volume. *$p$_FDR < .05.

**Table S5.** Sex-stratified hormone-volume associations: Baseline, Year 1, and Baseline-predicting Year 1 models for exploratory regions, with outliers.

| | Baseline | | | | | | Year 1 | | | | | | Baseline → Year 1 | | | | | |
|---|---|---|---|---|---|---|---|---|---|---|---|---|---|---|---|---|---|---|
| | E | | T | | P | | E | | T | | P | | E | | T | | P | |
| **Region** | *b (SE)* | *p* | *b (SE)* | *p* | *b (SE)* | *p* | *b (SE)* | *p* | *b (SE)* | *p* | *b (SE)* | *p* | *b (SE)* | *p* | *b (SE)* | *p* | *b (SE)* | *p* |
| **Females** | | | | | | | | | | | | | | | | | | |
| Left Lobule IX | -42.0 (61.6) | 0.623 | -1.1 (1.5) | 0.744 | 0.9 (0.7) | 0.343 | 109.7 (92.7) | 0.447 | 1.6 (2.0) | 0.625 | 0.2 (0.7) | 0.803 | 4.1 (23.5) | 0.861 | -0.3 (0.4) | 0.801 | 0.0 (0.2) | 0.964 |
| Right Lobule IX | -44.0 (63.0) | 0.623 | -0.3 (1.5) | 0.874 | 1.1 (0.7) | 0.282 | 132.4 (94.8) | 0.424 | 1.6 (2.1) | 0.625 | 0.3 (0.7) | 0.803 | -15.6 (24.9) | 0.689 | -0.2 (0.4) | 0.801 | 0.1 (0.2) | 0.860 |
| Vermis VI | -51.1 (27.2) | 0.186 | -0.4 (0.7) | 0.744 | -0.5 (0.3) | 0.282 | -78.9 (44.2) | 0.407 | -0.9 (1.0) | 0.625 | -0.7 (0.3) | 0.125 | -7.2 (12.0) | 0.689 | -0.1 (0.2) | 0.801 | -0.1 (0.1) | 0.860 |
| Vermis VII | -26.3 (14.8) | 0.186 | -0.4 (0.4) | 0.744 | -0.1 (0.2) | 0.692 | -39.6 (19.9) | 0.407 | -0.1 (0.5) | 0.891 | -0.1 (0.1) | 0.803 | 5.0 (7.8) | 0.689 | -0.1 (0.1) | 0.801 | 0.1 (0.1) | 0.860 |
| Vermis VIII | -14.3 (30.6) | 0.714 | 0.1 (0.8) | 0.874 | 0.0 (0.3) | 0.976 | -40.2 (50.5) | 0.478 | -0.1 (1.1) | 0.957 | -0.5 (0.4) | 0.412 | 4.3 (13.2) | 0.826 | 0.0 (0.2) | 0.879 | 0.0 (0.1) | 0.860 |
| Vermis IX | -25.9 (11.7) | 0.153 | -0.3 (0.3) | 0.744 | -0.1 (0.1) | 0.505 | -16.1 (17.4) | 0.449 | -0.3 (0.4) | 0.625 | -0.3 (0.1) | 0.125 | -9.3 (8.9) | 0.689 | -0.1 (0.1) | 0.801 | 0.0 (0.1) | 0.860 |
| Vermis X | -11.8 (6.9) | 0.186 | -0.1 (0.2) | 0.744 | -0.1 (0.1) | 0.384 | -12.5 (12.2) | 0.447 | -0.3 (0.3) | 0.625 | **-0.3 (0.1)** | **0.016*** | -6.7 (5.3) | 0.689 | -0.1 (0.1) | 0.801 | 0.0 (0.0) | 0.860 |
| Total Gray Matter | -2041.8 (2985.1) | 0.623 | -167.6 (68.8) | 0.088 | -58.8 (32.3) | 0.282 | -6232.7 (4121.8) | 0.424 | -62.1 (91.3) | 0.625 | -20.5 (30.5) | 0.803 | 1952.3 (2085.3) | 0.689 | 29.8 (33.4) | 0.801 | 9.7 (15.5) | 0.860 |
| Cortical Volume | 476.8 (2684.8) | 0.860 | -163.9 (60.8) | 0.088 | -46.8 (28.7) | 0.282 | -4126.2 (4041.1) | 0.447 | -61.5 (88.2) | 0.625 | -2.2 (30.0) | 0.942 | 1561.3 (1885.7) | 0.689 | 21.0 (30.8) | 0.801 | 9.6 (14.2) | 0.860 |
| Cerebral White Matter | **11602.5 (3286.1)** | **0.008*** | -44.2 (86.3) | 0.763 | -1.3 (39.7) | 0.976 | -1109.0 (5341.3) | 0.836 | 144.7 (113.8) | 0.625 | -16.8 (39.6) | 0.803 | -1904.4 (1768.4) | 0.689 | 4.2 (27.4) | 0.879 | -4.8 (13.0) | 0.860 |
| Left Hippocampus | 55.8 (41.9) | 0.187 | -0.2 (1.0) | 0.961 | 0.4 (0.5) | 0.573 | -66.6 (59.6) | 0.809 | 0.4 (1.3) | 0.763 | 0.3 (0.4) | 0.922 | -14.9 (16.4) | 0.550 | -0.1 (0.3) | 0.930 | -0.1 (0.1) | 0.522 |
| Right Hippocampus | 73.3 (43.1) | 0.187 | 1.1 (1.1) | 0.620 | 0.9 (0.5) | 0.390 | 3.4 (63.2) | 0.957 | 0.7 (1.4) | 0.763 | 0.4 (0.5) | 0.922 | -4.4 (15.9) | 0.784 | 0.0 (0.3) | 0.930 | -0.1 (0.1) | 0.522 |
| Left Amygdala | 24.8 (18.3) | 0.187 | -0.6 (0.5) | 0.474 | -0.2 (0.2) | 0.573 | -37.8 (28.1) | 0.809 | -0.2 (0.6) | 0.763 | 0.0 (0.2) | 0.922 | 24.4 (13.4) | 0.453 | 0.3 (0.2) | 0.567 | 0.1 (0.1) | 0.522 |
| Right Amygdala | 28.0 (18.0) | 0.187 | 0.0 (0.4) | 0.961 | -0.1 (0.2) | 0.800 | 3.9 (27.7) | 0.957 | 0.3 (0.6) | 0.763 | 0.1 (0.2) | 0.922 | 8.9 (11.9) | 0.550 | 0.0 (0.2) | 0.930 | 0.1 (0.1) | 0.522 |
| Left Thalamus | 107.2 (80.4) | 0.187 | 1.0 (2.0) | 0.926 | 0.6 (0.9) | 0.573 | 28.6 (125.2) | 0.957 | 4.0 (2.6) | 0.429 | 0.2 (0.9) | 0.922 | -37.5 (39.1) | 0.550 | -0.6 (0.6) | 0.620 | -0.2 (0.3) | 0.522 |
| Right Thalamus | 144.0 (68.8) | 0.187 | 2.9 (1.7) | 0.474 | 1.1 (0.8) | 0.418 | 34.3 (103.0) | 0.957 | 3.9 (2.2) | 0.429 | 0.8 (0.8) | 0.922 | -54.9 (41.0) | 0.550 | -0.9 (0.7) | 0.567 | -0.5 (0.3) | 0.507 |
| **Males** | | | | | | | | | | | | | | | | | | |
| Left Lobule IX | 106.3 (97.3) | 0.963 | 0.6 (0.9) | 0.818 | -0.2 (1.4) | 0.944 | -3.4 (80.4) | 0.966 | 1.8 (1.5) | 0.528 | 0.8 (1.1) | 0.635 | -15.2 (37.5) | 0.765 | -0.3 (0.3) | 0.721 | -0.2 (0.5) | 0.792 |
| Right Lobule IX | 104.1 (106.3) | 0.963 | 1.2 (0.9) | 0.739 | -0.3 (1.5) | 0.944 | 20.0 (83.9) | 0.958 | 2.4 (1.5) | 0.503 | 1.6 (1.1) | 0.574 | 23.9 (33.1) | 0.676 | -0.3 (0.3) | 0.721 | 0.3 (0.4) | 0.792 |
| Vermis VI | 48.4 (55.8) | 0.963 | 0.6 (0.5) | 0.739 | 0.2 (0.8) | 0.944 | 32.2 (48.1) | 0.958 | 0.8 (0.9) | 0.706 | 1.1 (0.6) | 0.574 | -16.3 (18.2) | 0.676 | 0.0 (0.2) | 0.896 | 0.2 (0.2) | 0.792 |
| Vermis VII | -6.6 (27.9) | 0.963 | 0.1 (0.3) | 0.818 | -0.4 (0.4) | 0.911 | -12.7 (23.7) | 0.958 | -0.6 (0.4) | 0.503 | -0.3 (0.3) | 0.619 | 8.2 (10.1) | 0.676 | 0.0 (0.1) | 0.822 | 0.1 (0.1) | 0.792 |
| Vermis VIII | -57.0 (54.4) | 0.963 | 0.4 (0.5) | 0.818 | -1.0 (0.7) | 0.911 | -11.2 (48.4) | 0.958 | 0.5 (0.9) | 0.791 | 0.1 (0.6) | 0.834 | -26.4 (19.9) | 0.676 | -0.1 (0.2) | 0.776 | -0.2 (0.3) | 0.792 |
| Vermis IX | 1.1 (22.7) | 0.963 | 0.1 (0.2) | 0.818 | -0.4 (0.3) | 0.911 | -3.6 (20.4) | 0.958 | -0.1 (0.4) | 0.901 | 0.1 (0.3) | 0.740 | 2.4 (11.3) | 0.832 | 0.0 (0.1) | 0.822 | 0.1 (0.2) | 0.792 |
| Vermis X | 2.1 (12.8) | 0.963 | 0.1 (0.1) | 0.818 | -0.2 (0.2) | 0.911 | 12.3 (11.3) | 0.935 | 0.0 (0.2) | 0.901 | 0.2 (0.1) | 0.574 | -2.9 (4.8) | 0.687 | -0.1 (0.0) | 0.627 | 0.0 (0.1) | 0.827 |
| Total Gray Matter | -2553.4 (6515.9) | 0.963 | 14.2 (58.3) | 0.818 | -37.4 (91.1) | 0.944 | -7924.1 (5683.2) | 0.935 | -67.7 (107.3) | 0.791 | 52.6 (78.6) | 0.635 | 2011.6 (2234.6) | 0.676 | 19.6 (19.2) | 0.721 | 49.9 (29.9) | 0.514 |
| Cortical Volume | -1114.8 (5618.6) | 0.963 | -15.3 (50.2) | 0.818 | 5.5 (78.4) | 0.944 | -5775.3 (4906.8) | 0.935 | -44.4 (92.3) | 0.791 | 60.8 (67.2) | 0.619 | 2091.4 (2036.7) | 0.676 | 23.4 (17.5) | 0.627 | 47.2 (27.0) | 0.514 |

| | Baseline | | | | | | Year 1 | | | | | | Baseline → Year 1 | | | | | |
|---|---|---|---|---|---|---|---|---|---|---|---|---|---|---|---|---|---|---|
| | E | | T | | P | | E | | T | | P | | E | | T | | P | |
| **Region** | *b (SE)* | *p* | *b (SE)* | *p* | *b (SE)* | *p* | *b (SE)* | *p* | *b (SE)* | *p* | *b (SE)* | *p* | *b (SE)* | *p* | *b (SE)* | *p* | *b (SE)* | *p* |
| Cerebral White Matter | -3447.8 (6711.6) | 0.963 | 86.0 (59.1) | 0.739 | -67.4 (92.6) | 0.939 | -2669.8 (6077.1) | 0.958 | -156.6 (106.9) | 0.503 | -96.5 (79.0) | 0.574 | 1840.7 (1392.1) | 0.676 | 18.0 (12.0) | 0.627 | -2.8 (19.5) | 0.884 |
| Left Hippocampus | **-221.4 (75.1)** | **0.019*** | -0.5 (0.7) | 0.804 | -1.1 (1.1) | 0.475 | -103.1 (70.9) | 0.308 | -1.3 (1.3) | 0.668 | -0.3 (0.9) | 0.925 | 40.9 (20.7) | 0.329 | 0.3 (0.2) | 0.287 | 0.1 (0.3) | 0.778 |
| Right Hippocampus | -166.6 (79.1) | 0.059 | -0.3 (0.7) | 0.804 | -0.2 (1.1) | 0.881 | -71.3 (73.2) | 0.504 | -2.2 (1.3) | 0.513 | -0.1 (1.0) | 0.925 | -1.9 (23.4) | 0.934 | 0.0 (0.2) | 0.974 | -0.1 (0.3) | 0.778 |
| Left Amygdala | -70.4 (31.0) | 0.053 | 0.0 (0.3) | 0.959 | -0.6 (0.4) | 0.475 | -60.6 (27.3) | 0.098 | -0.5 (0.5) | 0.668 | -0.5 (0.4) | 0.925 | -13.1 (18.5) | 0.582 | 0.0 (0.1) | 0.974 | 0.2 (0.2) | 0.778 |
| Right Amygdala | **-98.5 (34.8)** | **0.019*** | -0.6 (0.3) | 0.563 | -0.8 (0.5) | 0.475 | -76.8 (31.3) | 0.098 | -0.4 (0.6) | 0.688 | -0.3 (0.5) | 0.925 | 20.7 (16.8) | 0.530 | 0.2 (0.1) | 0.287 | 0.4 (0.2) | 0.264 |
| Left Thalamus | -45.1 (122.2) | 0.718 | 0.5 (1.1) | 0.804 | -1.7 (1.7) | 0.475 | 81.6 (107.7) | 0.543 | -0.8 (2.0) | 0.688 | 0.7 (1.5) | 0.925 | 39.9 (40.7) | 0.530 | 0.1 (0.4) | 0.974 | 0.2 (0.6) | 0.778 |
| Right Thalamus | 44.2 (121.9) | 0.718 | 1.1 (1.1) | 0.804 | -0.7 (1.7) | 0.839 | 42.6 (106.1) | 0.690 | -0.9 (2.0) | 0.688 | 0.1 (1.4) | 0.925 | 49.7 (52.8) | 0.530 | 0.7 (0.4) | 0.287 | 1.0 (0.7) | 0.487 |

*Note.* Sex-stratified (models fit separately within each sex; sex not included as a covariate). E = 17β-estradiol; T = testosterone; P = progesterone; b = unstandardized coefficient; *SE* = standard error. FDR correction was applied within analysis type and sex: Lobule IX, vermis subregions, and global volumes were corrected together as one family; limbic regions (hippocampus, amygdala, thalamus) were corrected as a separate family. Prediction models control for baseline volume. **p*_FDR < .05.

**Table S6.** Sex-stratified hormone-volume associations: Baseline, Year 1, and Baseline-predicting Year 1 models for exploratory regions, without outliers.

| | **Baseline** | | | | | | **Year 1** | | | | | | **Baseline → Year 1** | | | | | |
|---|---|---|---|---|---|---|---|---|---|---|---|---|---|---|---|---|---|---|
| | **E** | | **T** | | **P** | | **E** | | **T** | | **P** | | **E** | | **T** | | **P** | |
| **Region** | *b (SE)* | *p* | *b (SE)* | *p* | *b (SE)* | *p* | *b (SE)* | *p* | *b (SE)* | *p* | *b (SE)* | *p* | *b (SE)* | *p* | *b (SE)* | *p* | *b (SE)* | *p* |
| **Females** | | | | | | | | | | | | | | | | | | |
| Left Lobule IX | -29.5 (94.8) | 0.757 | -1.0 (1.7) | 0.821 | 2.1 (0.9) | 0.121 | 269.6 (133.0) | 0.422 | 2.5 (2.1) | 0.849 | 1.2 (1.0) | 0.394 | 6.6 (24.3) | 0.794 | -0.2 (0.4) | 0.778 | 0.2 (0.3) | 0.811 |
| Right Lobule IX | -32.8 (98.1) | 0.757 | -0.3 (1.7) | 0.958 | 2.3 (0.9) | 0.121 | 248.7 (140.4) | 0.422 | 2.2 (2.2) | 0.849 | 1.1 (1.0) | 0.436 | -15.0 (26.3) | 0.794 | -0.2 (0.4) | 0.778 | 0.3 (0.3) | 0.811 |
| Vermis VI | -85.7 (40.5) | 0.096 | 0.0 (0.8) | 0.981 | -0.4 (0.4) | 0.568 | -65.7 (65.3) | 0.534 | -0.5 (1.0) | 0.849 | -0.7 (0.5) | 0.394 | -9.8 (12.8) | 0.746 | -0.1 (0.2) | 0.778 | -0.1 (0.1) | 0.811 |
| Vermis VII | -28.4 (21.9) | 0.332 | -0.3 (0.4) | 0.821 | -0.2 (0.2) | 0.568 | -35.0 (30.2) | 0.534 | -0.1 (0.5) | 0.914 | -0.1 (0.2) | 0.793 | 2.2 (8.2) | 0.794 | -0.2 (0.1) | 0.778 | 0.0 (0.1) | 0.894 |
| Vermis VIII | -28.1 (46.5) | 0.685 | 0.5 (0.8) | 0.821 | 0.7 (0.4) | 0.394 | -32.1 (71.7) | 0.657 | 0.6 (1.1) | 0.849 | -0.6 (0.5) | 0.394 | 5.2 (13.2) | 0.794 | 0.1 (0.2) | 0.778 | 0.0 (0.1) | 0.894 |
| Vermis IX | **-54.4 (17.7)** | **0.019*** | -0.2 (0.3) | 0.821 | -0.1 (0.2) | 0.568 | -31.1 (25.4) | 0.534 | -0.2 (0.4) | 0.849 | -0.3 (0.2) | 0.242 | -9.2 (8.6) | 0.746 | 0.1 (0.1) | 0.778 | 0.0 (0.1) | 0.894 |
| Vermis X | **-29.9 (10.0)** | **0.019*** | -0.1 (0.2) | 0.821 | -0.1 (0.1) | 0.568 | -19.1 (17.3) | 0.534 | -0.2 (0.3) | 0.849 | -0.3 (0.1) | 0.242 | -6.3 (5.3) | 0.746 | 0.0 (0.1) | 0.778 | 0.0 (0.1) | 0.894 |
| Total Gray Matter | -6505.4 (4155.8) | 0.245 | -99.3 (70.7) | 0.821 | -26.5 (40.7) | 0.568 | -3146.7 (5241.7) | 0.657 | 4.0 (81.0) | 0.961 | -10.3 (38.0) | 0.876 | 2084.5 (2246.8) | 0.746 | 34.1 (36.5) | 0.778 | 26.9 (22.8) | 0.811 |
| Cortical Volume | -3057.5 (3777.9) | 0.602 | -98.2 (62.2) | 0.821 | -24.2 (35.8) | 0.568 | -2664.9 (5518.5) | 0.657 | -19.2 (85.0) | 0.914 | 4.0 (40.0) | 0.920 | 1597.7 (2006.7) | 0.746 | 23.1 (33.4) | 0.778 | 24.2 (20.7) | 0.811 |
| Cerebral White Matter | **11998.2 (4660.2)** | **0.042*** | 38.0 (85.1) | 0.821 | 47.0 (48.0) | 0.568 | -4609.4 (7329.5) | 0.657 | 175.9 (109.7) | 0.849 | 94.1 (51.0) | 0.242 | -1528.9 (1875.7) | 0.746 | 8.5 (29.9) | 0.778 | 6.9 (19.0) | 0.894 |
| Left Hippocampus | 40.2 (65.0) | 0.694 | -0.1 (1.1) | 0.906 | 0.6 (0.6) | 0.467 | -13.8 (89.7) | 0.890 | 0.4 (1.4) | 0.929 | 0.3 (0.6) | 0.960 | -12.0 (16.3) | 0.576 | 0.1 (0.3) | 0.693 | 0.1 (0.2) | 0.718 |
| Right Hippocampus | 64.3 (66.6) | 0.694 | 0.8 (1.2) | 0.592 | 1.0 (0.6) | 0.199 | 13.0 (93.2) | 0.890 | 0.1 (1.4) | 0.929 | 0.0 (0.7) | 0.960 | -5.6 (15.9) | 0.728 | 0.2 (0.3) | 0.693 | -0.1 (0.2) | 0.790 |
| Left Amygdala | -0.2 (26.9) | 0.995 | -0.5 (0.5) | 0.592 | -0.4 (0.3) | 0.199 | -21.4 (38.1) | 0.890 | -0.1 (0.6) | 0.929 | 0.0 (0.3) | 0.960 | 26.4 (13.6) | 0.351 | 0.5 (0.2) | 0.099 | 0.2 (0.1) | 0.718 |
| Right Amygdala | 15.3 (27.4) | 0.694 | 0.4 (0.5) | 0.592 | 0.0 (0.3) | 0.896 | 14.6 (38.7) | 0.890 | 0.4 (0.6) | 0.909 | 0.1 (0.3) | 0.960 | 8.7 (12.2) | 0.576 | 0.1 (0.2) | 0.693 | 0.1 (0.1) | 0.718 |
| Left Thalamus | 104.9 (124.1) | 0.694 | 2.2 (2.1) | 0.592 | 2.4 (1.2) | 0.137 | 245.4 (178.2) | 0.529 | 5.2 (2.7) | 0.173 | 2.7 (1.2) | 0.119 | -38.9 (40.7) | 0.576 | -0.3 (0.7) | 0.693 | 0.1 (0.5) | 0.790 |
| Right Thalamus | 141.8 (107.0) | 0.694 | 3.2 (1.8) | 0.528 | 2.5 (1.0) | 0.092 | 258.2 (146.0) | 0.508 | 4.7 (2.2) | 0.173 | **3.2 (1.0)** | **0.015*** | -51.1 (41.6) | 0.576 | -0.4 (0.7) | 0.693 | -0.4 (0.5) | 0.718 |
| **Males** | | | | | | | | | | | | | | | | | | |
| Left Lobule IX | 133.6 (117.9) | 0.821 | 0.1 (1.3) | 0.933 | -0.5 (1.6) | 0.865 | 158.9 (142.6) | 0.746 | 1.6 (1.6) | 0.730 | 1.1 (1.4) | 0.673 | -49.2 (48.1) | 0.573 | -0.7 (0.6) | 0.616 | -1.1 (0.6) | 0.484 |
| Right Lobule IX | 139.9 (129.5) | 0.821 | 1.0 (1.4) | 0.933 | -0.5 (1.8) | 0.865 | 197.6 (149.0) | 0.746 | 2.2 (1.7) | 0.681 | 2.3 (1.4) | 0.673 | -4.8 (40.6) | 0.907 | -0.7 (0.5) | 0.616 | -0.6 (0.5) | 0.522 |

| | Baseline | | | | | | Year 1 | | | | | | Baseline → Year 1 | | | | | |
|---|---|---|---|---|---|---|---|---|---|---|---|---|---|---|---|---|---|---|
| | E | | T | | P | | E | | T | | P | | E | | T | | P | |
| **Region** | *b (SE)* | *p* | *b (SE)* | *p* | *b (SE)* | *p* | *b (SE)* | *p* | *b (SE)* | *p* | *b (SE)* | *p* | *b (SE)* | *p* | *b (SE)* | *p* | *b (SE)* | *p* |
| Vermis VI | 16.2 (67.0) | 0.993 | 0.7 (0.7) | 0.933 | -0.3 (0.9) | 0.865 | 67.9 (81.8) | 0.746 | 0.1 (0.9) | 0.872 | 0.6 (0.8) | 0.673 | -30.6 (21.0) | 0.573 | -0.2 (0.3) | 0.616 | -0.3 (0.3) | 0.512 |
| Vermis VII | -2.9 (33.5) | 0.993 | 0.2 (0.4) | 0.933 | -0.2 (0.5) | 0.865 | -18.0 (41.8) | 0.746 | -0.7 (0.5) | 0.623 | -0.6 (0.4) | 0.673 | 12.5 (13.0) | 0.573 | -0.1 (0.2) | 0.623 | 0.1 (0.2) | 0.678 |
| Vermis VIII | -66.4 (67.3) | 0.821 | 0.8 (0.7) | 0.933 | -1.4 (0.9) | 0.715 | 32.5 (88.3) | 0.746 | 0.6 (1.0) | 0.786 | 0.4 (0.8) | 0.677 | -40.8 (25.0) | 0.573 | -0.2 (0.3) | 0.638 | -0.7 (0.3) | 0.372 |
| Vermis IX | 3.7 (27.5) | 0.993 | 0.3 (0.3) | 0.933 | -0.5 (0.4) | 0.715 | -15.4 (36.6) | 0.746 | -0.2 (0.4) | 0.786 | -0.2 (0.4) | 0.677 | -1.8 (13.6) | 0.907 | 0.0 (0.2) | 0.883 | -0.2 (0.2) | 0.512 |
| Vermis X | -1.4 (15.1) | 0.993 | 0.1 (0.2) | 0.933 | -0.2 (0.2) | 0.715 | 18.7 (20.2) | 0.746 | -0.1 (0.2) | 0.786 | 0.2 (0.2) | 0.673 | -7.1 (5.1) | 0.573 | -0.1 (0.1) | 0.616 | -0.1 (0.1) | 0.512 |
| Total Gray Matter | -613.5 (8011.5) | 0.993 | 30.4 (84.6) | 0.933 | -33.0 (110.0) | 0.865 | -8421.4 (10348.4) | 0.746 | -109.6 (119.3) | 0.730 | 42.9 (102.0) | 0.677 | 1067.3 (2522.4) | 0.844 | 16.7 (31.2) | 0.663 | -19.9 (35.6) | 0.678 |
| Cortical Volume | 65.4 (6948.6) | 0.993 | -21.5 (73.4) | 0.933 | 10.9 (95.3) | 0.909 | -5587.1 (8973.5) | 0.746 | -75.9 (103.6) | 0.782 | 64.0 (87.8) | 0.673 | 2756.9 (2404.2) | 0.573 | 28.3 (30.4) | 0.616 | 14.3 (34.3) | 0.678 |
| Cerebral White Matter | -11641.4 (7838.3) | 0.821 | 10.8 (84.5) | 0.933 | -116.0 (107.6) | 0.715 | -3403.3 (10426.4) | 0.746 | -238.7 (110.5) | 0.381 | -95.9 (98.5) | 0.673 | 1543.8 (1828.8) | 0.578 | 21.5 (22.3) | 0.616 | -11.7 (25.7) | 0.678 |
| Left Hippocampus | -215.8 (90.3) | 0.122 | 0.3 (1.0) | 0.989 | -1.5 (1.3) | 0.545 | -59.6 (126.6) | 0.980 | -1.3 (1.4) | 0.645 | -0.9 (1.2) | 0.686 | 30.7 (26.2) | 0.748 | 0.1 (0.3) | 0.839 | -0.1 (0.4) | 0.829 |
| Right Hippocampus | -178.2 (97.1) | 0.216 | 0.0 (1.1) | 0.989 | -0.3 (1.4) | 0.946 | -43.7 (133.2) | 0.980 | -2.7 (1.4) | 0.373 | -0.6 (1.2) | 0.686 | -13.2 (30.2) | 0.882 | -0.1 (0.4) | 0.839 | -0.6 (0.4) | 0.801 |
| Left Amygdala | -52.4 (37.7) | 0.256 | 0.2 (0.4) | 0.989 | -0.5 (0.5) | 0.545 | -45.5 (50.2) | 0.980 | -0.4 (0.6) | 0.645 | -0.6 (0.5) | 0.686 | -28.6 (21.7) | 0.748 | 0.1 (0.3) | 0.839 | -0.1 (0.3) | 0.829 |
| Right Amygdala | -61.1 (41.5) | 0.256 | -0.2 (0.4) | 0.989 | -0.8 (0.6) | 0.545 | -52.7 (54.1) | 0.980 | -0.2 (0.6) | 0.794 | -0.5 (0.5) | 0.686 | 14.6 (18.9) | 0.882 | 0.2 (0.2) | 0.839 | 0.2 (0.3) | 0.829 |
| Left Thalamus | -153.8 (147.4) | 0.362 | 0.3 (1.6) | 0.989 | -1.9 (2.0) | 0.545 | 4.9 (192.1) | 0.980 | -1.4 (2.2) | 0.645 | 1.0 (1.9) | 0.686 | -17.6 (51.6) | 0.882 | -0.9 (0.6) | 0.839 | -0.8 (0.7) | 0.801 |
| Right Thalamus | -48.9 (145.8) | 0.739 | 1.1 (1.5) | 0.989 | -0.1 (2.0) | 0.946 | -37.3 (184.3) | 0.980 | -1.7 (2.1) | 0.645 | 0.7 (1.8) | 0.686 | -2.5 (65.9) | 0.970 | 0.6 (0.8) | 0.839 | 0.1 (0.9) | 0.924 |

*Note.* Sex-stratified with outlying values removed (11 subjects, 3*SD* within sex and timepoint); models fit separately within each sex (sex not included as a covariate). E = 17β-estradiol; T = testosterone; P = progesterone; *b* = unstandardized coefficient; *SE* = standard error. FDR correction was applied within analysis type and sex: Lobule IX, vermis subregions, and global volumes were corrected together as one family; limbic regions (hippocampus, amygdala, thalamus) were corrected as a separate family. Prediction models control for baseline volume. *$p$_FDR < .05.

**Table S7.** Hormone-volume change-score associations for exploratory regions, with outliers.

| | E | | T | | P | |
|---|---|---|---|---|---|---|
| **Region** | *b (SE)* | *p* | *b (SE)* | *p* | *b (SE)* | *p* |
| **Full sample** | | | | | | |
| Left Lobule IX | 9.7 (16.9) | 0.898 | 0.0 (0.3) | 0.960 | 0.2 (0.2) | 0.793 |
| Right Lobule IX | 8.9 (16.5) | 0.898 | 0.0 (0.3) | 0.960 | 0.1 (0.2) | 0.793 |
| Vermis VI | 3.2 (7.6) | 0.898 | 0.0 (0.1) | 0.960 | -0.1 (0.1) | 0.793 |
| Vermis VII | 0.6 (4.8) | 0.951 | 0.1 (0.1) | 0.960 | 0.1 (0.1) | 0.665 |
| Vermis VIII | -4.0 (9.2) | 0.898 | -0.1 (0.2) | 0.960 | 0.0 (0.1) | 0.895 |
| Vermis IX | 3.4 (5.5) | 0.898 | 0.1 (0.1) | 0.847 | 0.0 (0.1) | 0.895 |
| Vermis X | 2.1 (2.6) | 0.898 | 0.1 (0.0) | 0.847 | 0.0 (0.0) | 0.793 |
| Total Gray Matter | 69.0 (1123.3) | 0.951 | 14.8 (18.3) | 0.960 | -1.8 (13.8) | 0.895 |
| Cortical Volume | -374.6 (1036.7) | 0.898 | 7.4 (17.0) | 0.960 | -7.8 (12.8) | 0.793 |
| Cerebral White Matter | 861.8 (867.1) | 0.898 | -2.9 (14.3) | 0.960 | -6.8 (10.8) | 0.793 |
| Left Hippocampus | 7.5 (10.5) | 0.951 | -0.2 (0.2) | 0.898 | 0.0 (0.1) | 0.982 |
| Right Hippocampus | -2.2 (10.6) | 0.998 | 0.0 (0.2) | 0.898 | 0.1 (0.1) | 0.982 |
| Left Amygdala | 0.0 (8.4) | 0.998 | 0.1 (0.1) | 0.898 | 0.0 (0.1) | 0.982 |
| Right Amygdala | -9.4 (6.9) | 0.951 | 0.0 (0.1) | 0.898 | -0.1 (0.1) | 0.982 |
| Left Thalamus | 20.4 (21.1) | 0.951 | 0.0 (0.3) | 0.898 | -0.1 (0.3) | 0.982 |
| Right Thalamus | 8.6 (24.8) | 0.998 | -0.1 (0.4) | 0.898 | 0.1 (0.3) | 0.982 |
| **Females** | | | | | | |
| Left Lobule IX | -7.9 (19.2) | 0.963 | -0.8 (0.5) | 0.372 | 0.1 (0.2) | 0.778 |
| Right Lobule IX | -9.0 (20.6) | 0.963 | -0.8 (0.5) | 0.372 | 0.1 (0.2) | 0.905 |
| Vermis VI | -3.5 (8.9) | 0.963 | -0.2 (0.2) | 0.723 | -0.2 (0.1) | 0.576 |
| Vermis VII | 1.0 (6.1) | 0.963 | 0.4 (0.2) | 0.232 | 0.2 (0.1) | 0.072 |
| Vermis VIII | -10.7 (9.8) | 0.963 | -0.5 (0.3) | 0.372 | -0.1 (0.1) | 0.715 |
| Vermis IX | -3.6 (6.8) | 0.963 | 0.0 (0.2) | 0.972 | 0.0 (0.1) | 0.965 |
| Vermis X | 2.0 (3.8) | 0.963 | 0.1 (0.1) | 0.972 | 0.0 (0.0) | 0.770 |
| Total Gray Matter | -70.7 (1515.6) | 0.963 | 2.3 (42.5) | 0.972 | -20.0 (17.8) | 0.664 |
| Cortical Volume | 214.9 (1392.3) | 0.963 | 1.4 (39.1) | 0.972 | -22.6 (16.3) | 0.576 |
| Cerebral White Matter | 801.6 (1345.2) | 0.963 | -1.6 (37.2) | 0.972 | -14.4 (15.9) | 0.715 |
| Left Hippocampus | 9.6 (14.2) | 0.781 | 0.0 (0.4) | 0.978 | 0.0 (0.2) | 0.993 |
| Right Hippocampus | -11.0 (12.8) | 0.781 | 0.0 (0.4) | 0.978 | 0.1 (0.1) | 0.751 |
| Left Amygdala | -2.2 (10.9) | 0.840 | -0.1 (0.3) | 0.978 | -0.1 (0.1) | 0.751 |
| Right Amygdala | -15.2 (8.7) | 0.516 | 0.0 (0.2) | 0.978 | -0.2 (0.1) | 0.604 |
| Left Thalamus | 14.6 (29.6) | 0.781 | 0.1 (0.8) | 0.978 | -0.3 (0.3) | 0.751 |
| Right Thalamus | 14.1 (30.9) | 0.781 | 0.9 (0.9) | 0.978 | 0.0 (0.4) | 0.993 |
| **Males** | | | | | | |
| Left Lobule IX | 48.9 (29.7) | 0.451 | 0.3 (0.4) | 0.895 | 0.1 (0.4) | 0.844 |
| Right Lobule IX | 45.2 (26.8) | 0.451 | 0.3 (0.3) | 0.895 | 0.1 (0.4) | 0.844 |
| Vermis VI | 12.3 (13.8) | 0.542 | 0.0 (0.2) | 0.973 | 0.1 (0.2) | 0.844 |
| Vermis VII | 1.1 (8.3) | 0.891 | 0.0 (0.1) | 0.973 | 0.0 (0.1) | 0.844 |
| Vermis VIII | 15.4 (16.1) | 0.542 | 0.0 (0.2) | 0.985 | -0.1 (0.2) | 0.844 |
| Vermis IX | 14.2 (9.3) | 0.451 | 0.1 (0.1) | 0.895 | 0.0 (0.1) | 0.844 |
| Vermis X | 3.3 (3.7) | 0.542 | 0.1 (0.0) | 0.895 | 0.0 (0.1) | 0.844 |
| Total Gray Matter | 546.2 (1818.5) | 0.851 | 17.6 (20.8) | 0.895 | 29.9 (23.7) | 0.844 |
| Cortical Volume | -1286.1 (1668.0) | 0.557 | 8.3 (19.5) | 0.973 | 22.9 (22.1) | 0.844 |
| Cerebral White Matter | 1044.0 (1158.8) | 0.542 | -2.3 (13.7) | 0.973 | 7.3 (15.7) | 0.844 |
| Left Hippocampus | 10.2 (16.7) | 0.816 | -0.2 (0.2) | 0.819 | -0.1 (0.2) | 0.909 |

| | E | | T | | P | |
|---|---|---|---|---|---|---|
| **Region** | *b (SE)* | *p* | *b (SE)* | *p* | *b (SE)* | *p* |
| Right Hippocampus | 19.5 (17.8) | 0.816 | 0.0 (0.2) | 0.954 | 0.0 (0.2) | 0.909 |
| Left Amygdala | 10.2 (13.6) | 0.816 | 0.2 (0.2) | 0.819 | 0.0 (0.2) | 0.909 |
| Right Amygdala | 2.4 (11.5) | 0.958 | 0.0 (0.1) | 0.954 | 0.0 (0.2) | 0.909 |
| Left Thalamus | 23.7 (32.8) | 0.816 | 0.1 (0.4) | 0.954 | 0.4 (0.4) | 0.909 |
| Right Thalamus | 2.3 (42.8) | 0.958 | -0.4 (0.5) | 0.819 | 0.4 (0.5) | 0.909 |

*Note.* Change-score models. Full sample includes sex as a covariate; sex-stratified sections were fit separately within each sex. E = 17β-estradiol; T = testosterone; P = progesterone; *b* = unstandardized coefficient; *SE* = standard error. FDR within family (exploratory: Lobule IX, vermis, global together; limbic separate). *$p$_FDR < .05.

**Table S8.** Hormone-volume change-score associations for exploratory regions, without outliers.

| | E | | T | | P | |
|---|---|---|---|---|---|---|
| **Region** | *b (SE)* | *p* | *b (SE)* | *p* | *b (SE)* | *p* |
| **Full sample** | | | | | | |
| Left Lobule IX | 22.7 (26.1) | 0.555 | -0.2 (0.4) | 0.795 | 0.0 (0.3) | 0.920 |
| Right Lobule IX | 14.1 (24.6) | 0.632 | -0.3 (0.4) | 0.667 | 0.0 (0.3) | 0.920 |
| Vermis VI | 23.3 (11.1) | 0.389 | 0.0 (0.2) | 0.795 | -0.1 (0.1) | 0.920 |
| Vermis VII | 5.7 (7.5) | 0.557 | 0.2 (0.1) | 0.667 | 0.1 (0.1) | 0.920 |
| Vermis VIII | -5.3 (13.7) | 0.701 | -0.4 (0.2) | 0.666 | -0.1 (0.1) | 0.920 |
| Vermis IX | 7.0 (8.0) | 0.555 | 0.1 (0.1) | 0.667 | 0.0 (0.1) | 0.920 |
| Vermis X | 6.2 (3.7) | 0.413 | 0.0 (0.1) | 0.795 | 0.0 (0.0) | 0.920 |
| Total Gray Matter | 2543.9 (1635.7) | 0.413 | 28.2 (25.0) | 0.667 | 4.8 (16.7) | 0.920 |
| Cortical Volume | 1601.3 (1555.8) | 0.555 | 25.4 (23.6) | 0.667 | 2.8 (15.8) | 0.920 |
| Cerebral White Matter | 1803.3 (1340.5) | 0.456 | 8.7 (20.6) | 0.795 | 4.3 (13.7) | 0.920 |
| Left Hippocampus | 21.0 (15.3) | 0.350 | 0.0 (0.2) | 0.973 | 0.0 (0.2) | 0.847 |
| Right Hippocampus | 15.5 (15.6) | 0.484 | 0.0 (0.2) | 0.973 | 0.2 (0.2) | 0.847 |
| Left Amygdala | 1.1 (12.8) | 0.935 | 0.0 (0.2) | 0.973 | -0.1 (0.1) | 0.847 |
| Right Amygdala | -4.1 (10.3) | 0.828 | 0.0 (0.2) | 0.973 | 0.0 (0.1) | 0.847 |
| Left Thalamus | 68.2 (31.8) | 0.142 | 0.3 (0.5) | 0.973 | -0.1 (0.3) | 0.847 |
| Right Thalamus | 73.7 (36.6) | 0.142 | 0.0 (0.6) | 0.973 | 0.3 (0.4) | 0.847 |
| **Females** | | | | | | |
| Left Lobule IX | -26.0 (30.6) | 0.655 | -0.9 (0.6) | 0.308 | -0.2 (0.3) | 0.615 |
| Right Lobule IX | -21.9 (33.5) | 0.655 | -1.0 (0.6) | 0.308 | -0.2 (0.3) | 0.615 |
| Vermis VI | 8.5 (14.3) | 0.655 | -0.2 (0.3) | 0.881 | -0.2 (0.1) | 0.404 |
| Vermis VII | 5.3 (9.7) | 0.655 | 0.5 (0.2) | 0.050 | 0.2 (0.1) | 0.052 |
| Vermis VIII | -30.1 (14.4) | 0.433 | -0.7 (0.3) | 0.050 | -0.4 (0.1) | 0.052 |
| Vermis IX | -2.4 (10.4) | 0.821 | -0.1 (0.2) | 0.948 | -0.1 (0.1) | 0.615 |
| Vermis X | 7.2 (5.9) | 0.655 | 0.0 (0.1) | 0.981 | -0.1 (0.1) | 0.615 |
| Total Gray Matter | 1566.3 (2447.5) | 0.655 | 1.1 (47.0) | 0.981 | -20.6 (24.8) | 0.615 |
| Cortical Volume | 2038.7 (2208.3) | 0.655 | 7.7 (42.7) | 0.981 | -16.5 (22.7) | 0.615 |
| Cerebral White Matter | 2789.2 (2126.0) | 0.655 | 4.6 (40.7) | 0.981 | -7.0 (21.9) | 0.750 |
| Left Hippocampus | 18.1 (21.1) | 0.482 | -0.2 (0.4) | 0.724 | 0.1 (0.2) | 0.761 |
| Right Hippocampus | -0.6 (19.1) | 0.977 | -0.2 (0.4) | 0.724 | 0.2 (0.2) | 0.675 |
| Left Amygdala | -14.6 (17.2) | 0.482 | -0.3 (0.3) | 0.724 | -0.2 (0.2) | 0.675 |
| Right Amygdala | -18.6 (13.7) | 0.365 | -0.1 (0.3) | 0.724 | -0.2 (0.1) | 0.675 |
| Left Thalamus | 94.7 (44.5) | 0.162 | -0.3 (0.9) | 0.724 | -0.3 (0.5) | 0.675 |
| Right Thalamus | 89.9 (45.3) | 0.162 | 0.6 (0.9) | 0.724 | 0.3 (0.5) | 0.675 |
| **Males** | | | | | | |
| Left Lobule IX | 74.0 (42.3) | 0.326 | 0.3 (0.6) | 0.869 | 0.2 (0.5) | 0.948 |
| Right Lobule IX | 57.1 (35.9) | 0.326 | 0.1 (0.5) | 0.917 | 0.1 (0.4) | 0.948 |
| Vermis VI | 40.6 (17.5) | 0.261 | 0.0 (0.2) | 0.917 | 0.1 (0.2) | 0.948 |
| Vermis VII | 6.8 (12.1) | 0.688 | 0.0 (0.2) | 0.917 | 0.0 (0.1) | 0.948 |
| Vermis VIII | 24.2 (22.9) | 0.465 | -0.2 (0.3) | 0.869 | 0.0 (0.3) | 0.948 |
| Vermis IX | 17.3 (12.7) | 0.361 | 0.2 (0.2) | 0.838 | 0.0 (0.1) | 0.948 |
| Vermis X | 4.6 (4.6) | 0.465 | 0.0 (0.1) | 0.869 | 0.0 (0.1) | 0.948 |
| Total Gray Matter | 3472.8 (2240.7) | 0.326 | 41.8 (28.3) | 0.838 | 43.0 (23.7) | 0.627 |
| Cortical Volume | 919.4 (2271.2) | 0.688 | 33.1 (28.4) | 0.838 | 37.4 (23.8) | 0.627 |
| Cerebral White Matter | 751.0 (1678.5) | 0.688 | 11.2 (21.6) | 0.869 | 20.0 (18.2) | 0.931 |
| Left Hippocampus | 20.8 (23.2) | 0.445 | 0.1 (0.3) | 0.821 | 0.0 (0.3) | 0.902 |
| Right Hippocampus | 33.5 (25.7) | 0.445 | 0.1 (0.3) | 0.821 | 0.1 (0.3) | 0.902 |

| | E | | T | | P | |
|---|---|---|---|---|---|---|
| **Region** | *b (SE)* | *p* | *b (SE)* | *p* | *b (SE)* | *p* |
| Left Amygdala | 16.5 (19.2) | 0.445 | 0.2 (0.3) | 0.821 | -0.1 (0.2) | 0.902 |
| Right Amygdala | 13.1 (15.7) | 0.445 | 0.1 (0.2) | 0.821 | 0.1 (0.2) | 0.902 |
| Left Thalamus | 35.6 (46.1) | 0.445 | 0.6 (0.6) | 0.821 | 0.6 (0.5) | 0.666 |
| Right Thalamus | 45.6 (59.1) | 0.445 | -0.3 (0.7) | 0.821 | 0.8 (0.6) | 0.666 |

*Note.* Change-score models, outliers removed (11 subjects; 3*SD* within sex and timepoint). Full sample includes sex as a covariate; sex-stratified sections fit within each sex. E = 17β-estradiol; T = testosterone; P = progesterone; *b* = unstandardized coefficient; *SE* = standard error. FDR within family (exploratory: Lobule IX, vermis, global together; limbic separate). **p*_FDR < .05.

**Table S9.** Estradiol robustness checks.

| | With outliers | | Without outliers | | With outliers | | Without outliers | | With outliers | | Without outliers | |
|---|---|---|---|---|---|---|---|---|---|---|---|---|
| | Proportional scaling | | | | Within-sex z-score | | | | Global cerebellar control | | | |
| **Region** | *b (SE)* | *p* | *b (SE)* | *p* | *b (SE)* | *p* | *b (SE)* | *p* | *b (SE)* | *p* | *b (SE)* | *p* |
| Left Crus I | **0.2 (0.1)** | **0.004*** | **0.2 (0.1)** | **0.023*** | **0.1 (0.1)** | **0.015*** | **0.2 (0.1)** | **0.026*** | 84.9 (70.3) | 0.231 | 84.0 (107.0) | 0.435 |
| Right Crus I | **0.2 (0.1)** | **0.006*** | **0.2 (0.1)** | **0.018*** | **0.1 (0.1)** | **0.045*** | **0.2 (0.1)** | **0.041*** | 46.5 (65.6) | 0.480 | 55.9 (99.4) | 0.576 |
| Left Crus II | 0.0 (0.1) | 0.491 | 0.0 (0.1) | 0.874 | 0.1 (0.1) | 0.506 | 0.0 (0.1) | 0.867 | **-166.3 (71.1)** | **0.022*** | **-300.8 (105.0)** | **0.005*** |
| Right Crus II | 0.1 (0.1) | 0.044* | 0.2 (0.1) | 0.062 | 0.2 (0.1) | 0.051 | 0.2 (0.1) | 0.087 | -34.7 (72.2) | 0.632 | -33.6 (113.7) | 0.768 |
| Left Lobule V | — | — | — | — | — | — | — | — | -19.8 (29.7) | 0.508 | -65.4 (46.4) | 0.163 |
| Right Lobule V | — | — | — | — | — | — | — | — | -8.1 (29.0) | 0.780 | 38.2 (44.4) | 0.392 |
| Left Lobule VI | **0.1 (0.0)** | **0.026*** | **0.1 (0.0)** | **0.011*** | 0.1 (0.0) | **0.030*** | **0.2 (0.1)** | **0.011*** | 13.9 (36.8) | 0.706 | 61.9 (56.6) | 0.278 |
| Right Lobule VI | **0.1 (0.0)** | **0.001*** | **0.2 (0.1)** | **0.002*** | **0.2 (0.1)** | **0.001*** | **0.3 (0.1)** | **0.001*** | 76.4 (44.1) | 0.087 | 118.9 (68.6) | 0.087 |

*Note.* All values are the estradiol × sex interaction term (ΔEstradiol:SexMale) from change-score models. E = 17β-estradiol; *b* = unstandardized coefficient; *SE* = standard error. Proportional scaling expresses volumes as [(volume/estimated total intracranial volume) × 1000] (no estimated total intracranial volume as covariate); within-sex z-score standardizes volumes within each sex; global cerebellar control adds Δtotal cerebellar volume as a covariate. Lobule V was assessed only under global cerebellar control and is shown as (—) for the scaling and z-score checks. Prediction of Year 1 volume controls for baseline volume throughout. *$p < .05$.

**Table S10.** Regression-to-the-mean sensitivity with outliers.

| | | Baseline Hormone | | | | | | ΔHormone (change) | | | | | |
|---|---|---|---|---|---|---|---|---|---|---|---|---|---|
| | | E | | T | | P | | E | | T | | P | |
| **Sample** | **Region** | *b (SE)* | *p* | *b (SE)* | *p* | *b (SE)* | *p* | *b (SE)* | *p* | *b (SE)* | *p* | *b (SE)* | *p* |
| **Full sample** | Left Crus I | -15.9 (67.0) | 0.813 | -0.6 (1.0) | 0.502 | -0.9 (0.5) | 0.086 | -39.2 (52.6) | 0.994 | -0.5 (1.1) | 0.834 | -0.1 (0.6) | 0.820 |
| | Right Crus I | 87.3 (68.5) | 0.206 | 0.2 (1.0) | 0.840 | 0.2 (0.6) | 0.682 | 0.4 (53.3) | 0.994 | 0.2 (1.2) | 0.834 | 0.2 (0.6) | 0.820 |
| | Left Crus II | 85.1 (72.6) | 0.245 | -0.3 (1.1) | 0.805 | 0.5 (0.6) | 0.356 | 7.2 (55.5) | 0.994 | 0.5 (1.2) | 0.834 | 1.4 (0.7) | 0.124 |
| | Right Crus II | 38.6 (78.3) | 0.623 | -0.3 (1.1) | 0.802 | -0.2 (0.6) | 0.717 | -21.5 (60.0) | 0.994 | 0.3 (1.3) | 0.834 | 1.3 (0.7) | 0.160 |
| | Left Lobule V | -6.5 (23.3) | 0.779 | -0.5 (0.3) | 0.172 | -0.2 (0.2) | 0.243 | 0.0 (17.8) | 0.999 | -0.5 (0.4) | 0.227 | -0.2 (0.2) | 0.556 |
| | Right Lobule V | 10.2 (25.6) | 0.691 | -0.3 (0.4) | 0.387 | -0.2 (0.2) | 0.432 | -12.8 (19.9) | 0.885 | -0.5 (0.4) | 0.297 | 0.0 (0.2) | 0.911 |
| | Left Lobule VI | 63.9 (33.2) | 0.058 | 0.9 (0.5) | 0.074 | 0.0 (0.3) | 0.894 | 11.0 (25.3) | 0.885 | 0.8 (0.6) | 0.227 | 0.4 (0.3) | 0.515 |
| | Right Lobule VI | 96.9 (41.6) | 0.022 | 0.6 (0.6) | 0.322 | 0.1 (0.3) | 0.823 | 36.3 (31.9) | 0.885 | 1.5 (0.7) | 0.131 | 0.9 (0.4) | 0.086 |
| | Left Hippocampus | 2.6 (14.2) | 0.853 | 0.1 (0.2) | 0.593 | -0.1 (0.1) | 0.196 | 2.7 (11.0) | 0.959 | 0.0 (0.2) | 0.936 | 0.0 (0.1) | 0.885 |
| | Right Hippocampus | -11.1 (14.8) | 0.457 | 0.0 (0.2) | 0.989 | -0.2 (0.1) | 0.091 | -6.2 (11.4) | 0.959 | 0.0 (0.3) | 0.936 | 0.1 (0.1) | 0.885 |
| | Left Amygdala | 5.8 (12.2) | 0.634 | 0.3 (0.2) | 0.074 | 0.0 (0.1) | 0.810 | 0.5 (9.5) | 0.959 | 0.4 (0.2) | 0.409 | 0.0 (0.1) | 0.885 |
| | Right Amygdala | 4.5 (10.1) | 0.656 | 0.2 (0.1) | 0.089 | 0.0 (0.1) | 0.980 | -8.3 (7.5) | 0.959 | 0.2 (0.2) | 0.770 | -0.1 (0.1) | 0.885 |
| | Left Thalamus | 6.0 (29.9) | 0.842 | -0.4 (0.4) | 0.364 | -0.1 (0.2) | 0.638 | 22.0 (22.9) | 0.959 | -0.3 (0.5) | 0.936 | -0.1 (0.3) | 0.885 |
| | Right Thalamus | -6.5 (35.3) | 0.853 | 0.1 (0.5) | 0.809 | -0.1 (0.3) | 0.672 | 5.4 (26.8) | 0.959 | 0.0 (0.6) | 0.936 | 0.1 (0.3) | 0.885 |
| **Females** | Left Crus I | -90.9 (85.2) | 0.293 | -1.6 (1.3) | 0.234 | -0.4 (0.6) | 0.505 | -158.9 (65.0) | 0.075 | -4.7 (1.8) | 0.051 | -0.9 (0.7) | 0.251 |
| | Right Crus I | 112.8 (101.4) | 0.272 | 2.0 (1.6) | 0.235 | 0.7 (0.7) | 0.296 | -61.4 (76.2) | 0.566 | -0.9 (2.2) | 0.927 | -0.1 (0.8) | 0.931 |
| | Left Crus II | 113.5 (125.4) | 0.371 | -0.3 (2.0) | 0.887 | 1.1 (0.8) | 0.180 | 10.5 (95.2) | 0.913 | 1.5 (2.7) | 0.927 | 1.7 (1.0) | 0.203 |
| | Right Crus II | 39.9 (115.3) | 0.731 | -1.0 (2.0) | 0.618 | 0.0 (0.8) | 0.961 | -113.8 (87.5) | 0.402 | -0.1 (2.6) | 0.962 | 1.5 (0.9) | 0.203 |
| | Left Lobule V | 5.4 (33.5) | 0.874 | -0.6 (0.5) | 0.222 | -0.2 (0.2) | 0.276 | -3.6 (25.6) | 0.888 | -1.2 (0.7) | 0.198 | -0.3 (0.3) | 0.489 |
| | Right Lobule V | 27.4 (36.4) | 0.455 | -0.1 (0.5) | 0.788 | -0.1 (0.2) | 0.794 | -17.6 (27.2) | 0.695 | **-2.2 (0.7)** | **0.015*** | -0.3 (0.3) | 0.489 |
| | Left Lobule VI | 12.2 (48.6) | 0.803 | 0.9 (0.8) | 0.231 | -0.1 (0.3) | 0.857 | -40.0 (36.2) | 0.611 | 0.5 (1.1) | 0.652 | 0.0 (0.4) | 0.955 |
| | Right Lobule VI | 88.7 (50.8) | 0.088 | 1.0 (0.9) | 0.259 | 0.0 (0.3) | 0.915 | -39.4 (38.0) | 0.611 | 1.2 (1.2) | 0.411 | 0.4 (0.4) | 0.489 |
| | Left Hippocampus | -26.4 (18.8) | 0.169 | -0.2 (0.3) | 0.558 | -0.2 (0.1) | 0.235 | -11.9 (14.6) | 0.842 | -0.1 (0.4) | 0.986 | -0.1 (0.2) | 0.847 |
| | Right Hippocampus | -19.1 (18.5) | 0.308 | -0.1 (0.3) | 0.672 | -0.1 (0.1) | 0.253 | -19.8 (14.0) | 0.499 | -0.2 (0.4) | 0.986 | 0.0 (0.1) | 0.893 |
| | Left Amygdala | 27.7 (16.1) | 0.094 | 0.4 (0.3) | 0.114 | 0.1 (0.1) | 0.575 | 4.0 (12.8) | 0.909 | 0.2 (0.4) | 0.986 | 0.0 (0.1) | 0.847 |
| | Right Amygdala | -4.9 (13.5) | 0.718 | 0.0 (0.2) | 0.944 | 0.0 (0.1) | 0.971 | -17.3 (10.2) | 0.499 | 0.0 (0.3) | 0.986 | -0.2 (0.1) | 0.761 |
| | Left Thalamus | -45.8 (46.0) | 0.326 | -1.1 (0.8) | 0.163 | -0.3 (0.3) | 0.273 | -3.0 (34.8) | 0.931 | -0.8 (1.0) | 0.986 | -0.4 (0.4) | 0.847 |
| | Right Thalamus | -70.3 (47.7) | 0.148 | -0.9 (0.8) | 0.291 | -0.7 (0.3) | 0.047 | -13.5 (35.7) | 0.909 | 0.2 (1.1) | 0.986 | -0.2 (0.4) | 0.847 |
| **Males** | Left Crus I | 57.7 (91.4) | 0.532 | -0.5 (1.2) | 0.697 | -1.2 (1.3) | 0.381 | 144.4 (76.7) | 0.163 | 0.2 (1.4) | 0.875 | 0.9 (1.1) | 0.672 |
| | Right Crus I | 89.5 (87.7) | 0.314 | -0.9 (1.2) | 0.471 | -0.3 (1.3) | 0.805 | 129.8 (72.5) | 0.163 | -0.3 (1.3) | 0.875 | 0.6 (1.0) | 0.672 |
| | Left Crus II | 71.2 (73.8) | 0.341 | -0.8 (1.0) | 0.421 | -0.3 (1.0) | 0.791 | 65.4 (60.2) | 0.284 | -0.2 (1.1) | 0.875 | 0.4 (0.8) | 0.672 |
| | Right Crus II | 3.5 (111.4) | 0.975 | -0.1 (1.5) | 0.956 | -0.9 (1.5) | 0.547 | 117.7 (90.7) | 0.270 | 0.5 (1.7) | 0.875 | 0.9 (1.3) | 0.672 |
| | Left Lobule V | -21.0 (36.3) | 0.566 | -0.3 (0.5) | 0.472 | -0.3 (0.5) | 0.494 | 13.0 (29.3) | 0.660 | -0.3 (0.5) | 0.734 | 0.1 (0.4) | 0.857 |
| | Right Lobule V | -7.8 (38.6) | 0.840 | -0.4 (0.5) | 0.399 | -0.8 (0.5) | 0.132 | 18.4 (31.4) | 0.660 | 0.0 (0.5) | 0.953 | 0.5 (0.4) | 0.335 |
| | Left Lobule VI | 109.0 (41.9) | 0.013 | 0.6 (0.6) | 0.302 | 0.8 (0.6) | 0.229 | **85.2 (34.1)** | **0.034*** | 0.7 (0.7) | 0.693 | 0.9 (0.5) | 0.178 |
| | Right Lobule VI | 85.4 (60.2) | 0.164 | 0.1 (0.9) | 0.872 | 0.7 (0.9) | 0.400 | **166.2 (49.1)** | **0.007**** | 1.3 (1.0) | 0.683 | 1.6 (0.7) | 0.142 |
| | Left Hippocampus | 46.6 (21.3) | 0.035 | 0.3 (0.3) | 0.232 | 0.1 (0.3) | 0.774 | 17.7 (16.3) | 0.679 | 0.1 (0.3) | 0.896 | -0.1 (0.2) | 0.960 |
| | Right Hippocampus | 3.7 (23.9) | 0.877 | 0.1 (0.3) | 0.717 | -0.1 (0.3) | 0.814 | 20.1 (18.7) | 0.679 | 0.1 (0.3) | 0.896 | 0.0 (0.3) | 0.960 |
| | Left Amygdala | -9.8 (19.4) | 0.616 | 0.2 (0.2) | 0.342 | 0.1 (0.2) | 0.547 | 8.6 (14.5) | 0.693 | 0.4 (0.3) | 0.543 | 0.0 (0.2) | 0.960 |
| | Right Amygdala | 23.5 (17.7) | 0.192 | 0.4 (0.2) | 0.057 | 0.4 (0.2) | 0.049 | 6.8 (12.0) | 0.693 | 0.3 (0.2) | 0.543 | 0.0 (0.2) | 0.960 |
| | Left Thalamus | 48.5 (41.7) | 0.252 | 0.0 (0.5) | 0.979 | 0.3 (0.5) | 0.639 | 32.8 (33.9) | 0.679 | 0.1 (0.6) | 0.896 | 0.5 (0.4) | 0.856 |
| | Right Thalamus | 51.9 (54.7) | 0.349 | 0.7 (0.7) | 0.277 | 1.1 (0.7) | 0.098 | 8.8 (44.1) | 0.844 | 0.2 (0.7) | 0.896 | 0.6 (0.5) | 0.856 |

*Note.* Year 1 volume ~ Baseline volume + ΔHormone + baseline hormone + covariates. Baseline hormone = coefficient for the baseline hormone term; Δ hormone = coefficient for the change term. Full sample with outliers retained. E = 17β-estradiol; T = testosterone; P = progesterone; *b* = unstandardized coefficient; *SE* = standard error. Δ-hormone *p*-values FDR-corrected within family (Crus I/II; Lobule V/VI; limbic); baseline *p*-values shown uncorrected. **p*_FDR < .05, ***p*_FDR < .01.

**Table S11.** Regression-to-the-mean sensitivity without outliers.

| | | Baseline Hormone | | | | | | ΔHormone (change) | | | | | |
|---|---|---|---|---|---|---|---|---|---|---|---|---|---|
| | | E | | T | | P | | E | | T | | P | |
| **Sample** | **Region** | *b (SE)* | *p* | *b (SE)* | *p* | *b (SE)* | *p* | *b (SE)* | *p* | *b (SE)* | *p* | *b (SE)* | *p* |
| **Full sample** | Left Crus I | -89.3 (78.2) | 0.257 | -0.5 (1.0) | 0.654 | -1.2 (0.8) | 0.104 | -71.3 (82.3) | 0.389 | -1.5 (1.2) | 0.769 | -1.2 (0.7) | 0.385 |
| | Right Crus I | 71.7 (83.2) | 0.391 | -0.6 (1.1) | 0.566 | -0.2 (0.8) | 0.769 | 88.7 (87.7) | 0.389 | -0.1 (1.2) | 0.980 | -0.7 (0.8) | 0.387 |
| | Left Crus II | 145.8 (87.9) | 0.101 | -0.2 (1.2) | 0.877 | 0.7 (0.9) | 0.449 | 119.3 (93.5) | 0.389 | 0.0 (1.3) | 0.980 | 0.9 (0.9) | 0.385 |
| | Right Crus II | 110.8 (97.8) | 0.261 | -0.5 (1.3) | 0.698 | 0.3 (1.0) | 0.754 | 130.6 (103.7) | 0.389 | 0.4 (1.5) | 0.980 | 1.1 (1.0) | 0.385 |
| | Left Lobule V | -6.6 (29.8) | 0.826 | -0.3 (0.4) | 0.422 | -0.1 (0.3) | 0.803 | 3.2 (31.3) | 0.920 | -0.7 (0.4) | 0.201 | -0.4 (0.3) | 0.276 |
| | Right Lobule V | 7.4 (32.3) | 0.820 | -0.2 (0.4) | 0.616 | -0.2 (0.3) | 0.479 | -9.6 (33.9) | 0.920 | -0.8 (0.5) | 0.201 | -0.3 (0.3) | 0.465 |
| | Left Lobule VI | 40.1 (40.2) | 0.322 | 0.7 (0.5) | 0.205 | -0.2 (0.4) | 0.576 | 15.7 (42.1) | 0.920 | 0.5 (0.6) | 0.405 | 0.2 (0.4) | 0.529 |
| | Right Lobule VI | 97.4 (51.5) | 0.063 | 0.5 (0.7) | 0.490 | 0.0 (0.5) | 0.980 | 49.5 (54.1) | 0.920 | 1.1 (0.8) | 0.201 | 0.9 (0.5) | 0.276 |
| | Left Hippocampus | 10.6 (17.1) | 0.537 | 0.0 (0.2) | 0.847 | 0.0 (0.2) | 0.957 | 27.5 (17.9) | 0.256 | 0.0 (0.3) | 0.965 | 0.0 (0.2) | 0.959 |
| | Right Hippocampus | -4.5 (18.0) | 0.802 | 0.0 (0.2) | 0.943 | -0.2 (0.2) | 0.183 | 14.1 (18.9) | 0.688 | 0.0 (0.3) | 0.965 | 0.1 (0.2) | 0.959 |
| | Left Amygdala | 6.5 (15.0) | 0.669 | 0.4 (0.2) | 0.047 | 0.0 (0.1) | 0.737 | 4.5 (15.7) | 0.932 | 0.2 (0.2) | 0.965 | -0.1 (0.1) | 0.959 |
| | Right Amygdala | 5.5 (12.0) | 0.651 | 0.2 (0.2) | 0.200 | 0.1 (0.1) | 0.634 | -0.7 (12.5) | 0.953 | 0.1 (0.2) | 0.965 | 0.0 (0.1) | 0.959 |
| | Left Thalamus | 9.1 (37.1) | 0.807 | -0.9 (0.5) | 0.080 | 0.0 (0.4) | 0.995 | 74.9 (39.0) | 0.253 | -0.1 (0.5) | 0.965 | -0.1 (0.3) | 0.959 |
| | Right Thalamus | 8.4 (42.9) | 0.845 | -0.3 (0.6) | 0.644 | 0.1 (0.4) | 0.802 | 78.3 (44.7) | 0.253 | -0.1 (0.6) | 0.965 | 0.3 (0.4) | 0.959 |
| **Females** | Left Crus I | -115.1 (95.4) | 0.235 | -1.0 (1.4) | 0.471 | -0.3 (0.9) | 0.732 | -231.1 (110.3) | 0.172 | -4.9 (1.9) | 0.053 | -2.1 (1.0) | 0.149 |
| | Right Crus I | 119.6 (113.9) | 0.300 | 2.1 (1.7) | 0.223 | 0.7 (1.0) | 0.469 | -31.7 (129.4) | 0.884 | -0.9 (2.3) | 0.879 | -1.6 (1.1) | 0.213 |
| | Left Crus II | 180.5 (133.5) | 0.185 | 0.2 (2.1) | 0.910 | 1.8 (1.2) | 0.135 | 178.0 (159.8) | 0.545 | 1.2 (2.8) | 0.879 | 1.3 (1.3) | 0.346 |
| | Right Crus II | 99.0 (128.1) | 0.445 | -0.2 (2.0) | 0.923 | 1.4 (1.2) | 0.222 | -22.5 (152.4) | 0.884 | 0.4 (2.7) | 0.879 | 1.9 (1.3) | 0.213 |
| | Left Lobule V | 27.0 (37.0) | 0.471 | -0.4 (0.5) | 0.497 | 0.0 (0.3) | 0.932 | 21.3 (42.8) | 0.621 | -1.1 (0.7) | 0.272 | -0.6 (0.4) | 0.193 |
| | Right Lobule V | 15.7 (40.7) | 0.702 | 0.1 (0.6) | 0.865 | 0.0 (0.4) | 0.894 | -59.5 (46.7) | 0.280 | **-2.3 (0.7)** | **0.015*** | -0.7 (0.4) | 0.193 |
| | Left Lobule VI | -13.5 (53.6) | 0.803 | 1.2 (0.8) | 0.134 | -0.2 (0.5) | 0.740 | -98.6 (61.3) | 0.280 | 0.2 (1.1) | 0.823 | -0.5 (0.6) | 0.520 |
| | Right Lobule VI | 73.6 (57.5) | 0.208 | 1.1 (0.9) | 0.241 | 0.3 (0.5) | 0.544 | -84.8 (66.1) | 0.280 | 1.5 (1.2) | 0.307 | 0.3 (0.6) | 0.614 |
| | Left Hippocampus | -15.4 (20.2) | 0.450 | 0.0 (0.3) | 0.899 | 0.1 (0.2) | 0.632 | 0.9 (23.5) | 0.969 | -0.1 (0.4) | 0.900 | 0.0 (0.2) | 0.954 |
| | Right Hippocampus | -12.9 (20.4) | 0.532 | 0.0 (0.3) | 0.891 | -0.1 (0.2) | 0.688 | -10.5 (23.6) | 0.969 | -0.2 (0.4) | 0.900 | 0.1 (0.2) | 0.954 |
| | Left Amygdala | 25.8 (17.7) | 0.154 | 0.6 (0.3) | 0.038 | 0.1 (0.2) | 0.409 | -1.7 (20.8) | 0.969 | 0.0 (0.4) | 0.900 | -0.1 (0.2) | 0.954 |
| | Right Amygdala | -5.7 (15.0) | 0.704 | 0.1 (0.2) | 0.736 | 0.1 (0.1) | 0.678 | -23.6 (17.3) | 0.360 | -0.1 (0.3) | 0.900 | -0.1 (0.1) | 0.954 |
| | Left Thalamus | 9.7 (49.8) | 0.846 | -0.7 (0.8) | 0.400 | -0.1 (0.5) | 0.857 | 102.6 (57.4) | 0.360 | -0.8 (1.1) | 0.900 | -0.4 (0.5) | 0.954 |
| | Right Thalamus | -12.3 (51.4) | 0.812 | -0.4 (0.8) | 0.664 | -0.4 (0.6) | 0.456 | 82.4 (58.7) | 0.360 | 0.4 (1.1) | 0.900 | 0.1 (0.5) | 0.954 |
| **Males** | Left Crus I | -33.2 (123.6) | 0.790 | -1.6 (1.5) | 0.279 | -3.2 (1.5) | 0.034 | 107.5 (114.3) | 0.354 | -0.4 (1.4) | 0.976 | -1.0 (1.1) | 0.991 |
| | Right Crus I | 24.5 (122.5) | 0.843 | -4.0 (1.3) | 0.004 | -3.1 (1.5) | 0.046 | 210.6 (113.1) | 0.144 | 0.0 (1.2) | 0.976 | 0.0 (1.1) | 0.991 |
| | Left Crus II | 115.4 (110.2) | 0.303 | -1.2 (1.3) | 0.371 | -1.3 (1.4) | 0.355 | 108.6 (100.6) | 0.354 | -0.7 (1.3) | 0.976 | -0.3 (1.0) | 0.991 |
| | Right Crus II | 68.0 (160.6) | 0.675 | -1.1 (2.0) | 0.608 | -2.9 (2.1) | 0.180 | 285.5 (146.3) | 0.144 | 0.2 (1.9) | 0.976 | 0.2 (1.6) | 0.991 |
| | Left Lobule V | -66.3 (53.0) | 0.220 | -0.3 (0.6) | 0.607 | -0.5 (0.7) | 0.415 | -25.6 (49.6) | 0.610 | -0.5 (0.6) | 0.620 | -0.3 (0.5) | 0.836 |
| | Right Lobule V | -25.5 (52.3) | 0.630 | -0.9 (0.6) | 0.160 | -1.4 (0.6) | 0.029 | 32.2 (48.0) | 0.610 | -0.1 (0.6) | 0.906 | 0.1 (0.4) | 0.852 |
| | Left Lobule VI | 111.1 (58.0) | 0.064 | -0.1 (0.7) | 0.869 | -0.5 (0.8) | 0.551 | **136.1 (53.0)** | **0.030*** | 0.5 (0.7) | 0.620 | 0.9 (0.5) | 0.237 |
| | Right Lobule VI | 123.7 (85.6) | 0.158 | -0.6 (1.1) | 0.596 | -0.8 (1.1) | 0.480 | **219.4 (80.6)** | **0.030*** | 1.0 (1.1) | 0.620 | 1.3 (0.8) | 0.237 |
| | Left Hippocampus | 60.6 (29.4) | 0.048 | 0.1 (0.4) | 0.729 | -0.2 (0.4) | 0.688 | 50.9 (26.0) | 0.357 | 0.2 (0.3) | 0.890 | 0.0 (0.3) | 0.922 |
| | Right Hippocampus | 9.9 (34.7) | 0.777 | 0.0 (0.4) | 0.920 | -0.6 (0.4) | 0.190 | 40.4 (31.0) | 0.404 | 0.1 (0.4) | 0.890 | 0.0 (0.3) | 0.922 |
| | Left Amygdala | -27.1 (25.7) | 0.300 | 0.2 (0.3) | 0.541 | -0.2 (0.3) | 0.617 | 2.7 (22.8) | 0.907 | 0.3 (0.3) | 0.890 | -0.1 (0.2) | 0.922 |
| | Right Amygdala | 31.3 (21.8) | 0.160 | 0.3 (0.2) | 0.233 | 0.2 (0.3) | 0.393 | 27.1 (18.5) | 0.404 | 0.2 (0.2) | 0.890 | 0.1 (0.2) | 0.922 |
| | Left Thalamus | 4.9 (61.1) | 0.937 | -1.1 (0.6) | 0.084 | -0.6 (0.7) | 0.398 | 39.2 (55.6) | 0.583 | 0.2 (0.6) | 0.890 | 0.5 (0.5) | 0.915 |
| | Right Thalamus | 29.9 (77.9) | 0.704 | 0.3 (0.9) | 0.754 | 0.4 (0.9) | 0.613 | 56.5 (71.3) | 0.583 | -0.1 (0.8) | 0.890 | 0.9 (0.6) | 0.915 |

*Note.* Regression-to-the-mean sensitivity: Year 1 volume ~ Baseline volume + ΔHormone + baseline hormone + covariates. Baseline hormone = coefficient for the baseline hormone term; Δ hormone = coefficient for the change term. Outliers removed (11 subjects; 3*SD* within sex and timepoint). E = 17β-estradiol; T = testosterone; P = progesterone; *b* = unstandardized coefficient; *SE* = standard error. Δ-hormone *p*-values FDR-corrected within family (Crus I/II; Lobule V/VI; limbic); baseline *p*-values shown uncorrected. *p_FDR < .05*.

**Table S12.** Between-session interval sensitivity check with outliers.

| | | E | | | | T | | | | P | | | |
|---|---|---|---|---|---|---|---|---|---|---|---|---|---|
| | | Scan-to-scan | | Saliva-to-saliva | | Scan-to-scan | | Saliva-to-saliva | | Scan-to-scan | | Saliva-to-saliva | |
| **Sample** | **Region** | *b (SE)* | *p* | *b (SE)* | *p* | *b (SE)* | *p* | *b (SE)* | *p* | *b (SE)* | *p* | *b (SE)* | *p* |
| **Full sample** | Left Crus I | -19.6 (51.8) | 0.744 | 14.6 (49.4) | 0.948 | 0.2 (0.8) | 0.823 | 0.0 (0.8) | 0.957 | 0.2 (0.6) | 0.715 | 0.1 (0.6) | 0.836 |
| | Right Crus I | -16.8 (51.5) | 0.744 | -3.3 (49.5) | 0.948 | 0.2 (0.8) | 0.823 | 0.1 (0.8) | 0.957 | 0.2 (0.6) | 0.715 | 0.1 (0.6) | 0.836 |
| | Left Crus II | -22.9 (56.0) | 0.744 | -6.9 (54.2) | 0.948 | 0.6 (0.9) | 0.823 | 0.6 (0.9) | 0.957 | 1.3 (0.6) | 0.199 | 1.2 (0.7) | 0.241 |
| | Right Crus II | -35.6 (61.2) | 0.744 | -14.8 (59.1) | 0.948 | 0.6 (1.0) | 0.823 | 0.5 (1.0) | 0.957 | 1.2 (0.7) | 0.199 | 1.1 (0.7) | 0.241 |
| | Left Lobule V | -1.5 (17.8) | 0.934 | 2.1 (17.3) | 0.961 | -0.1 (0.3) | 0.839 | -0.1 (0.3) | 0.852 | 0.0 (0.2) | 0.869 | 0.0 (0.2) | 0.885 |
| | Right Lobule V | -13.7 (19.4) | 0.934 | -7.1 (18.6) | 0.961 | -0.1 (0.3) | 0.839 | -0.2 (0.3) | 0.852 | 0.1 (0.2) | 0.869 | 0.0 (0.2) | 0.885 |
| | Left Lobule VI | -4.1 (25.3) | 0.934 | 1.2 (24.4) | 0.961 | 0.1 (0.4) | 0.839 | 0.1 (0.4) | 0.852 | 0.4 (0.3) | 0.381 | 0.4 (0.3) | 0.444 |
| | Right Lobule VI | 4.2 (32.1) | 0.934 | 13.9 (31.1) | 0.961 | 1.0 (0.5) | 0.174 | 1.0 (0.5) | 0.196 | 0.8 (0.3) | 0.071 | 0.8 (0.4) | 0.088 |
| | Left Hippocampus | 1.9 (11.0) | 0.933 | 9.9 (10.7) | 0.719 | -0.2 (0.2) | 0.902 | -0.2 (0.2) | 0.950 | 0.0 (0.1) | 0.980 | 0.0 (0.1) | 0.952 |
| | Right Hippocampus | -5.7 (11.3) | 0.933 | -0.2 (10.9) | 0.986 | 0.0 (0.2) | 0.902 | 0.0 (0.2) | 0.950 | 0.1 (0.1) | 0.980 | 0.1 (0.1) | 0.952 |
| | Left Amygdala | 1.1 (8.9) | 0.933 | 3.5 (8.5) | 0.821 | 0.1 (0.1) | 0.902 | 0.1 (0.1) | 0.950 | 0.0 (0.1) | 0.980 | 0.0 (0.1) | 0.952 |
| | Right Amygdala | -13.9 (7.2) | 0.334 | -8.9 (7.1) | 0.719 | 0.0 (0.1) | 0.902 | 0.0 (0.1) | 0.950 | -0.1 (0.1) | 0.980 | -0.1 (0.1) | 0.952 |
| | Left Thalamus | 17.0 (22.4) | 0.933 | 25.0 (21.7) | 0.719 | 0.0 (0.3) | 0.902 | 0.0 (0.3) | 0.950 | -0.1 (0.3) | 0.980 | -0.1 (0.3) | 0.952 |
| | Right Thalamus | 2.2 (26.3) | 0.933 | 15.5 (25.4) | 0.815 | -0.1 (0.4) | 0.902 | -0.2 (0.4) | 0.950 | 0.1 (0.3) | 0.980 | 0.1 (0.3) | 0.952 |
| **Females** | Left Crus I | -136.3 (56.9) | 0.085 | -116.8 (55.9) | 0.139 | -3.5 (1.5) | 0.097 | -3.6 (1.5) | 0.085 | -0.8 (0.7) | 0.315 | -0.8 (0.7) | 0.308 |
| | Right Crus I | -126.1 (67.2) | 0.099 | -121.3 (65.1) | 0.139 | -2.4 (1.8) | 0.404 | -2.4 (1.8) | 0.408 | -0.4 (0.8) | 0.628 | -0.4 (0.8) | 0.637 |
| | Left Crus II | -59.0 (92.1) | 0.525 | -45.5 (89.1) | 0.612 | 1.2 (2.4) | 0.795 | 1.2 (2.4) | 0.852 | 1.6 (1.0) | 0.241 | 1.6 (1.0) | 0.241 |
| | Right Crus II | -152.7 (83.4) | 0.099 | -130.0 (80.9) | 0.154 | 0.6 (2.3) | 0.795 | 0.4 (2.3) | 0.852 | 1.5 (0.9) | 0.241 | 1.5 (0.9) | 0.241 |
| | Left Lobule V | -13.7 (23.2) | 0.559 | -13.3 (22.4) | 0.556 | -0.7 (0.6) | 0.526 | -0.6 (0.6) | 0.601 | -0.2 (0.3) | 0.520 | -0.2 (0.3) | 0.553 |
| | Right Lobule V | -31.2 (24.9) | 0.288 | -25.3 (24.0) | 0.398 | **-1.9 (0.6)** | **0.013*** | **-2.0 (0.6)** | **0.009**** | -0.3 (0.3) | 0.520 | -0.3 (0.3) | 0.553 |
| | Left Lobule VI | -42.2 (32.4) | 0.288 | -40.9 (31.3) | 0.397 | -0.1 (0.9) | 0.893 | -0.2 (0.9) | 0.823 | 0.1 (0.4) | 0.799 | 0.1 (0.4) | 0.813 |
| | Right Lobule VI | -73.1 (35.3) | 0.177 | -74.6 (34.1) | 0.137 | 0.5 (1.0) | 0.810 | 0.5 (1.0) | 0.823 | 0.4 (0.4) | 0.520 | 0.4 (0.4) | 0.553 |
| | Left Hippocampus | 0.1 (14.4) | 0.993 | 12.1 (14.5) | 0.808 | 0.0 (0.4) | 0.999 | 0.0 (0.4) | 0.992 | 0.0 (0.2) | 0.961 | 0.0 (0.2) | 0.981 |
| | Right Hippocampus | -17.4 (13.2) | 0.591 | -9.6 (13.1) | 0.808 | 0.0 (0.4) | 0.999 | 0.0 (0.4) | 0.992 | 0.1 (0.1) | 0.765 | 0.1 (0.2) | 0.672 |
| | Left Amygdala | 1.3 (11.5) | 0.993 | 0.3 (11.0) | 0.978 | -0.1 (0.3) | 0.999 | -0.2 (0.3) | 0.992 | -0.1 (0.1) | 0.765 | -0.1 (0.1) | 0.672 |
| | Right Amygdala | -21.1 (8.7) | 0.123 | -16.6 (8.8) | 0.405 | 0.0 (0.2) | 0.999 | 0.0 (0.2) | 0.992 | -0.2 (0.1) | 0.580 | -0.2 (0.1) | 0.660 |
| | Left Thalamus | -10.2 (29.1) | 0.993 | 13.0 (30.5) | 0.808 | 0.0 (0.8) | 0.999 | 0.1 (0.8) | 0.992 | -0.3 (0.3) | 0.765 | -0.3 (0.3) | 0.672 |
| | Right Thalamus | -8.7 (30.9) | 0.993 | 16.4 (31.8) | 0.808 | 0.8 (0.8) | 0.999 | 0.9 (0.9) | 0.992 | 0.0 (0.3) | 0.961 | 0.0 (0.4) | 0.981 |
| **Males** | Left Crus I | 204.8 (85.0) | 0.075 | **241.0 (74.6)** | **0.010*** | 1.2 (1.0) | 0.603 | 1.0 (1.0) | 0.672 | 1.5 (1.1) | 0.660 | 1.2 (1.1) | 0.764 |
| | Right Crus I | 171.4 (79.6) | 0.075 | **185.9 (72.9)** | **0.030*** | 0.9 (0.9) | 0.603 | 0.8 (0.9) | 0.672 | 1.0 (1.1) | 0.660 | 0.8 (1.1) | 0.764 |
| | Left Crus II | 52.9 (62.4) | 0.402 | 67.2 (58.8) | 0.260 | 0.4 (0.7) | 0.603 | 0.3 (0.7) | 0.672 | 0.5 (0.8) | 0.660 | 0.3 (0.8) | 0.764 |
| | Right Crus II | 144.8 (94.2) | 0.177 | 158.2 (88.7) | 0.110 | 0.6 (1.1) | 0.603 | 0.5 (1.1) | 0.672 | 0.6 (1.2) | 0.660 | 0.4 (1.3) | 0.764 |
| | Left Lobule V | 17.9 (30.4) | 0.559 | 25.8 (29.3) | 0.384 | 0.1 (0.3) | 0.707 | 0.1 (0.3) | 0.766 | 0.3 (0.4) | 0.461 | 0.3 (0.4) | 0.513 |
| | Right Lobule V | 28.1 (31.9) | 0.512 | 32.1 (30.2) | 0.384 | 0.4 (0.4) | 0.646 | 0.3 (0.4) | 0.738 | 0.6 (0.4) | 0.275 | 0.6 (0.4) | 0.343 |
| | Left Lobule VI | 78.0 (39.1) | 0.107 | 83.0 (36.9) | 0.061 | 0.2 (0.5) | 0.707 | 0.1 (0.5) | 0.766 | 0.7 (0.5) | 0.275 | 0.6 (0.5) | 0.343 |
| | Right Lobule VI | **142.3 (52.4)** | **0.039*** | **162.6 (49.2)** | **0.008**** | 1.2 (0.6) | 0.225 | 1.1 (0.6) | 0.312 | 1.5 (0.7) | 0.132 | 1.4 (0.7) | 0.231 |
| | Left Hippocampus | 11.4 (18.2) | 0.783 | 12.6 (17.4) | 0.711 | -0.2 (0.2) | 0.749 | -0.2 (0.2) | 0.696 | -0.1 (0.2) | 0.928 | -0.1 (0.2) | 0.917 |
| | Right Hippocampus | 23.7 (19.3) | 0.679 | 23.2 (18.5) | 0.593 | 0.0 (0.2) | 0.958 | 0.0 (0.2) | 0.993 | 0.0 (0.3) | 0.928 | 0.0 (0.3) | 0.933 |
| | Left Amygdala | 8.9 (14.7) | 0.783 | 14.9 (13.8) | 0.593 | 0.2 (0.2) | 0.749 | 0.2 (0.2) | 0.696 | 0.0 (0.2) | 0.928 | -0.1 (0.2) | 0.917 |
| | Right Amygdala | 1.2 (12.5) | 0.922 | 5.3 (11.9) | 0.751 | 0.0 (0.1) | 0.958 | 0.0 (0.1) | 0.993 | 0.0 (0.2) | 0.928 | -0.1 (0.2) | 0.917 |
| | Left Thalamus | 54.9 (32.8) | 0.613 | 35.3 (33.4) | 0.593 | 0.0 (0.3) | 0.958 | 0.0 (0.4) | 0.993 | 0.3 (0.4) | 0.928 | 0.4 (0.4) | 0.917 |
| | Right Thalamus | 20.8 (45.8) | 0.783 | 14.1 (44.0) | 0.751 | -0.4 (0.5) | 0.749 | -0.4 (0.5) | 0.696 | 0.3 (0.5) | 0.928 | 0.3 (0.5) | 0.917 |

*Note.* Change-score models (ΔHormone predicting ΔVolume) repeated with the between-session interval as a covariate, shown for both scan-to-scan and saliva-to-saliva intervals. Full sample with outliers retained. E = 17β-estradiol; T = testosterone; P = progesterone; *b* = unstandardized coefficient; *SE* = standard error. *p*-values FDR-corrected within family (Crus I/II; Lobule V/VI; limbic). **p*_FDR < .05, ***p*_FDR < .01.

**Table S13.** Between-session interval sensitivity check without outliers.

| | | E | | | | T | | | | P | | | |
|---|---|---|---|---|---|---|---|---|---|---|---|---|---|
| | | Scan-to-scan | | Saliva-to-saliva | | Scan-to-scan | | Saliva-to-saliva | | Scan-to-scan | | Saliva-to-saliva | |
| **Sample** | **Region** | *b (SE)* | *p* | *b (SE)* | *p* | *b (SE)* | *p* | *b (SE)* | *p* | *b (SE)* | *p* | *b (SE)* | *p* |
| **Full sample** | Left Crus I | -52.6 (75.4) | 0.955 | -5.2 (74.0) | 0.945 | -1.4 (1.1) | 0.822 | -1.4 (1.1) | 0.779 | -0.6 (0.7) | 0.489 | -0.5 (0.7) | 0.499 |
| | Right Crus I | 35.9 (76.4) | 0.955 | 47.6 (73.5) | 0.945 | 0.4 (1.1) | 0.990 | 0.3 (1.1) | 0.983 | -0.5 (0.7) | 0.489 | -0.5 (0.7) | 0.499 |
| | Left Crus II | 4.8 (84.7) | 0.955 | 11.5 (81.4) | 0.945 | 0.0 (1.2) | 0.990 | 0.0 (1.2) | 0.983 | 0.6 (0.8) | 0.489 | 0.7 (0.8) | 0.499 |
| | Right Crus II | 17.3 (95.7) | 0.955 | 33.3 (92.0) | 0.945 | 0.4 (1.4) | 0.990 | 0.4 (1.4) | 0.983 | 0.7 (0.9) | 0.489 | 0.8 (0.9) | 0.499 |
| | Left Lobule V | 2.9 (27.8) | 0.917 | 4.6 (26.7) | 0.879 | -0.4 (0.4) | 0.432 | -0.4 (0.4) | 0.421 | -0.3 (0.3) | 0.416 | -0.3 (0.3) | 0.430 |
| | Right Lobule V | -22.0 (29.8) | 0.917 | -12.2 (28.8) | 0.879 | -0.6 (0.4) | 0.391 | -0.6 (0.4) | 0.417 | -0.2 (0.3) | 0.507 | -0.2 (0.3) | 0.519 |
| | Left Lobule VI | -7.0 (37.5) | 0.917 | -5.5 (36.0) | 0.879 | 0.3 (0.5) | 0.572 | 0.3 (0.5) | 0.575 | 0.4 (0.4) | 0.416 | 0.4 (0.4) | 0.430 |
| | Right Lobule VI | -23.6 (48.0) | 0.917 | -13.4 (46.4) | 0.879 | 0.9 (0.7) | 0.391 | 0.9 (0.7) | 0.417 | 0.8 (0.4) | 0.179 | 0.9 (0.4) | 0.181 |
| | Left Hippocampus | 14.9 (16.4) | 0.445 | 25.2 (15.7) | 0.225 | 0.0 (0.2) | 0.987 | 0.0 (0.2) | 0.965 | 0.0 (0.2) | 0.842 | 0.0 (0.2) | 0.848 |
| | Right Hippocampus | 15.1 (16.8) | 0.445 | 18.2 (16.1) | 0.395 | 0.0 (0.2) | 0.987 | 0.0 (0.2) | 0.965 | 0.2 (0.2) | 0.842 | 0.2 (0.2) | 0.848 |
| | Left Amygdala | 3.7 (13.8) | 0.790 | 5.1 (13.1) | 0.699 | 0.0 (0.2) | 0.987 | 0.0 (0.2) | 0.965 | -0.1 (0.1) | 0.842 | -0.1 (0.1) | 0.848 |
| | Right Amygdala | -11.3 (10.8) | 0.445 | -4.9 (10.6) | 0.699 | 0.0 (0.2) | 0.987 | 0.0 (0.2) | 0.965 | 0.0 (0.1) | 0.842 | 0.0 (0.1) | 0.848 |
| | Left Thalamus | 63.4 (34.1) | 0.310 | 71.2 (32.8) | 0.118 | 0.3 (0.5) | 0.987 | 0.3 (0.5) | 0.965 | -0.1 (0.3) | 0.842 | -0.1 (0.3) | 0.848 |
| | Right Thalamus | 64.7 (39.2) | 0.310 | 79.2 (37.7) | 0.118 | 0.0 (0.6) | 0.987 | 0.0 (0.6) | 0.965 | 0.3 (0.4) | 0.842 | 0.3 (0.4) | 0.848 |
| **Females** | Left Crus I | -185.7 (93.4) | 0.216 | -158.4 (89.9) | 0.344 | **-4.2 (1.6)** | **0.048*** | **-4.3 (1.6)** | **0.043*** | -1.9 (0.9) | 0.104 | -1.9 (0.9) | 0.112 |
| | Right Crus I | -122.6 (109.4) | 0.359 | -118.2 (103.9) | 0.350 | -2.4 (1.9) | 0.438 | -2.4 (2.0) | 0.458 | -2.0 (1.0) | 0.104 | -2.0 (1.0) | 0.112 |
| | Left Crus II | -22.8 (142.3) | 0.873 | -33.0 (135.4) | 0.809 | 0.3 (2.5) | 0.914 | 0.2 (2.6) | 0.939 | 1.0 (1.4) | 0.452 | 1.0 (1.4) | 0.456 |
| | Right Crus II | -168.6 (133.7) | 0.359 | -156.1 (127.1) | 0.350 | 0.3 (2.5) | 0.914 | 0.2 (2.5) | 0.939 | 1.5 (1.3) | 0.337 | 1.5 (1.3) | 0.351 |
| | Left Lobule V | 1.4 (36.6) | 0.969 | -3.7 (34.5) | 0.916 | -0.9 (0.6) | 0.343 | -0.8 (0.6) | 0.391 | -0.5 (0.3) | 0.245 | -0.5 (0.3) | 0.267 |
| | Right Lobule V | -79.0 (39.4) | 0.104 | -72.4 (37.4) | 0.105 | **-2.3 (0.6)** | **0.003**** | **-2.4 (0.6)** | **0.002**** | -0.7 (0.4) | 0.245 | -0.7 (0.4) | 0.246 |
| | Left Lobule VI | -85.1 (51.3) | 0.141 | -88.1 (48.8) | 0.105 | -0.5 (1.0) | 0.611 | -0.5 (1.0) | 0.593 | -0.2 (0.5) | 0.712 | -0.2 (0.5) | 0.709 |
| | Right Lobule VI | -139.1 (56.3) | 0.072 | -138.0 (53.5) | 0.055 | 0.8 (1.1) | 0.611 | 0.8 (1.1) | 0.593 | 0.3 (0.5) | 0.712 | 0.3 (0.5) | 0.709 |
| | Left Hippocampus | 7.4 (22.0) | 0.791 | 19.5 (21.5) | 0.555 | -0.2 (0.4) | 0.755 | -0.2 (0.4) | 0.772 | 0.1 (0.2) | 0.677 | 0.1 (0.2) | 0.788 |
| | Right Hippocampus | -5.4 (20.3) | 0.791 | 0.0 (19.5) | 1.000 | -0.2 (0.4) | 0.755 | -0.2 (0.4) | 0.772 | 0.2 (0.2) | 0.623 | 0.2 (0.2) | 0.686 |
| | Left Amygdala | -6.4 (18.0) | 0.791 | -12.5 (17.3) | 0.570 | -0.3 (0.3) | 0.755 | -0.4 (0.3) | 0.772 | -0.2 (0.2) | 0.577 | -0.2 (0.2) | 0.642 |
| | Right Amygdala | -27.5 (14.1) | 0.346 | -20.2 (13.8) | 0.303 | -0.1 (0.3) | 0.755 | -0.1 (0.3) | 0.772 | -0.2 (0.1) | 0.577 | -0.2 (0.1) | 0.642 |
| | Left Thalamus | 63.2 (45.0) | 0.366 | 91.9 (45.2) | 0.172 | -0.3 (0.8) | 0.755 | -0.3 (0.9) | 0.772 | -0.3 (0.4) | 0.623 | -0.3 (0.5) | 0.686 |
| | Right Thalamus | 63.3 (46.7) | 0.366 | 90.4 (46.1) | 0.172 | 0.6 (0.9) | 0.755 | 0.6 (0.9) | 0.772 | 0.3 (0.5) | 0.623 | 0.3 (0.5) | 0.686 |
| **Males** | Left Crus I | 100.9 (113.0) | 0.505 | 215.2 (115.6) | 0.095 | 0.3 (1.4) | 0.907 | 0.2 (1.5) | 0.910 | -0.1 (1.2) | 0.949 | 0.1 (1.3) | 0.961 |
| | Right Crus I | 207.0 (105.3) | 0.231 | 261.7 (103.2) | 0.064 | 2.0 (1.3) | 0.549 | 1.9 (1.3) | 0.664 | 0.9 (1.1) | 0.949 | 1.0 (1.2) | 0.961 |
| | Left Crus II | 27.4 (93.5) | 0.772 | 73.7 (92.2) | 0.430 | -0.1 (1.1) | 0.907 | -0.2 (1.1) | 0.910 | -0.1 (1.0) | 0.949 | 0.1 (1.0) | 0.961 |
| | Right Crus II | 223.5 (139.3) | 0.236 | 275.7 (136.3) | 0.095 | 0.6 (1.7) | 0.907 | 0.5 (1.8) | 0.910 | -0.2 (1.5) | 0.949 | -0.1 (1.5) | 0.961 |
| | Left Lobule V | -5.9 (45.1) | 0.897 | 13.7 (45.4) | 0.765 | -0.1 (0.5) | 0.843 | -0.1 (0.6) | 0.797 | 0.0 (0.5) | 0.956 | 0.0 (0.5) | 0.961 |
| | Right Lobule V | 29.4 (43.4) | 0.670 | 46.4 (43.4) | 0.390 | 0.4 (0.5) | 0.652 | 0.3 (0.5) | 0.733 | 0.3 (0.4) | 0.623 | 0.4 (0.5) | 0.572 |
| | Left Lobule VI | 79.2 (52.2) | 0.321 | 98.7 (51.2) | 0.125 | 0.8 (0.6) | 0.459 | 0.8 (0.6) | 0.632 | 0.9 (0.5) | 0.191 | 1.0 (0.5) | 0.165 |
| | Right Lobule VI | 104.3 (72.7) | 0.321 | 153.2 (73.2) | 0.125 | 1.0 (0.8) | 0.459 | 0.9 (0.9) | 0.632 | 1.2 (0.7) | 0.191 | 1.4 (0.8) | 0.165 |
| | Left Hippocampus | 23.5 (25.8) | 0.554 | 31.0 (25.0) | 0.365 | 0.1 (0.3) | 0.830 | 0.1 (0.3) | 0.806 | 0.0 (0.3) | 0.909 | 0.0 (0.3) | 0.854 |
| | Right Hippocampus | 43.8 (28.4) | 0.398 | 45.7 (27.7) | 0.365 | 0.1 (0.4) | 0.830 | 0.1 (0.4) | 0.806 | 0.1 (0.3) | 0.909 | 0.1 (0.3) | 0.854 |
| | Left Amygdala | 7.6 (21.1) | 0.722 | 24.7 (20.8) | 0.365 | 0.2 (0.2) | 0.830 | 0.2 (0.3) | 0.806 | -0.1 (0.2) | 0.909 | 0.0 (0.2) | 0.854 |
| | Right Amygdala | 9.4 (17.4) | 0.712 | 19.8 (16.9) | 0.365 | 0.1 (0.2) | 0.830 | 0.1 (0.2) | 0.806 | 0.1 (0.2) | 0.909 | 0.1 (0.2) | 0.854 |
| | Left Thalamus | 83.5 (47.3) | 0.398 | 52.0 (49.9) | 0.365 | 0.5 (0.5) | 0.830 | 0.6 (0.6) | 0.806 | 0.7 (0.4) | 0.496 | 0.6 (0.5) | 0.653 |
| | Right Thalamus | 67.6 (65.3) | 0.554 | 59.3 (64.5) | 0.365 | -0.3 (0.7) | 0.830 | -0.3 (0.7) | 0.806 | 0.8 (0.6) | 0.496 | 0.8 (0.6) | 0.653 |

*Note.* Change-score models (ΔHormone predicting ΔVolume) repeated with the between-session interval as a covariate, shown for both scan-to-scan and saliva-to-saliva intervals. Outliers removed (11 subjects; 3*SD* within sex and timepoint). E = 17β-estradiol; T = testosterone; P = progesterone; *b* = unstandardized coefficient; *SE* = standard error. *p*-values FDR-corrected within family (Crus I/II; Lobule V/VI; limbic). **p*_FDR < .05, ***p*_FDR < .01.

**Table S14.** Within-session saliva-to-scan collection interval sensitivity check with outliers.

| | | E | | | | T | | | | P | | | |
|---|---|---|---|---|---|---|---|---|---|---|---|---|---|
| | | Baseline | | Year 1 | | Baseline | | Year 1 | | Baseline | | Year 1 | |
| **Sample** | **Region** | *b (SE)* | *p* | *b (SE)* | *p* | *b (SE)* | *p* | *b (SE)* | *p* | *b (SE)* | *p* | *b (SE)* | *p* |
| **Full sample** | Left Crus I | -219.8 (174.5) | 0.210 | -457.7 (201.2) | 0.067 | 3.1 (2.5) | 0.413 | -2.4 (4.2) | 0.564 | -3.4 (1.9) | 0.158 | -4.4 (1.9) | 0.042* |
| | Right Crus I | -306.8 (179.2) | 0.166 | -467.2 (216.3) | 0.067 | 1.1 (2.6) | 0.660 | -6.5 (4.4) | 0.484 | -4.5 (2.0) | 0.094 | -5.1 (2.0) | 0.042* |
| | Left Crus II | -254.8 (130.6) | 0.166 | -81.0 (163.1) | 0.621 | 1.3 (1.9) | 0.657 | 3.1 (3.3) | 0.484 | -1.1 (1.5) | 0.605 | 0.7 (1.6) | 0.676 |
| | Right Crus II | -216.0 (139.7) | 0.166 | -116.9 (164.2) | 0.621 | 3.3 (2.0) | 0.372 | 3.0 (3.3) | 0.484 | 0.4 (1.6) | 0.794 | 1.7 (1.6) | 0.362 |
| | Left Lobule V | -18.8 (47.2) | 0.691 | 7.3 (54.4) | 0.893 | 0.9 (0.7) | 0.328 | -1.6 (1.1) | 0.581 | 0.2 (0.5) | 0.808 | -0.4 (0.5) | 0.730 |
| | Right Lobule V | -21.9 (43.1) | 0.691 | -64.9 (52.2) | 0.434 | 1.0 (0.6) | 0.328 | -0.3 (1.1) | 0.805 | -0.1 (0.5) | 0.808 | 0.0 (0.5) | 0.970 |
| | Left Lobule VI | -218.9 (103.3) | 0.144 | -217.9 (126.6) | 0.356 | -0.7 (1.5) | 0.842 | -1.9 (2.6) | 0.805 | -2.1 (1.2) | 0.164 | -1.4 (1.2) | 0.730 |
| | Right Lobule VI | -171.1 (99.2) | 0.174 | -31.4 (121.4) | 0.893 | -0.3 (1.4) | 0.857 | -0.6 (2.4) | 0.805 | -1.9 (1.1) | 0.164 | -0.7 (1.2) | 0.730 |
| | Left Hippocampus | -26.0 (40.4) | 0.787 | -101.6 (45.1) | 0.080 | -0.4 (0.6) | 0.790 | -0.5 (0.9) | 0.857 | 0.0 (0.5) | 0.999 | 0.1 (0.4) | 0.785 |
| | Right Hippocampus | 11.3 (41.7) | 0.787 | -51.5 (47.8) | 0.427 | 0.1 (0.6) | 0.802 | -1.1 (0.9) | 0.857 | 0.5 (0.5) | 0.440 | 0.2 (0.5) | 0.785 |
| | Left Amygdala | -5.4 (16.7) | 0.787 | **-54.0 (19.1)** | **0.035*** | -0.1 (0.2) | 0.802 | -0.3 (0.4) | 0.857 | -0.4 (0.2) | 0.302 | -0.2 (0.2) | 0.785 |
| | Right Amygdala | -8.6 (18.1) | 0.787 | -40.5 (20.9) | 0.111 | -0.3 (0.3) | 0.606 | 0.0 (0.4) | 0.970 | -0.3 (0.2) | 0.440 | -0.1 (0.2) | 0.785 |
| | Left Thalamus | 68.0 (67.8) | 0.787 | 47.7 (80.4) | 0.650 | 0.6 (1.0) | 0.790 | 0.7 (1.6) | 0.857 | 0.2 (0.8) | 0.955 | 0.4 (0.8) | 0.785 |
| | Right Thalamus | 117.2 (62.3) | 0.373 | 33.2 (73.0) | 0.650 | 1.6 (0.9) | 0.451 | 0.5 (1.5) | 0.857 | 0.7 (0.7) | 0.440 | 0.7 (0.7) | 0.785 |
| **Females** | Left Crus I | -354.4 (184.1) | 0.117 | -598.8 (274.6) | 0.074 | -3.3 (4.5) | 0.730 | -2.3 (6.6) | 0.729 | -2.6 (2.0) | 0.384 | -3.3 (2.0) | 0.221 |
| | Right Crus I | -487.8 (202.0) | 0.074 | -701.6 (325.3) | 0.074 | -7.2 (4.9) | 0.604 | -8.8 (7.7) | 0.729 | -3.6 (2.2) | 0.384 | -4.3 (2.4) | 0.221 |
| | Left Crus II | -225.0 (157.3) | 0.208 | 83.3 (245.7) | 0.889 | -1.9 (3.9) | 0.730 | 4.0 (5.6) | 0.729 | 0.0 (1.8) | 0.998 | 0.2 (1.9) | 0.894 |
| | Right Crus II | -205.1 (161.5) | 0.208 | -33.5 (238.3) | 0.889 | 1.4 (3.9) | 0.730 | 2.7 (5.4) | 0.729 | 1.2 (1.8) | 0.652 | 1.7 (1.8) | 0.437 |
| | Left Lobule V | -63.2 (50.7) | 0.217 | 43.4 (79.2) | 0.587 | -0.3 (1.2) | 0.780 | -1.1 (1.8) | 0.674 | -0.4 (0.6) | 0.533 | -0.5 (0.6) | 0.378 |
| | Right Lobule V | -82.6 (45.8) | 0.217 | -82.5 (68.9) | 0.476 | 0.8 (1.1) | 0.780 | -1.0 (1.6) | 0.674 | -0.5 (0.5) | 0.476 | -0.5 (0.5) | 0.378 |
| | Left Lobule VI | -189.5 (118.4) | 0.217 | -330.1 (194.2) | 0.386 | -1.4 (2.9) | 0.780 | -3.2 (4.6) | 0.674 | -2.7 (1.3) | 0.084 | -3.2 (1.4) | 0.136 |
| | Right Lobule VI | -147.1 (116.6) | 0.217 | -129.0 (188.0) | 0.587 | -2.1 (2.9) | 0.780 | -1.8 (4.3) | 0.674 | -2.6 (1.2) | 0.084 | -2.1 (1.4) | 0.294 |
| | Left Hippocampus | 42.7 (41.5) | 0.307 | -73.8 (59.1) | 0.656 | -0.4 (1.0) | 0.806 | -0.1 (1.4) | 0.929 | 0.4 (0.5) | 0.577 | 0.2 (0.4) | 0.967 |
| | Right Hippocampus | 70.6 (43.9) | 0.270 | -5.9 (61.9) | 0.924 | 1.0 (1.1) | 0.698 | -0.1 (1.4) | 0.929 | 0.9 (0.5) | 0.397 | 0.2 (0.5) | 0.967 |
| | Left Amygdala | 19.0 (18.2) | 0.307 | -38.0 (28.6) | 0.656 | -0.8 (0.4) | 0.311 | -0.2 (0.7) | 0.929 | -0.2 (0.2) | 0.577 | 0.0 (0.2) | 0.967 |
| | Right Amygdala | 27.8 (18.4) | 0.270 | 4.7 (28.1) | 0.924 | 0.0 (0.5) | 0.932 | 0.4 (0.6) | 0.929 | -0.1 (0.2) | 0.802 | 0.1 (0.2) | 0.967 |
| | Left Thalamus | 105.0 (82.0) | 0.307 | 19.7 (126.1) | 0.924 | 0.9 (2.0) | 0.806 | 3.6 (2.8) | 0.642 | 0.6 (0.9) | 0.577 | 0.0 (1.0) | 0.967 |
| | Right Thalamus | 140.0 (70.1) | 0.270 | 24.7 (103.2) | 0.924 | 2.8 (1.7) | 0.311 | 3.4 (2.3) | 0.642 | 1.1 (0.8) | 0.423 | 0.6 (0.8) | 0.967 |
| **Males** | Left Crus I | 67.8 (359.7) | 0.851 | -197.4 (318.3) | 0.695 | 4.9 (3.1) | 0.245 | -1.6 (5.9) | 0.787 | -5.9 (5.0) | 0.243 | -6.2 (4.3) | 0.304 |
| | Right Crus I | 73.3 (331.7) | 0.851 | -121.2 (306.7) | 0.695 | 3.1 (2.9) | 0.298 | -5.8 (5.6) | 0.786 | -6.0 (4.6) | 0.243 | -6.7 (4.0) | 0.304 |
| | Left Crus II | -345.3 (247.8) | 0.675 | -321.1 (232.9) | 0.492 | 2.6 (2.2) | 0.298 | 2.4 (4.4) | 0.786 | -7.4 (3.4) | 0.135 | -0.7 (3.3) | 0.982 |
| | Right Crus II | -261.6 (276.0) | 0.694 | -290.9 (246.9) | 0.492 | 4.5 (2.4) | 0.245 | 3.9 (4.6) | 0.786 | -5.2 (3.8) | 0.243 | -0.1 (3.4) | 0.982 |
| | Left Lobule V | 67.0 (91.2) | 0.466 | -15.6 (75.3) | 0.837 | 1.5 (0.8) | 0.219 | -1.4 (1.4) | 0.398 | 1.0 (1.3) | 0.832 | -0.9 (1.0) | 0.791 |
| | Right Lobule V | 103.6 (80.0) | 0.278 | -36.5 (70.8) | 0.837 | 1.0 (0.7) | 0.315 | -0.4 (1.3) | 0.786 | 1.3 (1.1) | 0.832 | -0.3 (1.0) | 0.791 |
| | Left Lobule VI | -316.4 (204.2) | 0.278 | -155.4 (164.7) | 0.837 | -0.6 (1.9) | 0.816 | -3.9 (3.0) | 0.398 | -0.6 (2.9) | 0.837 | 1.3 (2.3) | 0.791 |
| | Right Lobule VI | -245.0 (192.6) | 0.278 | -31.6 (144.5) | 0.837 | 0.4 (1.7) | 0.816 | -3.3 (2.5) | 0.398 | -1.3 (2.7) | 0.832 | -0.6 (1.9) | 0.791 |
| | Left Hippocampus | **-220.5 (76.2)** | **0.018*** | -145.5 (65.9) | 0.067 | -0.5 (0.7) | 0.862 | -1.8 (1.2) | 0.428 | -1.2 (1.1) | 0.503 | -0.8 (0.9) | 0.802 |
| | Right Hippocampus | -157.2 (79.4) | 0.079 | -102.8 (71.9) | 0.241 | -0.3 (0.7) | 0.862 | -2.6 (1.2) | 0.246 | -0.3 (1.1) | 0.762 | -0.4 (1.0) | 0.919 |
| | Left Amygdala | -69.5 (31.4) | 0.062 | -66.3 (28.0) | 0.067 | 0.0 (0.3) | 0.934 | -0.6 (0.5) | 0.518 | -0.7 (0.4) | 0.415 | -0.6 (0.4) | 0.802 |
| | Right Amygdala | **-100.2 (35.2)** | **0.018*** | -84.5 (31.9) | 0.067 | -0.6 (0.3) | 0.562 | -0.5 (0.6) | 0.518 | -0.8 (0.5) | 0.415 | -0.4 (0.5) | 0.802 |
| | Left Thalamus | -46.2 (123.9) | 0.726 | 49.9 (108.9) | 0.780 | 0.5 (1.1) | 0.862 | -1.4 (2.0) | 0.518 | -1.7 (1.7) | 0.503 | 0.2 (1.5) | 0.919 |
| | Right Thalamus | 43.4 (123.6) | 0.726 | 23.6 (109.0) | 0.830 | 1.1 (1.1) | 0.862 | -1.3 (2.0) | 0.518 | -0.6 (1.7) | 0.762 | -0.2 (1.5) | 0.919 |

*Note.* Cross-sectional models repeated with the within-session scan-to-saliva interval as a covariate (Baseline models use the Baseline interval; Year 1 models use the Year 1 interval). Full sample with outliers retained. E = 17β-estradiol; T = testosterone; P = progesterone; *b* = unstandardized coefficient; *SE* = standard error. *p*-values FDR-corrected within family (Crus I/II; Lobule V/VI; limbic). **p*_FDR < .05.

**Table S15.** Within-session saliva-to-scan collection interval sensitivity check without outliers.

| | | E | | | | T | | | | P | | | |
|---|---|---|---|---|---|---|---|---|---|---|---|---|---|
| | | **Baseline** | | **Year 1** | | **Baseline** | | **Year 1** | | **Baseline** | | **Year 1** | |
| **Sample** | **Region** | *b (SE)* | *p* | *b (SE)* | *p* | *b (SE)* | *p* | *b (SE)* | *p* | *b (SE)* | *p* | *b (SE)* | *p* |
| **Full sample** | Left Crus I | -141.6 (229.0) | 0.537 | -279.9 (302.4) | 0.358 | 2.1 (3.0) | 0.664 | -1.2 (4.0) | 0.766 | -1.7 (2.4) | 0.919 | -3.9 (2.4) | 0.234 |
| | Right Crus I | -236.4 (235.7) | 0.439 | -404.4 (328.0) | 0.358 | 0.7 (3.2) | 0.815 | -5.0 (4.3) | 0.546 | -2.9 (2.5) | 0.919 | -5.4 (2.6) | 0.160 |
| | Left Crus II | -247.4 (186.6) | 0.439 | 368.2 (269.3) | 0.358 | 2.7 (2.5) | 0.576 | 3.5 (3.6) | 0.546 | -0.8 (2.0) | 0.919 | 1.0 (2.2) | 0.639 |
| | Right Crus II | -195.5 (199.4) | 0.439 | 287.6 (268.8) | 0.358 | 4.5 (2.6) | 0.338 | 2.9 (3.5) | 0.546 | -0.1 (2.1) | 0.962 | 2.1 (2.2) | 0.469 |
| | Left Lobule V | -66.0 (65.4) | 0.420 | -71.2 (88.4) | 0.500 | 0.3 (0.9) | 0.839 | -2.4 (1.1) | 0.155 | -0.4 (0.7) | 0.552 | -1.2 (0.7) | 0.327 |
| | Right Lobule V | -37.5 (58.0) | 0.519 | -55.7 (82.1) | 0.500 | 0.7 (0.8) | 0.839 | -0.8 (1.1) | 0.788 | -0.7 (0.6) | 0.499 | -0.8 (0.7) | 0.443 |
| | Left Lobule VI | -224.8 (142.5) | 0.235 | -202.2 (203.9) | 0.500 | 0.4 (1.9) | 0.839 | -1.4 (2.7) | 0.788 | -1.2 (1.5) | 0.549 | -1.1 (1.7) | 0.522 |
| | Right Lobule VI | -272.5 (140.4) | 0.218 | -176.5 (200.0) | 0.500 | 0.7 (1.9) | 0.839 | -0.6 (2.6) | 0.825 | -1.8 (1.5) | 0.499 | -1.2 (1.6) | 0.522 |
| | Left Hippocampus | -67.3 (55.6) | 0.698 | -44.6 (74.6) | 0.662 | 0.2 (0.7) | 0.900 | -0.6 (1.0) | 0.812 | -0.1 (0.6) | 0.888 | 0.0 (0.6) | 0.968 |
| | Right Hippocampus | -32.3 (58.5) | 0.698 | -26.0 (79.2) | 0.744 | 0.4 (0.8) | 0.900 | -1.4 (1.0) | 0.812 | 0.4 (0.6) | 0.639 | -0.1 (0.6) | 0.968 |
| | Left Amygdala | -25.2 (22.5) | 0.698 | -34.9 (31.1) | 0.528 | 0.0 (0.3) | 0.900 | -0.3 (0.4) | 0.812 | -0.5 (0.2) | 0.178 | -0.2 (0.3) | 0.954 |
| | Right Amygdala | -17.7 (24.2) | 0.698 | -19.7 (32.7) | 0.662 | 0.1 (0.3) | 0.900 | 0.1 (0.4) | 0.812 | -0.3 (0.3) | 0.331 | 0.0 (0.3) | 0.968 |
| | Left Thalamus | 7.6 (96.0) | 0.937 | 158.7 (132.0) | 0.528 | 0.9 (1.3) | 0.900 | 0.8 (1.8) | 0.812 | 1.3 (1.0) | 0.331 | 2.1 (1.1) | 0.160 |
| | Right Thalamus | 67.2 (87.6) | 0.698 | 145.1 (118.0) | 0.528 | 1.8 (1.1) | 0.724 | 0.4 (1.6) | 0.812 | 1.7 (0.9) | 0.178 | 2.3 (0.9) | 0.090 |
| **Females** | Left Crus I | -374.0 (273.0) | 0.352 | -391.4 (386.0) | 0.317 | -1.4 (4.9) | 0.876 | -0.8 (6.2) | 0.899 | -1.6 (2.7) | 0.555 | -3.1 (2.8) | 0.379 |
| | Right Crus I | -603.3 (302.2) | 0.202 | -699.3 (463.4) | 0.303 | -5.9 (5.4) | 0.876 | -7.6 (7.5) | 0.741 | -3.0 (3.0) | 0.555 | -5.8 (3.4) | 0.365 |
| | Left Crus II | -244.3 (244.7) | 0.430 | 521.0 (361.7) | 0.303 | -2.0 (4.4) | 0.876 | 4.8 (5.9) | 0.741 | 1.9 (2.4) | 0.555 | 1.2 (2.7) | 0.648 |
| | Right Crus II | -182.2 (251.9) | 0.472 | 427.5 (348.4) | 0.303 | 0.7 (4.4) | 0.876 | 3.3 (5.6) | 0.741 | 1.6 (2.5) | 0.555 | 2.8 (2.6) | 0.379 |
| | Left Lobule V | -106.1 (75.5) | 0.201 | -22.8 (115.6) | 0.845 | -1.3 (1.3) | 0.943 | -0.7 (1.8) | 0.890 | -0.7 (0.8) | 0.392 | -0.5 (0.8) | 0.522 |
| | Right Lobule V | -133.9 (65.5) | 0.181 | -98.0 (100.2) | 0.495 | -0.4 (1.2) | 0.943 | -0.5 (1.6) | 0.890 | -0.9 (0.7) | 0.392 | -0.7 (0.7) | 0.522 |
| | Left Lobule VI | -231.0 (178.8) | 0.201 | -447.4 (272.0) | 0.433 | -0.2 (3.2) | 0.943 | -1.8 (4.5) | 0.890 | -1.6 (1.7) | 0.392 | -3.0 (2.0) | 0.522 |
| | Right Lobule VI | -237.4 (178.4) | 0.201 | -244.9 (270.6) | 0.495 | -1.7 (3.3) | 0.943 | -0.6 (4.4) | 0.890 | -2.1 (1.7) | 0.392 | -1.7 (2.0) | 0.522 |
| | Left Hippocampus | 28.4 (63.6) | 0.788 | -28.9 (89.7) | 0.972 | -0.5 (1.1) | 0.688 | -0.1 (1.4) | 0.922 | 0.6 (0.6) | 0.399 | 0.1 (0.7) | 0.858 |
| | Right Hippocampus | 60.9 (67.3) | 0.788 | -0.8 (93.7) | 0.993 | 0.7 (1.2) | 0.656 | -0.4 (1.5) | 0.922 | 1.0 (0.7) | 0.198 | -0.1 (0.7) | 0.858 |
| | Left Amygdala | -3.9 (26.7) | 0.886 | -29.9 (37.5) | 0.858 | -0.6 (0.5) | 0.634 | -0.4 (0.6) | 0.922 | -0.4 (0.3) | 0.198 | -0.1 (0.3) | 0.858 |
| | Right Amygdala | 15.3 (27.8) | 0.788 | 9.4 (39.0) | 0.972 | 0.4 (0.5) | 0.645 | 0.3 (0.6) | 0.922 | 0.0 (0.3) | 0.894 | 0.1 (0.3) | 0.858 |
| | Left Thalamus | 104.3 (125.6) | 0.788 | 206.8 (175.6) | 0.739 | 2.2 (2.2) | 0.634 | 4.1 (2.8) | 0.426 | 2.4 (1.2) | 0.143 | 2.3 (1.3) | 0.229 |
| | Right Thalamus | 139.2 (108.3) | 0.788 | 235.2 (146.3) | 0.698 | 3.1 (1.9) | 0.591 | 4.1 (2.3) | 0.426 | 2.5 (1.0) | 0.094 | **3.0 (1.0)** | **0.029*** |
| **Males** | Left Crus I | 210.0 (396.5) | 0.599 | -191.8 (532.3) | 0.912 | 3.1 (4.2) | 0.530 | -2.3 (6.0) | 0.707 | -0.2 (5.6) | 0.975 | -6.0 (5.0) | 0.478 |
| | Right Crus I | 331.5 (359.8) | 0.599 | -143.6 (502.1) | 0.912 | 2.4 (3.8) | 0.530 | -6.0 (5.5) | 0.707 | 0.6 (5.1) | 0.975 | -6.0 (4.6) | 0.478 |
| | Left Crus II | -283.7 (302.1) | 0.599 | -70.8 (433.7) | 0.912 | 5.8 (3.1) | 0.132 | 2.5 (4.9) | 0.707 | **-11.3 (4.0)** | **0.028*** | 0.0 (4.1) | 0.994 |
| | Right Crus II | -239.3 (337.1) | 0.599 | -51.1 (457.8) | 0.912 | 7.5 (3.4) | 0.124 | 3.2 (5.2) | 0.707 | -7.6 (4.6) | 0.213 | 0.6 (4.4) | 0.994 |
| | Left Lobule V | -25.3 (108.8) | 0.817 | -192.8 (127.7) | 0.281 | 1.4 (1.1) | 0.516 | -3.0 (1.4) | 0.176 | -0.4 (1.5) | 0.915 | -2.2 (1.2) | 0.279 |
| | Right Lobule V | 105.8 (95.0) | 0.494 | -92.0 (125.3) | 0.540 | 1.1 (1.0) | 0.516 | -1.6 (1.4) | 0.273 | 0.7 (1.4) | 0.915 | -1.5 (1.2) | 0.355 |
| | Left Lobule VI | -218.7 (242.1) | 0.494 | -180.5 (291.7) | 0.540 | 0.6 (2.6) | 0.825 | -4.7 (3.2) | 0.201 | 0.4 (3.4) | 0.915 | 0.8 (2.8) | 0.763 |
| | Right Lobule VI | -350.3 (234.4) | 0.494 | -458.8 (255.6) | 0.281 | 2.2 (2.5) | 0.516 | -4.4 (2.8) | 0.201 | -2.2 (3.3) | 0.915 | -2.7 (2.4) | 0.355 |
| | Left Hippocampus | -212.8 (91.0) | 0.139 | -152.2 (122.1) | 0.585 | 0.3 (1.0) | 0.964 | -2.0 (1.3) | 0.453 | -1.8 (1.3) | 0.510 | -1.3 (1.1) | 0.707 |
| | Right Hippocampus | -170.4 (96.3) | 0.248 | -111.7 (135.2) | 0.622 | 0.0 (1.0) | 0.964 | -3.2 (1.4) | 0.180 | -0.9 (1.4) | 0.639 | -0.8 (1.2) | 0.759 |
| | Left Amygdala | -51.5 (38.1) | 0.273 | -56.5 (52.8) | 0.585 | 0.2 (0.4) | 0.964 | -0.5 (0.6) | 0.538 | -0.6 (0.5) | 0.510 | -0.7 (0.5) | 0.707 |
| | Right Amygdala | -61.7 (41.9) | 0.273 | -62.3 (57.1) | 0.585 | -0.2 (0.5) | 0.964 | -0.2 (0.7) | 0.749 | -0.8 (0.6) | 0.510 | -0.5 (0.5) | 0.707 |
| | Left Thalamus | -153.1 (149.0) | 0.371 | -90.9 (195.4) | 0.645 | 0.3 (1.6) | 0.964 | -2.3 (2.2) | 0.464 | -1.9 (2.1) | 0.546 | 0.5 (1.9) | 0.837 |
| | Right Thalamus | -47.8 (147.4) | 0.747 | -104.5 (191.2) | 0.645 | 1.1 (1.5) | 0.964 | -2.3 (2.1) | 0.464 | -0.1 (2.1) | 0.959 | 0.4 (1.8) | 0.837 |

*Note.* Cross-sectional models repeated with the within-session scan-to-saliva interval as a covariate (Baseline models use the Baseline interval; Year 1 models use the Year 1 interval). Outliers removed (11 subjects; 3*SD* within sex and timepoint). E = 17β-estradiol; T = testosterone; P = progesterone; *b* = unstandardized coefficient; *SE* = standard error. *p*-values FDR-corrected within family (Crus I/II; Lobule V/VI; limbic). *$p$_FDR < .05.

**Table S16.** Reproductive stage-stratified hormone-volume associations: Baseline, Year 1, and Baseline-predicting Year 1 models for all regions, with outliers.

| | | Baseline | | | | | | Year 1 | | | | | | Baseline→Year 1 | | | | | |
|---|---|---|---|---|---|---|---|---|---|---|---|---|---|---|---|---|---|---|---|
| | | E | | T | | P | | E | | T | | P | | E | | T | | P | |
| **Stage** | **Region** | *b (SE)* | *p* | *b (SE)* | *p* | *b (SE)* | *p* | *b (SE)* | *p* | *b (SE)* | *p* | *b (SE)* | *p* | *b (SE)* | *p* | *b (SE)* | *p* | *b (SE)* | *p* |
| | | *n* = 17 | | *n* = 17 | | *n* = 16 | | *n* = 10 | | *n* = 10 | | *n* = 10 | | *n* = 11 | | *n* = 10 | | *n* = 10 | |
| **Reproductive** | Left Crus I | -769.6 (634.5) | 0.386 | -9.1 (9.8) | 0.520 | -1.1 (2.0) | 0.678 | 512.4 (964.5) | 0.765 | -7.6 (17.2) | 0.679 | -1.9 (3.0) | 0.786 | 45.5 (151.9) | 0.777 | -6.0 (2.4) | 0.257 | -0.4 (0.5) | 0.970 |
| | Right Crus I | -792.8 (715.7) | 0.386 | -12.7 (10.4) | 0.520 | -0.9 (2.2) | 0.678 | 638.1 (1231.8) | 0.765 | -13.2 (21.6) | 0.679 | -2.2 (3.9) | 0.786 | 314.4 (215.6) | 0.409 | -6.1 (4.3) | 0.456 | 0.0 (0.7) | 0.970 |
| | Left Crus II | -643.5 (506.4) | 0.386 | -9.2 (10.3) | 0.520 | 1.0 (2.1) | 0.678 | 601.0 (1060.2) | 0.765 | -13.3 (18.4) | 0.679 | -1.1 (3.4) | 0.786 | 683.9 (363.3) | 0.409 | 0.5 (10.6) | 0.964 | 0.8 (1.4) | 0.970 |
| | Right Crus II | -327.2 (510.4) | 0.534 | 4.3 (9.8) | 0.667 | 1.9 (1.9) | 0.678 | 282.3 (892.8) | 0.765 | -16.2 (14.2) | 0.679 | -0.8 (2.8) | 0.786 | 313.0 (485.8) | 0.730 | -10.5 (11.6) | 0.556 | -0.7 (2.0) | 0.970 |
| | Left Lobule V | -24.9 (243.8) | 0.937 | 1.0 (4.7) | 0.831 | 0.4 (0.9) | 0.831 | 20.6 (376.0) | 0.958 | -7.2 (5.8) | 0.404 | -0.3 (1.2) | 0.819 | -18.7 (162.0) | 0.913 | -5.0 (2.7) | 0.441 | -0.6 (0.4) | 0.888 |
| | Right Lobule V | -14.0 (174.4) | 0.937 | 2.3 (3.3) | 0.673 | 0.1 (0.7) | 0.831 | -105.3 (262.3) | 0.958 | -7.2 (3.4) | 0.366 | -0.5 (0.8) | 0.716 | 89.6 (118.8) | 0.913 | -3.0 (2.6) | 0.441 | -0.2 (0.4) | 0.888 |
| | Left Lobule VI | -929.4 (372.8) | 0.089 | -12.5 (8.3) | 0.591 | -1.6 (1.7) | 0.831 | -455.3 (828.1) | 0.958 | -11.5 (14.2) | 0.454 | -4.4 (1.8) | 0.253 | -20.8 (106.6) | 0.913 | -1.9 (1.7) | 0.441 | 0.0 (0.2) | 0.888 |
| | Right Lobule VI | -792.6 (352.9) | 0.089 | -8.9 (8.2) | 0.591 | -0.8 (1.6) | 0.831 | -289.2 (770.2) | 0.958 | -14.1 (12.3) | 0.404 | -2.8 (2.1) | 0.475 | 219.1 (232.4) | 0.913 | -4.2 (5.1) | 0.449 | -0.3 (0.6) | 0.888 |
| | Left Hippocampus | -30.3 (148.6) | 0.844 | 0.5 (2.8) | 0.871 | 0.6 (0.5) | 0.524 | 282.3 (126.7) | 0.246 | 2.4 (3.0) | 0.688 | 0.6 (0.5) | 0.492 | -22.8 (49.3) | 0.664 | -1.3 (1.0) | 0.707 | -0.1 (0.2) | 0.873 |
| | Right Hippocampus | 31.4 (156.5) | 0.844 | 4.0 (2.6) | 0.454 | 0.8 (0.5) | 0.524 | 291.8 (178.9) | 0.246 | 3.1 (3.7) | 0.688 | 0.9 (0.6) | 0.492 | 31.2 (48.3) | 0.656 | -1.1 (1.0) | 0.707 | -0.1 (0.2) | 0.873 |
| | Left Amygdala | 79.3 (43.4) | 0.558 | 1.9 (0.9) | 0.325 | 0.1 (0.2) | 0.674 | 86.9 (44.7) | 0.246 | 1.8 (0.7) | 0.130 | 0.2 (0.2) | 0.492 | -36.4 (26.0) | 0.656 | -0.7 (1.2) | 0.720 | 0.0 (0.1) | 0.873 |
| | Right Amygdala | -28.3 (42.4) | 0.844 | 0.2 (0.9) | 0.871 | 0.0 (0.2) | 0.859 | 89.0 (50.0) | 0.246 | 2.0 (0.7) | 0.130 | 0.2 (0.2) | 0.492 | 26.5 (36.2) | 0.656 | 0.1 (0.9) | 0.912 | 0.0 (0.1) | 0.873 |
| | Left Thalamus | -465.0 (397.6) | 0.795 | -2.8 (7.8) | 0.871 | 0.9 (1.6) | 0.674 | 919.7 (760.5) | 0.281 | 8.8 (14.8) | 0.694 | -1.1 (2.7) | 0.750 | -85.9 (126.3) | 0.656 | -1.7 (2.2) | 0.720 | -0.3 (0.3) | 0.873 |
| | Right Thalamus | -101.3 (306.3) | 0.844 | 3.5 (5.7) | 0.871 | 1.3 (1.0) | 0.524 | 671.1 (535.1) | 0.281 | 4.1 (10.7) | 0.717 | -0.6 (1.9) | 0.750 | -121.7 (109.5) | 0.656 | -2.9 (2.6) | 0.707 | -0.7 (0.3) | 0.649 |
| | Left Lobule IX | -243.7 (277.8) | 0.614 | 2.1 (5.6) | 0.896 | 1.0 (1.1) | 0.918 | 507.9 (415.2) | 0.847 | 3.5 (8.2) | 0.810 | -0.3 (1.5) | 0.998 | 58.9 (64.5) | 0.864 | -1.4 (1.3) | 0.777 | -0.1 (0.2) | 0.998 |
| | Right Lobule IX | -210.0 (311.8) | 0.642 | 5.1 (6.1) | 0.836 | 1.3 (1.2) | 0.918 | 655.2 (415.6) | 0.847 | 3.5 (8.9) | 0.810 | 0.0 (1.6) | 0.998 | 77.3 (106.2) | 0.864 | -1.7 (2.1) | 0.777 | 0.1 (0.3) | 0.998 |
| | Vermis VI | -198.2 (72.3) | 0.180 | -2.7 (1.8) | 0.490 | -0.6 (0.3) | 0.918 | -41.1 (89.2) | 0.847 | -2.1 (1.3) | 0.810 | -0.5 (0.2) | 0.383 | 22.4 (35.3) | 0.864 | -1.3 (0.6) | 0.570 | -0.1 (0.1) | 0.829 |
| | Vermis VII | -49.2 (60.3) | 0.614 | -0.2 (1.2) | 0.912 | 0.0 (0.2) | 0.918 | -39.5 (54.4) | 0.847 | 0.4 (1.0) | 0.810 | -0.1 (0.2) | 0.998 | **47.1 (8.8)** | **0.031*** | 0.8 (0.4) | 0.570 | 0.1 (0.1) | 0.829 |
| | Vermis VIII | -44.2 (102.9) | 0.719 | 1.0 (2.1) | 0.896 | -0.1 (0.4) | 0.918 | 125.8 (151.8) | 0.847 | -0.7 (2.8) | 0.810 | -0.2 (0.5) | 0.998 | 84.2 (36.4) | 0.343 | -1.0 (1.0) | 0.777 | 0.0 (0.1) | 0.998 |
| | Vermis IX | -70.2 (45.5) | 0.496 | 0.5 (1.0) | 0.896 | -0.1 (0.2) | 0.918 | 49.5 (69.2) | 0.847 | 0.4 (1.3) | 0.810 | -0.2 (0.2) | 0.998 | -17.1 (31.0) | 0.864 | -0.6 (0.4) | 0.693 | 0.0 (0.1) | 0.998 |
| | Vermis X | -28.6 (31.1) | 0.614 | 0.1 (0.6) | 0.912 | 0.0 (0.1) | 0.918 | -18.8 (42.6) | 0.847 | -0.6 (0.7) | 0.810 | -0.2 (0.1) | 0.383 | -0.6 (17.3) | 0.992 | -0.2 (0.3) | 0.901 | 0.0 (0.0) | 0.998 |
| | Total Gray Matter | -16016.5 (8958.2) | 0.495 | -380.3 (184.5) | 0.437 | -19.8 (37.3) | 0.918 | 12196.0 (18017.3) | 0.847 | 302.0 (304.6) | 0.810 | -7.3 (59.5) | 0.998 | -79.7 (7594.1) | 0.992 | 23.1 (190.0) | 0.965 | -0.1 (25.6) | 0.998 |
| | Cortical Volume | -9675.4 (10220.8) | 0.614 | -370.1 (198.9) | 0.437 | -27.5 (37.7) | 0.918 | 5683.8 (17995.2) | 0.850 | 372.6 (275.2) | 0.810 | 12.1 (57.3) | 0.998 | -2994.3 (7668.2) | 0.890 | 41.7 (197.3) | 0.965 | 2.5 (28.2) | 0.998 |
| | Cerebral White Matter | -5718.8 (15542.5) | 0.719 | -375.9 (274.4) | 0.490 | -10.9 (58.1) | 0.918 | -3760.3 (21085.8) | 0.865 | -111.8 (371.1) | 0.810 | -55.7 (62.2) | 0.998 | -7957.7 (6249.4) | 0.863 | -8.2 (176.3) | 0.965 | -3.0 (25.1) | 0.998 |
| | | *n* = 14 | | *n* = 14 | | *n* = 14 | | *n* = 10 | | *n* = 10 | | *n* = 10 | | *n* = 10 | | *n* = 10 | | *n* = 10 | |
| **Early Postmenopausal** | Left Crus I | -516.3 (745.5) | 0.508 | -5.6 (14.8) | 0.949 | -8.1 (9.0) | 0.390 | -86.6 (1887.2) | 0.965 | 4.5 (22.8) | 0.851 | -6.6 (8.9) | 0.547 | -122.0 (307.6) | 0.712 | 1.3 (6.6) | 0.855 | -3.8 (3.9) | 0.648 |

| | | Baseline | | | | | | Year 1 | | | | | | Baseline→Year 1 | | | | | |
|---|---|---|---|---|---|---|---|---|---|---|---|---|---|---|---|---|---|---|---|
| | | E | | T | | P | | E | | T | | P | | E | | T | | P | |
| **Stage** | **Region** | *b* (SE) | *p* | *b* (SE) | *p* | *b* (SE) | *p* | *b* (SE) | *p* | *b* (SE) | *p* | *b* (SE) | *p* | *b* (SE) | *p* | *b* (SE) | *p* | *b* (SE) | *p* |
| | Right Crus I | -571.0 (827.8) | 0.508 | 0.1 (16.6) | 0.993 | -13.5 (9.4) | 0.390 | -1723.6 (1931.1) | 0.826 | 13.2 (24.5) | 0.851 | -14.2 (8.2) | 0.424 | 440.9 (266.9) | 0.348 | 10.9 (4.6) | 0.159 | 6.8 (4.9) | 0.648 |
| | Left Crus II | -931.2 (543.3) | 0.421 | -21.4 (9.9) | 0.234 | -8.5 (7.1) | 0.390 | 1075.4 (655.4) | 0.647 | -2.6 (9.8) | 0.851 | 4.9 (3.4) | 0.424 | -693.6 (367.9) | 0.348 | -19.1 (8.1) | 0.159 | 2.3 (6.6) | 0.744 |
| | Right Crus II | -935.9 (694.4) | 0.421 | -20.8 (13.1) | 0.294 | -8.8 (8.8) | 0.390 | 556.5 (1163.1) | 0.870 | -26.6 (8.2) | 0.090 | 3.7 (5.7) | 0.547 | -326.8 (305.8) | 0.461 | -6.5 (6.8) | 0.528 | 3.2 (4.2) | 0.648 |
| | Left Lobule V | -425.1 (211.9) | 0.152 | -8.0 (4.2) | 0.257 | **-8.3 (1.4)** | **0.001*** ** | -511.7 (557.7) | 0.401 | -10.4 (5.6) | 0.257 | -4.0 (2.4) | 0.370 | 35.2 (126.5) | 0.965 | 1.4 (2.4) | 0.650 | 2.2 (3.6) | 0.778 |
| | Right Lobule V | -344.9 (158.2) | 0.152 | -5.6 (3.3) | 0.257 | **-6.2 (1.2)** | **0.001** ** | -455.4 (424.9) | 0.401 | -8.1 (4.4) | 0.257 | -3.0 (1.9) | 0.370 | 9.0 (193.7) | 0.965 | 1.7 (3.5) | 0.650 | 0.9 (3.9) | 0.825 |
| | Left Lobule VI | -611.9 (579.7) | 0.319 | -5.8 (11.8) | 0.680 | -5.9 (7.3) | 0.437 | -1614.2 (1301.4) | 0.401 | -6.7 (17.8) | 0.899 | -3.1 (7.3) | 0.686 | 88.5 (114.5) | 0.965 | 2.7 (2.0) | 0.494 | 1.0 (1.3) | 0.778 |
| | Right Lobule VI | -759.8 (656.2) | 0.319 | -5.8 (13.6) | 0.680 | -8.3 (8.1) | 0.437 | -2261.5 (1312.0) | 0.401 | -2.7 (20.0) | 0.899 | -5.4 (7.9) | 0.686 | 288.5 (113.2) | 0.254 | 4.8 (2.6) | 0.494 | 2.5 (1.7) | 0.778 |
| | Left Hippocampus | -3.7 (179.8) | 0.984 | -1.5 (3.5) | 0.675 | -2.8 (2.0) | 0.594 | 65.9 (402.0) | 0.876 | 3.8 (4.6) | 0.527 | -0.5 (2.0) | 0.964 | -4.0 (47.1) | 0.937 | -0.2 (1.0) | 0.839 | 0.9 (0.5) | 0.641 |
| | Right Hippocampus | -46.7 (172.7) | 0.951 | -2.5 (3.3) | 0.675 | -2.9 (1.9) | 0.594 | -123.9 (311.7) | 0.876 | -3.1 (3.6) | 0.527 | -1.1 (1.5) | 0.964 | -17.4 (33.9) | 0.936 | -0.9 (0.6) | 0.441 | 0.5 (0.6) | 0.641 |
| | Left Amygdala | 63.0 (45.1) | 0.588 | 0.5 (1.0) | 0.675 | -0.3 (0.6) | 0.801 | -114.9 (146.2) | 0.876 | 2.7 (1.4) | 0.375 | -0.7 (0.7) | 0.964 | 74.2 (29.9) | 0.409 | **1.8 (0.3)** | **0.026*** | 0.3 (0.6) | 0.641 |
| | Right Amygdala | 94.6 (66.6) | 0.588 | 1.0 (1.4) | 0.675 | 0.3 (0.9) | 0.801 | -38.2 (121.7) | 0.876 | 2.4 (1.0) | 0.363 | -0.1 (0.6) | 0.964 | 43.5 (34.0) | 0.542 | 0.5 (0.8) | 0.839 | -0.4 (0.5) | 0.641 |
| | Left Thalamus | 353.6 (324.8) | 0.609 | 4.9 (6.5) | 0.675 | 1.1 (4.2) | 0.801 | -205.6 (859.7) | 0.876 | 13.1 (8.7) | 0.382 | 0.2 (4.3) | 0.964 | 19.8 (66.1) | 0.936 | -0.3 (1.3) | 0.839 | 0.4 (0.8) | 0.641 |
| | Right Thalamus | 135.8 (207.9) | 0.795 | 2.5 (4.1) | 0.675 | 1.9 (2.5) | 0.801 | -84.1 (458.8) | 0.876 | 2.4 (5.5) | 0.675 | 1.2 (2.2) | 0.964 | -104.5 (74.6) | 0.542 | -2.6 (1.3) | 0.357 | -0.8 (1.1) | 0.641 |
| | Left Lobule IX | -317.9 (193.8) | 0.461 | -6.8 (3.7) | 0.616 | -2.7 (2.5) | 0.797 | 635.7 (577.9) | 0.536 | -7.0 (7.2) | 0.881 | 0.9 (3.2) | 0.784 | 129.2 (76.0) | 0.445 | 2.9 (1.4) | 0.384 | 1.7 (0.9) | 0.508 |
| | Right Lobule IX | -334.7 (251.9) | 0.461 | -7.3 (4.8) | 0.616 | -2.6 (3.3) | 0.797 | 908.8 (618.9) | 0.536 | -4.6 (8.7) | 0.969 | 1.6 (3.6) | 0.784 | 95.0 (55.6) | 0.445 | 2.5 (0.9) | 0.231 | 1.2 (0.7) | 0.508 |
| | Vermis VI | -3.8 (109.1) | 0.973 | 0.3 (2.1) | 0.956 | 1.2 (1.3) | 0.797 | 93.0 (253.7) | 0.729 | 2.6 (2.9) | 0.881 | 0.7 (1.2) | 0.784 | -29.3 (18.1) | 0.445 | -0.6 (0.4) | 0.384 | 0.0 (0.3) | 0.995 |
| | Vermis VII | -13.8 (51.1) | 0.881 | -0.1 (1.0) | 0.956 | 0.0 (0.6) | 0.994 | 202.2 (124.2) | 0.536 | 0.4 (1.9) | 0.985 | 0.5 (0.7) | 0.784 | -37.6 (29.2) | 0.445 | -0.8 (0.6) | 0.446 | 0.0 (0.4) | 0.995 |
| | Vermis VIII | -192.1 (103.8) | 0.461 | -3.1 (2.1) | 0.616 | -2.7 (1.2) | 0.496 | -153.3 (293.4) | 0.693 | 0.1 (3.7) | 0.985 | -1.4 (1.4) | 0.784 | 45.7 (14.0) | 0.308 | **1.1 (0.2)** | **0.027*** | -0.1 (0.4) | 0.995 |
| | Vermis IX | -55.1 (47.5) | 0.461 | -0.6 (1.0) | 0.803 | -1.0 (0.5) | 0.496 | -77.6 (105.8) | 0.693 | -1.7 (1.1) | 0.881 | -0.7 (0.5) | 0.784 | -39.0 (30.1) | 0.445 | -0.4 (0.7) | 0.774 | -0.8 (0.4) | 0.508 |
| | Vermis X | -28.0 (23.0) | 0.461 | -0.4 (0.5) | 0.645 | -0.2 (0.3) | 0.797 | 141.3 (60.8) | 0.536 | -0.1 (1.1) | 0.985 | 0.5 (0.4) | 0.784 | 8.6 (9.9) | 0.574 | 0.0 (0.2) | 0.930 | 0.1 (0.1) | 0.771 |
| | Total Gray Matter | 4543.3 (11627.3) | 0.881 | 46.8 (228.2) | 0.956 | -86.5 (140.5) | 0.797 | -26331.8 (23244.1) | 0.536 | -247.7 (295.8) | 0.881 | -102.0 (121.8) | 0.784 | 3942.3 (4820.6) | 0.574 | 128.0 (89.2) | 0.446 | 7.6 (67.4) | 0.995 |
| | Cortical Volume | 13398.0 (11238.5) | 0.461 | 205.2 (225.8) | 0.645 | 58.3 (146.7) | 0.875 | -16993.9 (29878.5) | 0.693 | -161.2 (366.7) | 0.969 | -44.5 (152.5) | 0.784 | 2877.8 (4454.2) | 0.615 | 91.3 (96.2) | 0.566 | -31.1 (59.8) | 0.995 |
| | Cerebral White Matter | 12719.4 (12782.1) | 0.494 | 270.7 (246.7) | 0.645 | -15.3 (164.7) | 0.994 | -40324.7 (32929.7) | 0.536 | 613.9 (362.9) | 0.881 | -152.3 (174.5) | 0.784 | 2399.3 (5002.9) | 0.657 | 10.0 (106.9) | 0.930 | 11.4 (63.8) | 0.995 |
| | | *n* = 37 | | *n* = 37 | | *n* = 36 | | *n* = 27 | | *n* = 27 | | *n* = 26 | | *n* = 27 | | *n* = 27 | | *n* = 26 | |
| **Late Postmenopausal** | Left Crus I | -326.5 (194.6) | 0.206 | -5.4 (6.1) | 0.472 | -0.1 (6.3) | 0.983 | **-811.4 (269.9)** | **0.026*** | -4.8 (8.2) | 0.564 | -6.2 (3.2) | 0.137 | 27.4 (102.9) | 0.793 | 1.4 (1.9) | 0.448 | 3.8 (2.1) | 0.319 |
| | Right Crus I | -412.5 (238.7) | 0.206 | -8.6 (7.5) | 0.472 | -1.4 (7.8) | 0.983 | -847.5 (380.5) | 0.073 | -12.0 (10.6) | 0.358 | -8.8 (4.6) | 0.137 | 55.1 (114.9) | 0.793 | 2.1 (2.1) | 0.448 | 3.0 (2.3) | 0.416 |
| | Left Crus II | -67.9 (219.7) | 0.759 | 4.9 (6.7) | 0.472 | 1.0 (6.9) | 0.983 | 57.8 (342.8) | 0.868 | 11.0 (8.5) | 0.358 | -0.1 (4.3) | 0.978 | 99.4 (89.6) | 0.560 | 1.5 (1.6) | 0.448 | 1.1 (1.9) | 0.734 |

| | | Baseline | | | | | | Year 1 | | | | | | Baseline→Year 1 | | | | | |
|---|---|---|---|---|---|---|---|---|---|---|---|---|---|---|---|---|---|---|---|
| | | E | | T | | P | | E | | T | | P | | E | | T | | P | |
| **Stage** | **Region** | *b* (SE) | *p* | *b* (SE) | *p* | *b* (SE) | *p* | *b* (SE) | *p* | *b* (SE) | *p* | *b* (SE) | *p* | *b* (SE) | *p* | *b* (SE) | *p* | *b* (SE) | *p* |
| | Right Crus II | -122.9 (212.1) | 0.755 | 7.6 (6.4) | 0.472 | 3.1 (6.6) | 0.983 | -161.2 (315.2) | 0.819 | 9.8 (7.9) | 0.358 | -0.2 (3.8) | 0.978 | 121.0 (84.6) | 0.560 | 1.3 (1.6) | 0.448 | 0.3 (1.8) | 0.852 |
| | Left Lobule V | -49.9 (49.6) | 0.930 | 0.0 (1.5) | 0.999 | -1.6 (1.6) | 0.606 | 43.1 (78.3) | 0.777 | 0.8 (2.0) | 0.711 | -1.0 (0.9) | 0.938 | -0.7 (25.1) | 0.979 | 0.5 (0.4) | 0.526 | 0.7 (0.5) | 0.509 |
| | Right Lobule V | -39.3 (53.1) | 0.930 | 1.5 (1.6) | 0.954 | -0.9 (1.7) | 0.689 | -66.7 (85.7) | 0.777 | 2.6 (2.2) | 0.591 | -0.2 (1.1) | 0.938 | 33.2 (26.3) | 0.585 | 0.9 (0.5) | 0.235 | 0.4 (0.6) | 0.509 |
| | Left Lobule VI | 4.4 (129.2) | 0.973 | 2.8 (3.9) | 0.954 | 1.5 (3.8) | 0.689 | -152.7 (216.2) | 0.777 | 4.5 (5.6) | 0.591 | 0.2 (2.7) | 0.938 | 20.8 (60.9) | 0.979 | 0.3 (1.1) | 0.801 | -1.3 (1.2) | 0.509 |
| | Right Lobule VI | -7.4 (124.8) | 0.973 | -0.7 (3.8) | 0.999 | -4.4 (3.7) | 0.606 | 56.9 (198.6) | 0.777 | 3.9 (5.1) | 0.591 | 1.0 (2.5) | 0.938 | 63.6 (58.9) | 0.585 | 0.8 (1.1) | 0.612 | -0.7 (1.0) | 0.509 |
| | Left Hippocampus | 65.5 (53.2) | 0.273 | -0.5 (1.7) | 0.809 | -1.4 (1.7) | 0.894 | -148.0 (74.0) | 0.348 | -0.8 (2.1) | 0.946 | 0.0 (1.0) | 0.960 | -25.0 (20.9) | 0.733 | -0.1 (0.4) | 0.917 | -0.4 (0.4) | 0.874 |
| | Right Hippocampus | 82.5 (50.1) | 0.220 | 0.9 (1.6) | 0.809 | 0.8 (1.6) | 0.929 | -41.1 (78.5) | 0.967 | 0.6 (2.0) | 0.946 | 0.1 (1.0) | 0.960 | -4.7 (20.5) | 0.927 | 0.4 (0.4) | 0.862 | -0.1 (0.4) | 0.874 |
| | Left Amygdala | 22.7 (25.7) | 0.384 | -0.9 (0.8) | 0.633 | -1.3 (0.8) | 0.664 | -47.3 (33.7) | 0.524 | 0.0 (0.9) | 0.975 | 0.3 (0.4) | 0.666 | 21.7 (18.0) | 0.733 | 0.5 (0.3) | 0.588 | 0.4 (0.4) | 0.874 |
| | Right Amygdala | 27.6 (22.1) | 0.273 | -0.2 (0.7) | 0.809 | 0.1 (0.7) | 0.930 | -1.4 (34.2) | 0.967 | 0.4 (0.9) | 0.946 | 0.5 (0.4) | 0.451 | -5.2 (17.8) | 0.927 | 0.0 (0.3) | 0.917 | 0.2 (0.3) | 0.874 |
| | Left Thalamus | 155.5 (72.9) | 0.122 | 2.4 (2.3) | 0.633 | 0.7 (2.4) | 0.930 | -12.7 (98.8) | 0.967 | 4.0 (2.4) | 0.332 | 2.6 (1.1) | 0.079 | 6.2 (66.5) | 0.927 | 0.2 (1.1) | 0.917 | 0.8 (1.5) | 0.874 |
| | Right Thalamus | 172.0 (71.2) | 0.122 | 3.8 (2.3) | 0.630 | 1.8 (2.3) | 0.894 | -30.0 (104.1) | 0.967 | 4.5 (2.5) | 0.332 | 2.7 (1.1) | 0.079 | -20.4 (71.6) | 0.927 | -0.1 (1.3) | 0.917 | -0.3 (1.6) | 0.874 |
| | Left Lobule IX | -43.3 (76.6) | 0.719 | -1.2 (2.4) | 0.774 | 1.9 (2.4) | 0.856 | 14.9 (107.6) | 0.891 | 2.8 (2.7) | 0.758 | -0.2 (1.4) | 0.971 | -20.7 (24.1) | 0.653 | -0.5 (0.4) | 0.722 | 0.5 (0.5) | 0.652 |
| | Right Lobule IX | -47.9 (70.7) | 0.719 | -0.7 (2.2) | 0.774 | 0.8 (2.2) | 0.856 | 37.3 (105.4) | 0.808 | 1.8 (2.7) | 0.758 | -0.7 (1.3) | 0.843 | -80.6 (27.3) | 0.076 | -0.7 (0.6) | 0.722 | -0.7 (0.7) | 0.652 |
| | Vermis VI | -40.2 (36.1) | 0.685 | 1.0 (1.1) | 0.691 | -0.3 (1.1) | 0.856 | -70.0 (61.2) | 0.529 | -0.5 (1.6) | 0.821 | -1.1 (0.7) | 0.354 | 7.5 (19.0) | 0.696 | 0.2 (0.3) | 0.722 | 0.2 (0.4) | 0.747 |
| | Vermis VII | -26.0 (17.8) | 0.510 | -0.6 (0.6) | 0.691 | -0.4 (0.6) | 0.856 | -54.1 (26.6) | 0.461 | -0.4 (0.7) | 0.758 | -0.3 (0.3) | 0.689 | 4.1 (10.1) | 0.696 | -0.2 (0.2) | 0.722 | -0.3 (0.2) | 0.506 |
| | Vermis VIII | -29.0 (36.2) | 0.719 | -1.1 (1.1) | 0.691 | -0.2 (1.1) | 0.864 | -90.3 (59.1) | 0.461 | -1.3 (1.6) | 0.758 | -1.6 (0.7) | 0.119 | -8.2 (17.8) | 0.696 | 0.2 (0.3) | 0.722 | 0.0 (0.4) | 0.919 |
| | Vermis IX | -26.4 (15.0) | 0.442 | -0.7 (0.5) | 0.691 | -0.7 (0.5) | 0.808 | -32.0 (21.6) | 0.461 | -0.8 (0.6) | 0.758 | -0.6 (0.3) | 0.119 | -7.8 (10.1) | 0.653 | 0.1 (0.2) | 0.722 | 0.2 (0.2) | 0.729 |
| | Vermis X | -6.2 (8.3) | 0.719 | -0.2 (0.3) | 0.691 | -0.1 (0.3) | 0.856 | -12.8 (14.6) | 0.558 | -0.3 (0.4) | 0.758 | -0.4 (0.2) | 0.125 | -7.4 (5.5) | 0.653 | -0.1 (0.1) | 0.722 | 0.0 (0.1) | 0.919 |
| | Total Gray Matter | -1356.9 (3333.7) | 0.763 | -72.1 (101.6) | 0.691 | -104.3 (103.9) | 0.808 | -5776.5 (4216.8) | 0.461 | 68.3 (112.3) | 0.758 | 0.0 (52.2) | 1.000 | 3054.6 (3107.1) | 0.653 | 30.1 (57.0) | 0.722 | 96.6 (61.6) | 0.506 |
| | Cortical Volume | 170.1 (2764.5) | 0.951 | -106.3 (82.6) | 0.691 | -96.3 (85.3) | 0.808 | -3632.0 (3912.1) | 0.558 | 11.3 (102.7) | 0.914 | 19.6 (49.0) | 0.866 | 2838.5 (2619.7) | 0.653 | 17.3 (48.2) | 0.722 | 79.1 (51.4) | 0.506 |
| | Cerebral White Matter | **11212.0 (3584.6)** | **0.037*** | -36.2 (125.4) | 0.774 | -148.2 (126.7) | 0.808 | -3103.3 (5621.4) | 0.733 | 77.3 (144.9) | 0.758 | 58.7 (70.3) | 0.689 | -1842.7 (2432.8) | 0.653 | 32.6 (43.4) | 0.722 | -26.8 (50.0) | 0.747 |
| | | *n* = 64 | | *n* = 63 | | *n* = 62 | | *n* = 44 | | *n* = 43 | | *n* = 43 | | *n* = 44 | | *n* = 44 | | *n* = 44 | |
| | Left Crus I | 56.2 (354.9) | 0.928 | 4.8 (3.1) | 0.257 | -5.4 (4.9) | 0.274 | -277.7 (312.8) | 0.603 | -3.1 (5.8) | 0.596 | -7.3 (4.1) | 0.173 | 21.9 (92.4) | 0.814 | -0.5 (0.8) | 0.671 | -1.4 (1.3) | 0.725 |
| | Right Crus I | 29.9 (331.3) | 0.928 | 2.8 (2.9) | 0.343 | -5.1 (4.6) | 0.274 | -155.9 (297.0) | 0.603 | -6.5 (5.4) | 0.472 | -7.1 (3.9) | 0.173 | 56.8 (88.3) | 0.814 | -0.5 (0.8) | 0.671 | -0.4 (1.2) | 0.725 |
| **Males** | Left Crus II | -314.4 (247.2) | 0.768 | 2.7 (2.2) | 0.291 | -7.7 (3.3) | 0.102 | -196.5 (242.9) | 0.603 | 4.2 (4.5) | 0.472 | 1.1 (3.3) | 0.751 | 53.1 (72.1) | 0.814 | -0.6 (0.6) | 0.671 | -0.4 (1.0) | 0.725 |
| | Right Crus II | -239.8 (273.4) | 0.768 | 4.5 (2.4) | 0.241 | -5.2 (3.7) | 0.274 | -181.8 (251.5) | 0.603 | 5.4 (4.6) | 0.472 | 1.4 (3.4) | 0.751 | -28.9 (109.5) | 0.814 | -0.3 (0.9) | 0.745 | -1.1 (1.5) | 0.725 |
| | Left Lobule V | 68.4 (90.0) | 0.450 | 1.5 (0.8) | 0.212 | 1.0 (1.2) | 0.709 | -14.0 (72.6) | 0.858 | -1.4 (1.3) | 0.842 | -0.9 (1.0) | 0.748 | -23.7 (35.4) | 0.687 | -0.1 (0.3) | 0.662 | -0.4 (0.5) | 0.621 |

| | | Baseline | | | | | | Year 1 | | | | | | Baseline→Year 1 | | | | | |
|---|---|---|---|---|---|---|---|---|---|---|---|---|---|---|---|---|---|---|---|
| | | E | | T | | P | | E | | T | | P | | E | | T | | P | |
| **Stage** | **Region** | *b (SE)* | *p* | *b (SE)* | *p* | *b (SE)* | *p* | *b (SE)* | *p* | *b (SE)* | *p* | *b (SE)* | *p* | *b (SE)* | *p* | *b (SE)* | *p* | *b (SE)* | *p* |
| | Right Lobule V | 110.6 (79.4) | 0.322 | 1.1 (0.7) | 0.283 | 1.1 (1.1) | 0.709 | -12.7 (70.6) | 0.858 | 0.0 (1.3) | 0.972 | 0.1 (1.0) | 0.910 | -11.7 (37.7) | 0.758 | -0.4 (0.3) | 0.502 | -0.9 (0.5) | 0.330 |
| | Left Lobule VI | -304.3 (201.9) | 0.322 | -0.6 (1.8) | 0.765 | -0.8 (2.9) | 0.781 | -73.7 (170.0) | 0.858 | -2.5 (3.1) | 0.842 | 2.3 (2.3) | 0.748 | 85.7 (43.6) | 0.227 | 0.2 (0.4) | 0.662 | 0.5 (0.6) | 0.621 |
| | Right Lobule VI | -226.4 (191.2) | 0.322 | 0.5 (1.7) | 0.765 | -1.7 (2.7) | 0.709 | 69.6 (158.2) | 0.858 | -1.3 (2.9) | 0.883 | 1.2 (2.1) | 0.765 | 43.7 (66.5) | 0.687 | -0.8 (0.6) | 0.502 | 0.4 (0.9) | 0.684 |
| | Left Hippocampus | **-221.4 (75.1)** | **0.019*** | -0.5 (0.7) | 0.804 | -1.1 (1.1) | 0.475 | -103.1 (70.9) | 0.308 | -1.3 (1.3) | 0.668 | -0.3 (0.9) | 0.925 | 40.9 (20.7) | 0.329 | 0.3 (0.2) | 0.287 | 0.1 (0.3) | 0.778 |
| | Right Hippocampus | -166.6 (79.1) | 0.059 | -0.3 (0.7) | 0.804 | -0.2 (1.1) | 0.881 | -71.3 (73.2) | 0.504 | -2.2 (1.3) | 0.513 | -0.1 (1.0) | 0.925 | -1.9 (23.4) | 0.934 | 0.0 (0.2) | 0.974 | -0.1 (0.3) | 0.778 |
| | Left Amygdala | -70.4 (31.0) | 0.053 | 0.0 (0.3) | 0.959 | -0.6 (0.4) | 0.475 | -60.6 (27.3) | 0.098 | -0.5 (0.5) | 0.668 | -0.5 (0.4) | 0.925 | -13.1 (18.5) | 0.582 | 0.0 (0.1) | 0.974 | 0.2 (0.2) | 0.778 |
| | Right Amygdala | **-98.5 (34.8)** | **0.019*** | -0.6 (0.3) | 0.563 | -0.8 (0.5) | 0.475 | -76.8 (31.3) | 0.098 | -0.4 (0.6) | 0.688 | -0.3 (0.5) | 0.925 | 20.7 (16.8) | 0.530 | 0.2 (0.1) | 0.287 | 0.4 (0.2) | 0.264 |
| | Left Thalamus | -45.1 (122.2) | 0.718 | 0.5 (1.1) | 0.804 | -1.7 (1.7) | 0.475 | 81.6 (107.7) | 0.543 | -0.8 (2.0) | 0.688 | 0.7 (1.5) | 0.925 | 39.9 (40.7) | 0.530 | 0.1 (0.4) | 0.974 | 0.2 (0.6) | 0.778 |
| | Right Thalamus | 44.2 (121.9) | 0.718 | 1.1 (1.1) | 0.804 | -0.7 (1.7) | 0.839 | 42.6 (106.1) | 0.690 | -0.9 (2.0) | 0.688 | 0.1 (1.4) | 0.925 | 49.7 (52.8) | 0.530 | 0.7 (0.4) | 0.287 | 1.0 (0.7) | 0.487 |
| | Left Lobule IX | 106.3 (97.3) | 0.963 | 0.6 (0.9) | 0.818 | -0.2 (1.4) | 0.944 | -3.4 (80.4) | 0.966 | 1.8 (1.5) | 0.528 | 0.8 (1.1) | 0.635 | -15.2 (37.5) | 0.765 | -0.3 (0.3) | 0.721 | -0.2 (0.5) | 0.792 |
| | Right Lobule IX | 104.1 (106.3) | 0.963 | 1.2 (0.9) | 0.739 | -0.3 (1.5) | 0.944 | 20.0 (83.9) | 0.958 | 2.4 (1.5) | 0.503 | 1.6 (1.1) | 0.574 | 23.9 (33.1) | 0.676 | -0.3 (0.3) | 0.721 | 0.3 (0.4) | 0.792 |
| | Vermis VI | 48.4 (55.8) | 0.963 | 0.6 (0.5) | 0.739 | 0.2 (0.8) | 0.944 | 32.2 (48.1) | 0.958 | 0.8 (0.9) | 0.706 | 1.1 (0.6) | 0.574 | -16.3 (18.2) | 0.676 | 0.0 (0.2) | 0.896 | 0.2 (0.2) | 0.792 |
| | Vermis VII | -6.6 (27.9) | 0.963 | 0.1 (0.3) | 0.818 | -0.4 (0.4) | 0.911 | -12.7 (23.7) | 0.958 | -0.6 (0.4) | 0.503 | -0.3 (0.3) | 0.619 | 8.2 (10.1) | 0.676 | 0.0 (0.1) | 0.822 | 0.1 (0.1) | 0.792 |
| | Vermis VIII | -57.0 (54.4) | 0.963 | 0.4 (0.5) | 0.818 | -1.0 (0.7) | 0.911 | -11.2 (48.4) | 0.958 | 0.5 (0.9) | 0.791 | 0.1 (0.6) | 0.834 | -26.4 (19.9) | 0.676 | -0.1 (0.2) | 0.776 | -0.2 (0.3) | 0.792 |
| | Vermis IX | 1.1 (22.7) | 0.963 | 0.1 (0.2) | 0.818 | -0.4 (0.3) | 0.911 | -3.6 (20.4) | 0.958 | -0.1 (0.4) | 0.901 | 0.1 (0.3) | 0.740 | 2.4 (11.3) | 0.832 | 0.0 (0.1) | 0.822 | 0.1 (0.2) | 0.792 |
| | Vermis X | 2.1 (12.8) | 0.963 | 0.1 (0.1) | 0.818 | -0.2 (0.2) | 0.911 | 12.3 (11.3) | 0.935 | 0.0 (0.2) | 0.901 | 0.2 (0.1) | 0.574 | -2.9 (4.8) | 0.687 | -0.1 (0.0) | 0.627 | 0.0 (0.1) | 0.827 |
| | Total Gray Matter | -2553.4 (6515.9) | 0.963 | 14.2 (58.3) | 0.818 | -37.4 (91.1) | 0.944 | -7924.1 (5683.2) | 0.935 | -67.7 (107.3) | 0.791 | 52.6 (78.6) | 0.635 | 2011.6 (2234.6) | 0.676 | 19.6 (19.2) | 0.721 | 49.9 (29.9) | 0.514 |
| | Cortical Volume | -1114.8 (5618.6) | 0.963 | -15.3 (50.2) | 0.818 | 5.5 (78.4) | 0.944 | -5775.3 (4906.8) | 0.935 | -44.4 (92.3) | 0.791 | 60.8 (67.2) | 0.619 | 2091.4 (2036.7) | 0.676 | 23.4 (17.5) | 0.627 | 47.2 (27.0) | 0.514 |
| | Cerebral White Matter | -3447.8 (6711.6) | 0.963 | 86.0 (59.1) | 0.739 | -67.4 (92.6) | 0.939 | -2669.8 (6077.1) | 0.958 | -156.6 (106.9) | 0.503 | -96.5 (79.0) | 0.574 | 1840.7 (1392.1) | 0.676 | 18.0 (12.0) | 0.627 | -2.8 (19.5) | 0.884 |

*Note.* Reproductive-stage stratified analyses across all region families. With outliers retained. Reproductive stage classified for female participants; hysterectomy ($n$ = 1) and undetermined stage ($n$ = 3) excluded owing to insufficient subgroup size; males shown separately. E = 17β-estradiol; T = testosterone; P = progesterone; *b* = unstandardized coefficient; *SE* = standard error; $n$ = analytic sample per hormone within stage. Post-hoc: all regions FDR-corrected within family. Prediction (Baseline→Year 1) controls for baseline volume. *$p$_FDR < .05, **$p$_FDR < .01, ***$p$_FDR < .001.

**Table S17.** Reproductive stage-stratified hormone-volume associations: Baseline, Year 1, and Baseline-predicting Year 1 models for all regions, without outliers.

| | | Baseline | | | | | | Year 1 | | | | | | Baseline→Year 1 | | | | | |
|---|---|---|---|---|---|---|---|---|---|---|---|---|---|---|---|---|---|---|---|
| | | E | | T | | P | | E | | T | | P | | E | | T | | P | |
| Stage | Region | b (SE) | p | b (SE) | p | b (SE) | p | b (SE) | p | b (SE) | p | b (SE) | p | b (SE) | p | b (SE) | p | b (SE) | p |
| | | *n* = 13 | | *n* = 13 | | *n* = 12 | | *n* = 7 | | *n* = 7 | | *n* = 7 | | *n* = 8 | | *n* = 7 | | *n* = 7 | |
| | Left Crus I | 477.4 (700.4) | 0.730 | 9.3 (14.9) | 0.777 | 0.5 (2.4) | 0.843 | 1185.7 (2549.4) | 0.688 | -51.4 (19.9) | 0.246 | 6.4 (9.7) | 0.861 | 25.0 (116.8) | 0.866 | -2.2 (14.6) | 0.904 | 0.4 (0.0) | 0.088 |
| | Right Crus I | 764.6 (818.5) | 0.730 | 8.7 (15.2) | 0.777 | 2.5 (2.8) | 0.534 | 2493.9 (3562.2) | 0.688 | -79.2 (25.2) | 0.246 | 11.6 (13.6) | 0.861 | -147.4 (317.6) | 0.866 | -10.2 (17.9) | 0.904 | **1.6 (0.0)** | **0.020*** |
| | Left Crus II | -314.6 (842.2) | 0.730 | -0.3 (18.7) | 0.990 | 7.1 (3.0) | 0.208 | 5265.2 (2696.4) | 0.380 | -54.4 (59.7) | 0.458 | 3.6 (18.2) | 0.861 | 505.5 (515.9) | 0.866 | 4.7 (22.9) | 0.904 | -1.3 (1.6) | 0.747 |
| | Right Crus II | -308.5 (863.1) | 0.730 | 14.6 (17.4) | 0.777 | 4.6 (3.7) | 0.492 | 5144.0 (1895.7) | 0.380 | -63.4 (44.8) | 0.391 | 7.0 (15.6) | 0.861 | 80.2 (417.8) | 0.866 | -24.3 (6.4) | 0.652 | -0.4 (2.2) | 0.886 |
| | Left Lobule V | -402.8 (325.2) | 0.334 | -2.5 (7.6) | 0.772 | 2.1 (1.5) | 0.397 | 2109.0 (650.6) | 0.167 | -16.9 (22.1) | 0.944 | 3.4 (6.1) | 0.760 | -30.4 (233.0) | 0.908 | -16.3 (4.0) | 0.521 | -1.2 (0.3) | 0.646 |
| | Right Lobule V | -227.0 (250.8) | 0.392 | 2.3 (5.6) | 0.772 | 1.0 (1.1) | 0.397 | 1475.2 (330.8) | 0.167 | -18.2 (10.9) | 0.944 | 3.1 (3.8) | 0.760 | 68.8 (126.7) | 0.908 | -4.5 (11.8) | 0.827 | -0.0 (0.8) | 0.967 |
| | Left Lobule VI | -760.4 (562.2) | 0.334 | 3.8 (12.5) | 0.772 | 3.0 (2.6) | 0.397 | 1208.9 (2603.2) | 0.688 | 3.3 (42.3) | 0.944 | -6.2 (10.0) | 0.760 | 85.3 (64.8) | 0.908 | 6.3 (2.7) | 0.521 | 0.3 (0.2) | 0.647 |
| | Right Lobule VI | -1110.7 (545.0) | 0.304 | -7.5 (14.5) | 0.772 | 2.8 (2.9) | 0.397 | 2251.4 (2664.3) | 0.649 | -14.1 (46.9) | 0.944 | -4.2 (12.0) | 0.760 | 156.3 (376.4) | 0.908 | 7.3 (26.0) | 0.827 | -0.4 (1.3) | 0.967 |
| | Left Hippocampus | -11.9 (243.7) | 0.962 | 2.7 (4.8) | 0.583 | 1.4 (0.9) | 0.258 | 778.7 (447.2) | 0.302 | -4.8 (10.4) | 0.955 | 0.5 (2.8) | 0.934 | 74.6 (75.0) | 0.637 | 7.4 (5.5) | 0.570 | 0.3 (0.3) | 0.774 |
| | Right Hippocampus | -100.0 (256.4) | 0.848 | 6.0 (4.7) | 0.278 | 1.0 (1.1) | 0.446 | 845.4 (505.9) | 0.302 | -8.1 (10.7) | 0.955 | 0.3 (3.1) | 0.934 | 106.9 (97.9) | 0.637 | 2.4 (4.0) | 0.657 | 0.1 (0.2) | 0.791 |
| | Left Amygdala | 82.0 (66.1) | 0.848 | 2.8 (1.2) | 0.278 | -0.2 (0.3) | 0.631 | -155.2 (97.4) | 0.302 | -0.2 (2.3) | 0.955 | -0.4 (0.5) | 0.934 | -51.2 (13.7) | 0.238 | -2.5 (2.3) | 0.570 | -0.1 (0.1) | 0.774 |
| Reproductive | Right Amygdala | -32.0 (51.3) | 0.848 | 1.2 (0.9) | 0.278 | -0.3 (0.2) | 0.258 | -154.5 (131.8) | 0.362 | 2.0 (2.2) | 0.955 | 0.1 (0.7) | 0.934 | 106.9 (32.1) | 0.238 | 7.3 (1.4) | 0.570 | 0.2 (0.6) | 0.791 |
| | Left Thalamus | 277.7 (499.8) | 0.848 | 14.7 (11.0) | 0.278 | 4.7 (1.6) | 0.064 | 1828.1 (931.3) | 0.302 | -1.6 (24.6) | 0.955 | 6.5 (4.4) | 0.851 | -88.6 (267.7) | 0.772 | 15.8 (11.4) | 0.570 | 3.4 (2.4) | 0.774 |
| | Right Thalamus | 345.6 (445.2) | 0.848 | 13.8 (9.3) | 0.278 | 4.2 (1.4) | 0.064 | 1753.2 (425.6) | 0.302 | -14.8 (17.3) | 0.955 | 6.6 (2.4) | 0.670 | 92.8 (206.1) | 0.772 | 17.7 (6.1) | 0.570 | 4.7 (3.9) | 0.774 |
| | Left Lobule IX | -273.0 (427.0) | 0.873 | 10.3 (9.3) | 0.550 | 3.5 (1.6) | 0.435 | 1646.9 (1107.4) | 0.829 | 0.9 (24.8) | 0.974 | 2.7 (6.1) | 0.997 | 118.7 (52.3) | 0.504 | 4.7 (2.3) | 0.572 | 0.5 (0.9) | 0.914 |
| | Right Lobule IX | -270.8 (512.3) | 0.873 | 15.0 (10.2) | 0.508 | 4.0 (2.0) | 0.435 | 2271.1 (794.1) | 0.829 | -11.9 (26.4) | 0.974 | 4.2 (6.5) | 0.997 | 93.3 (129.9) | 0.869 | -4.2 (4.8) | 0.686 | -0.2 (0.8) | 0.914 |
| | Vermis VI | -137.6 (114.6) | 0.873 | -0.9 (3.3) | 0.798 | -0.4 (0.6) | 0.711 | 196.6 (141.0) | 0.829 | -2.8 (2.3) | 0.686 | -0.0 (0.8) | 0.997 | 13.9 (23.1) | 0.869 | -2.3 (2.7) | 0.686 | -0.0 (0.1) | 0.914 |
| | Vermis VII | -32.5 (82.5) | 0.880 | 1.2 (1.7) | 0.675 | 0.1 (0.4) | 0.795 | -82.0 (187.7) | 0.881 | 4.0 (1.2) | 0.266 | 0.0 (0.8) | 0.997 | 42.6 (9.8) | 0.331 | 3.7 (6.9) | 0.686 | 0.0 (0.1) | 0.914 |
| | Vermis VIII | 18.4 (138.7) | 0.897 | 5.4 (2.9) | 0.508 | 0.7 (0.6) | 0.711 | 290.2 (284.3) | 0.829 | -5.2 (4.0) | 0.686 | -0.3 (1.4) | 0.997 | 115.9 (31.4) | 0.331 | 5.5 (0.4) | 0.232 | 0.2 (0.1) | 0.914 |
| | Vermis IX | -66.4 (77.7) | 0.873 | 3.2 (1.3) | 0.433 | -0.0 (0.4) | 0.973 | 20.0 (230.8) | 0.939 | -1.6 (3.4) | 0.974 | -0.9 (0.7) | 0.997 | 1.8 (29.0) | 0.957 | -1.5 (0.9) | 0.604 | 0.0 (0.1) | 0.914 |
| | Vermis X | -22.1 (38.3) | 0.873 | 1.1 (0.8) | 0.508 | 0.2 (0.2) | 0.711 | 73.9 (96.0) | 0.870 | -0.2 (1.7) | 0.974 | -0.1 (0.4) | 0.997 | -1.6 (19.8) | 0.957 | 0.9 (1.7) | 0.686 | -0.1 (0.1) | 0.914 |
| | Total Gray Matter | -7445.4 (11681.5) | 0.873 | -80.9 (127.0) | 0.675 | 16.9 (28.3) | 0.711 | -28721.4 (52635.1) | 0.881 | 1179.8 (253.4) | 0.216 | -36.6 (224.1) | 0.997 | 16408.3 (10130.3) | 0.617 | 1563.3 (213.3) | 0.288 | 31.3 (71.6) | 0.914 |
| | Cortical Volume | -3852.1 (15666.4) | 0.897 | -247.2 (238.5) | 0.550 | -45.0 (53.0) | 0.711 | -67467.4 (56465.6) | 0.829 | 1546.8 (330.0) | 0.216 | -78.8 (290.3) | 0.997 | 11207.0 (10495.8) | 0.795 | 1379.9 (52.5) | 0.232 | 48.9 (58.1) | 0.914 |
| | Cerebral White Matter | 21500.1 (21401.8) | 0.873 | -246.8 (461.6) | 0.675 | 68.2 (98.5) | 0.711 | 11842.4 (69066.7) | 0.939 | -90.4 (1073.3) | 0.974 | 312.8 (168.7) | 0.997 | 1196.8 (9395.9) | 0.957 | 606.8 (287.5) | 0.572 | 26.9 (22.7) | 0.914 |
| | | *n* = 14 | | *n* = 14 | | *n* = 14 | | *n* = 10 | | *n* = 10 | | *n* = 10 | | *n* = 10 | | *n* = 10 | | *n* = 10 | |

| | | Baseline | | | | | | Year 1 | | | | | | Baseline→Year 1 | | | | | |
|---|---|---|---|---|---|---|---|---|---|---|---|---|---|---|---|---|---|---|---|
| | | E | | T | | P | | E | | T | | P | | E | | T | | P | |
| Stage | Region | b (SE) | p | b (SE) | p | b (SE) | p | b (SE) | p | b (SE) | p | b (SE) | p | b (SE) | p | b (SE) | p | b (SE) | p |
| Early Postmenopausal | Left Crus I | -516.3 (745.5) | 0.508 | -5.6 (14.8) | 0.949 | -8.1 (9.0) | 0.390 | -86.6 (1887.2) | 0.965 | 4.5 (22.8) | 0.851 | -6.6 (8.9) | 0.547 | -122.0 (307.6) | 0.712 | 1.3 (6.6) | 0.855 | -3.8 (3.9) | 0.648 |
| | Right Crus I | -571.0 (827.8) | 0.508 | 0.1 (16.6) | 0.993 | -13.5 (9.4) | 0.390 | -1723.6 (1931.1) | 0.826 | 13.2 (24.5) | 0.851 | -14.2 (8.2) | 0.424 | 440.9 (266.9) | 0.348 | 10.9 (4.6) | 0.159 | 6.8 (4.9) | 0.648 |
| | Left Crus II | -931.2 (543.3) | 0.421 | -21.4 (9.9) | 0.234 | -8.5 (7.1) | 0.390 | 1075.4 (655.4) | 0.647 | -2.6 (9.8) | 0.851 | 4.9 (3.4) | 0.424 | -693.6 (367.9) | 0.348 | -19.1 (8.1) | 0.159 | 2.3 (6.6) | 0.744 |
| | Right Crus II | -935.9 (694.4) | 0.421 | -20.8 (13.1) | 0.294 | -8.8 (8.8) | 0.390 | 556.5 (1163.1) | 0.870 | -26.6 (8.2) | 0.090 | 3.7 (5.7) | 0.547 | -326.8 (305.8) | 0.461 | -6.5 (6.8) | 0.528 | 3.2 (4.2) | 0.648 |
| | Left Lobule V | -425.1 (211.9) | 0.152 | -8.0 (4.2) | 0.257 | **-8.3 (1.4)** | **0.001***** | -511.7 (557.7) | 0.401 | -10.4 (5.6) | 0.257 | -4.0 (2.4) | 0.370 | 35.2 (126.5) | 0.965 | 1.4 (2.4) | 0.650 | 2.2 (3.6) | 0.778 |
| | Right Lobule V | -344.9 (158.2) | 0.152 | -5.6 (3.3) | 0.257 | **-6.2 (1.2)** | **0.001***** | -455.4 (424.9) | 0.401 | -8.1 (4.4) | 0.257 | -3.0 (1.9) | 0.370 | 9.0 (193.7) | 0.965 | 1.7 (3.5) | 0.650 | 0.9 (3.9) | 0.825 |
| | Left Lobule VI | -611.9 (579.7) | 0.319 | -5.8 (11.8) | 0.680 | -5.9 (7.3) | 0.437 | -1614.2 (1301.4) | 0.401 | -6.7 (17.8) | 0.899 | -3.1 (7.3) | 0.686 | 88.5 (114.5) | 0.965 | 2.7 (2.0) | 0.494 | 1.0 (1.3) | 0.778 |
| | Right Lobule VI | -759.8 (656.2) | 0.319 | -5.8 (13.6) | 0.680 | -8.3 (8.1) | 0.437 | -2261.5 (1312.0) | 0.401 | -2.7 (20.0) | 0.899 | -5.4 (7.9) | 0.686 | 288.5 (113.2) | 0.254 | 4.8 (2.6) | 0.494 | 2.5 (1.7) | 0.778 |
| | Left Hippocampus | -3.7 (179.8) | 0.984 | -1.5 (3.5) | 0.675 | -2.8 (2.0) | 0.594 | 65.9 (402.0) | 0.876 | 3.8 (4.6) | 0.527 | -0.5 (2.0) | 0.964 | -4.0 (47.1) | 0.937 | -0.2 (1.0) | 0.839 | 0.9 (0.5) | 0.641 |
| | Right Hippocampus | -46.7 (172.7) | 0.951 | -2.5 (3.3) | 0.675 | -2.9 (1.9) | 0.594 | -123.9 (311.7) | 0.876 | -3.1 (3.6) | 0.527 | -1.1 (1.5) | 0.964 | -17.4 (33.9) | 0.936 | -0.9 (0.6) | 0.441 | 0.5 (0.6) | 0.641 |
| | Left Amygdala | 63.0 (45.1) | 0.588 | 0.5 (1.0) | 0.675 | -0.3 (0.6) | 0.801 | -114.9 (146.2) | 0.876 | 2.7 (1.4) | 0.375 | -0.7 (0.7) | 0.964 | 74.2 (29.9) | 0.409 | **1.8 (0.3)** | **0.026*** | 0.3 (0.6) | 0.641 |
| | Right Amygdala | 94.6 (66.6) | 0.588 | 1.0 (1.4) | 0.675 | 0.3 (0.9) | 0.801 | -38.2 (121.7) | 0.876 | 2.4 (1.0) | 0.363 | -0.1 (0.6) | 0.964 | 43.5 (34.0) | 0.542 | 0.5 (0.8) | 0.839 | -0.4 (0.5) | 0.641 |
| | Left Thalamus | 353.6 (324.8) | 0.609 | 4.9 (6.5) | 0.675 | 1.1 (4.2) | 0.801 | -205.6 (859.7) | 0.876 | 13.1 (8.7) | 0.382 | 0.2 (4.3) | 0.964 | 19.8 (66.1) | 0.936 | -0.3 (1.3) | 0.839 | 0.4 (0.8) | 0.641 |
| | Right Thalamus | 135.8 (207.9) | 0.795 | 2.5 (4.1) | 0.675 | 1.9 (2.5) | 0.801 | -84.1 (458.8) | 0.876 | 2.4 (5.5) | 0.675 | 1.2 (2.2) | 0.964 | -104.5 (74.6) | 0.542 | -2.6 (1.3) | 0.357 | -0.8 (1.1) | 0.641 |
| | Left Lobule IX | -317.9 (193.8) | 0.461 | -6.8 (3.7) | 0.616 | -2.7 (2.5) | 0.797 | 635.7 (577.9) | 0.536 | -7.0 (7.2) | 0.881 | 0.9 (3.2) | 0.784 | 129.2 (76.0) | 0.445 | 2.9 (1.4) | 0.384 | 1.7 (0.9) | 0.508 |
| | Right Lobule IX | -334.7 (251.9) | 0.461 | -7.3 (4.8) | 0.616 | -2.6 (3.3) | 0.797 | 908.8 (618.9) | 0.536 | -4.6 (8.7) | 0.969 | 1.6 (3.6) | 0.784 | 95.0 (55.6) | 0.445 | 2.5 (0.9) | 0.231 | 1.2 (0.7) | 0.508 |
| | Vermis VI | -3.8 (109.1) | 0.973 | 0.3 (2.1) | 0.956 | 1.2 (1.3) | 0.797 | 93.0 (253.7) | 0.729 | 2.6 (2.9) | 0.881 | 0.7 (1.2) | 0.784 | -29.3 (18.1) | 0.445 | -0.6 (0.4) | 0.384 | 0.0 (0.3) | 0.995 |
| | Vermis VII | -13.8 (51.1) | 0.881 | -0.1 (1.0) | 0.956 | 0.0 (0.6) | 0.994 | 202.2 (124.2) | 0.536 | 0.4 (1.9) | 0.985 | 0.5 (0.7) | 0.784 | -37.6 (29.2) | 0.445 | -0.8 (0.6) | 0.446 | 0.0 (0.4) | 0.995 |
| | Vermis VIII | -192.1 (103.8) | 0.461 | -3.1 (2.1) | 0.616 | -2.7 (1.2) | 0.496 | -153.3 (293.4) | 0.693 | 0.1 (3.7) | 0.985 | -1.4 (1.4) | 0.784 | 45.7 (14.0) | 0.308 | **1.1 (0.2)** | **0.027*** | -0.1 (0.4) | 0.995 |
| | Vermis IX | -55.1 (47.5) | 0.461 | -0.6 (1.0) | 0.803 | -1.0 (0.5) | 0.496 | -77.6 (105.8) | 0.693 | -1.7 (1.1) | 0.881 | -0.7 (0.5) | 0.784 | -39.0 (30.1) | 0.445 | -0.4 (0.7) | 0.774 | -0.8 (0.4) | 0.508 |
| | Vermis X | -28.0 (23.0) | 0.461 | -0.4 (0.5) | 0.645 | -0.2 (0.3) | 0.797 | 141.3 (60.8) | 0.536 | -0.1 (1.1) | 0.985 | 0.5 (0.4) | 0.784 | 8.6 (9.9) | 0.574 | 0.0 (0.2) | 0.930 | 0.1 (0.1) | 0.771 |
| | Total Gray Matter | 4543.3 (11627.3) | 0.881 | 46.8 (228.2) | 0.956 | -86.5 (140.5) | 0.797 | -26331.8 (23244.1) | 0.536 | -247.7 (295.8) | 0.881 | -102.0 (121.8) | 0.784 | 3942.3 (4820.6) | 0.574 | 128.0 (89.2) | 0.446 | 7.6 (67.4) | 0.995 |
| | Cortical Volume | 13398.0 (11238.5) | 0.461 | 205.2 (225.8) | 0.645 | 58.3 (146.7) | 0.875 | -16993.9 (29878.5) | 0.693 | -161.2 (366.7) | 0.969 | -44.5 (152.5) | 0.784 | 2877.8 (4454.2) | 0.615 | 91.3 (96.2) | 0.566 | -31.1 (59.8) | 0.995 |
| | Cerebral White Matter | 12719.4 (12782.1) | 0.494 | 270.7 (246.7) | 0.645 | -15.3 (164.7) | 0.994 | -40324.7 (32929.7) | 0.536 | 613.9 (362.9) | 0.881 | -152.3 (174.5) | 0.784 | 2399.3 (5002.9) | 0.657 | 10.0 (106.9) | 0.930 | 11.4 (63.8) | 0.995 |
| | | *n* = 35 | | *n* = 35 | | *n* = 34 | | *n* = 26 | | *n* = 26 | | *n* = 26 | | *n* = 26 | | *n* = 26 | | *n* = 25 | |
| Late Postmenopausal | Left Crus I | -569.4 (311.0) | 0.154 | -8.7 (5.5) | 0.245 | -5.6 (5.8) | 0.806 | -492.7 (423.8) | 0.387 | -6.6 (7.1) | 0.359 | -6.2 (3.2) | 0.137 | -26.8 (110.4) | 0.905 | 0.9 (1.9) | 0.633 | 3.4 (2.1) | 0.486 |
| | Right Crus I | -767.0 (410.3) | 0.154 | -11.8 (7.2) | 0.245 | -6.6 (7.8) | 0.806 | -728.8 (609.9) | 0.387 | -13.7 (10.0) | 0.355 | -8.8 (4.6) | 0.137 | 15.2 (126.0) | 0.905 | 1.8 (2.1) | 0.633 | 2.6 (2.4) | 0.591 |

| Stage | Region | Baseline | | | | | | Year 1 | | | | | | Baseline→Year 1 | | | | | |
|---|---|---|---|---|---|---|---|---|---|---|---|---|---|---|---|---|---|---|---|
| | | E | | T | | P | | E | | T | | P | | E | | T | | P | |
| Stage | Region | b (SE) | p | b (SE) | p | b (SE) | p | b (SE) | p | b (SE) | p | b (SE) | p | b (SE) | p | b (SE) | p | b (SE) | p |
| | Left Crus II | 64.9 (404.9) | 0.893 | 4.0 (7.0) | 0.568 | -0.7 (7.4) | 0.929 | 538.1 (533.2) | 0.387 | 10.6 (8.7) | 0.355 | -0.1 (4.3) | 0.978 | 58.3 (92.6) | 0.905 | 1.2 (1.6) | 0.633 | 0.3 (1.9) | 0.860 |
| | Right Crus II | -52.5 (386.0) | 0.893 | 6.4 (6.6) | 0.456 | 0.6 (7.0) | 0.929 | 422.6 (478.2) | 0.387 | 8.9 (7.8) | 0.355 | -0.2 (3.8) | 0.978 | 56.1 (79.4) | 0.905 | 0.8 (1.4) | 0.633 | -0.9 (1.6) | 0.777 |
| | Left Lobule V | -82.5 (89.0) | 0.777 | 0.2 (1.6) | 0.904 | -1.5 (1.6) | 0.602 | -93.9 (119.6) | 0.811 | 1.0 (2.0) | 0.620 | -1.0 (0.9) | 0.938 | -0.8 (27.0) | 0.976 | 0.5 (0.5) | 0.539 | 0.7 (0.5) | 0.377 |
| | Right Lobule V | -54.5 (98.2) | 0.777 | 1.3 (1.7) | 0.904 | -1.4 (1.8) | 0.602 | -82.5 (137.4) | 0.811 | 2.5 (2.2) | 0.620 | -0.2 (1.1) | 0.938 | 35.3 (28.5) | 0.860 | 0.9 (0.5) | 0.266 | 0.5 (0.6) | 0.457 |
| | Left Lobule VI | 175.9 (235.1) | 0.777 | 2.3 (4.1) | 0.904 | 0.6 (4.1) | 0.881 | -62.8 (346.2) | 0.858 | 4.2 (5.6) | 0.620 | 0.2 (2.7) | 0.938 | 5.8 (64.6) | 0.976 | 0.2 (1.1) | 0.874 | -1.7 (1.2) | 0.377 |
| | Right Lobule VI | -1.0 (231.9) | 0.996 | -0.7 (4.0) | 0.904 | -4.7 (4.0) | 0.602 | 165.1 (317.4) | 0.811 | 3.9 (5.2) | 0.620 | 1.0 (2.5) | 0.938 | 51.1 (63.4) | 0.860 | 0.7 (1.1) | 0.719 | -1.0 (1.0) | 0.422 |
| | Left Hippocampus | 44.0 (96.6) | 0.652 | -0.5 (1.7) | 0.916 | -1.5 (1.7) | 0.644 | -140.4 (118.8) | 0.504 | -1.1 (2.0) | 0.880 | 0.0 (1.0) | 0.960 | -34.3 (21.0) | 0.708 | -0.1 (0.4) | 0.887 | -0.6 (0.4) | 0.910 |
| | Right Hippocampus | 123.2 (92.7) | 0.498 | 1.2 (1.6) | 0.702 | 1.4 (1.7) | 0.644 | -56.2 (125.9) | 0.792 | 0.5 (2.1) | 0.880 | 0.1 (1.0) | 0.960 | -15.0 (21.1) | 0.733 | 0.3 (0.4) | 0.887 | -0.3 (0.4) | 0.910 |
| | Left Amygdala | -24.0 (45.6) | 0.652 | -0.9 (0.8) | 0.474 | -1.4 (0.8) | 0.526 | -25.7 (53.8) | 0.792 | -0.1 (0.9) | 0.880 | 0.3 (0.4) | 0.666 | 19.2 (19.3) | 0.733 | 0.5 (0.3) | 0.760 | 0.3 (0.4) | 0.910 |
| | Right Amygdala | 19.5 (40.9) | 0.652 | -0.0 (0.7) | 0.956 | 0.3 (0.7) | 0.681 | 12.6 (54.8) | 0.820 | 0.4 (0.9) | 0.880 | 0.5 (0.4) | 0.451 | -13.2 (18.7) | 0.733 | -0.1 (0.3) | 0.887 | 0.0 (0.3) | 0.910 |
| | Left Thalamus | 158.7 (135.0) | 0.498 | 2.8 (2.3) | 0.474 | 1.4 (2.4) | 0.678 | 196.9 (147.2) | 0.504 | 3.8 (2.4) | 0.395 | 2.6 (1.1) | 0.079 | -10.5 (69.1) | 0.881 | 0.2 (1.1) | 0.887 | 0.6 (1.5) | 0.910 |
| | Right Thalamus | 175.6 (131.5) | 0.498 | 4.3 (2.2) | 0.367 | 2.7 (2.3) | 0.644 | 183.4 (155.8) | 0.504 | 4.2 (2.5) | 0.395 | 2.7 (1.1) | 0.079 | -38.3 (74.2) | 0.733 | -0.2 (1.3) | 0.887 | -0.5 (1.7) | 0.910 |
| | Left Lobule IX | -17.1 (141.1) | 0.904 | -1.6 (2.4) | 0.652 | 1.3 (2.6) | 0.829 | 98.2 (171.2) | 0.818 | 2.7 (2.8) | 0.772 | -0.2 (1.4) | 0.971 | -25.5 (25.8) | 0.621 | -0.5 (0.4) | 0.732 | 0.5 (0.6) | 0.721 |
| | Right Lobule IX | -22.8 (131.3) | 0.904 | -0.9 (2.3) | 0.765 | 0.5 (2.4) | 0.845 | 28.9 (169.2) | 0.873 | 1.9 (2.8) | 0.772 | -0.7 (1.3) | 0.843 | -87.6 (29.2) | 0.070 | -0.7 (0.6) | 0.732 | -0.7 (0.7) | 0.721 |
| | Vermis VI | -116.6 (64.1) | 0.148 | 0.9 (1.2) | 0.652 | -0.7 (1.2) | 0.829 | -85.1 (98.1) | 0.659 | -0.7 (1.6) | 0.772 | -1.1 (0.7) | 0.354 | 5.7 (21.1) | 0.879 | 0.2 (0.3) | 0.790 | 0.2 (0.4) | 0.775 |
| | Vermis VII | -44.5 (31.1) | 0.234 | -0.8 (0.5) | 0.628 | -0.8 (0.5) | 0.712 | -57.5 (42.6) | 0.639 | -0.5 (0.7) | 0.772 | -0.3 (0.3) | 0.689 | 1.5 (11.1) | 0.892 | -0.2 (0.2) | 0.732 | -0.4 (0.2) | 0.563 |
| | Vermis VIII | -120.6 (62.4) | 0.148 | -1.3 (1.1) | 0.652 | -0.5 (1.2) | 0.829 | -97.2 (94.8) | 0.659 | -1.5 (1.6) | 0.772 | -1.6 (0.7) | 0.119 | -10.0 (19.9) | 0.774 | 0.2 (0.3) | 0.790 | 0.0 (0.4) | 0.927 |
| | Vermis IX | **-84.3 (24.8)** | **0.019*** | -0.8 (0.5) | 0.628 | -0.8 (0.5) | 0.712 | -64.3 (33.4) | 0.639 | -0.8 (0.6) | 0.772 | -0.6 (0.3) | 0.119 | -12.2 (10.9) | 0.621 | 0.0 (0.2) | 0.790 | 0.1 (0.2) | 0.775 |
| | Vermis X | -26.0 (14.8) | 0.148 | -0.2 (0.3) | 0.652 | -0.1 (0.3) | 0.845 | -35.0 (22.5) | 0.639 | -0.3 (0.4) | 0.772 | -0.4 (0.2) | 0.125 | -9.9 (5.6) | 0.471 | -0.1 (0.1) | 0.732 | -0.0 (0.1) | 0.927 |
| | Total Gray Matter | -10807.1 (5676.9) | 0.148 | -83.4 (103.0) | 0.652 | -130.3 (108.1) | 0.712 | -1074.9 (6636.9) | 0.873 | 53.2 (108.9) | 0.772 | -0.0 (52.2) | 1.000 | 3151.6 (3455.9) | 0.621 | 28.2 (58.9) | 0.790 | 97.4 (65.3) | 0.563 |
| | Cortical Volume | -8612.8 (4717.3) | 0.148 | -105.4 (84.0) | 0.652 | -97.6 (89.5) | 0.712 | -1285.5 (6244.7) | 0.873 | 2.3 (103.1) | 0.982 | 19.6 (49.0) | 0.866 | 2805.8 (2916.8) | 0.621 | 14.2 (49.9) | 0.790 | 78.8 (55.1) | 0.563 |
| | Cerebral White Matter | 7517.8 (6591.2) | 0.329 | 12.1 (116.7) | 0.918 | -76.5 (123.3) | 0.829 | -8224.6 (8906.6) | 0.659 | 78.6 (148.9) | 0.772 | 58.7 (70.3) | 0.689 | -2070.3 (2629.1) | 0.629 | 33.2 (44.9) | 0.790 | -29.5 (53.5) | 0.775 |
| | | *n* = 59 | | *n* = 58 | | *n* = 57 | | *n* = 39 | | *n* = 38 | | *n* = 38 | | *n* = 39 | | *n* = 39 | | *n* = 39 | |
| Males | Left Crus I | 214.9 (392.3) | 0.586 | 3.1 (4.1) | 0.536 | -0.2 (5.4) | 0.968 | -285.5 (504.8) | 0.774 | -3.2 (5.8) | 0.591 | -6.5 (4.8) | 0.380 | -93.1 (105.7) | 0.578 | -1.2 (1.3) | 0.555 | -2.9 (1.4) | 0.091 |
| | Right Crus I | 320.4 (356.8) | 0.586 | 2.4 (3.8) | 0.536 | 1.2 (4.9) | 0.968 | -136.8 (473.5) | 0.774 | -6.1 (5.3) | 0.482 | -6.0 (4.5) | 0.380 | -90.5 (109.7) | 0.578 | **-3.6 (1.2)** | **0.017*** | -3.1 (1.4) | 0.091 |
| | Left Crus II | -268.6 (301.0) | 0.586 | 5.9 (3.1) | 0.124 | **-11.3 (3.9)** | **0.021*** | 188.7 (434.6) | 0.774 | 4.6 (5.0) | 0.482 | 1.5 (4.3) | 0.719 | 52.9 (94.0) | 0.578 | -1.0 (1.2) | 0.555 | -1.3 (1.4) | 0.355 |
| | Right Crus II | -227.3 (334.7) | 0.586 | 7.5 (3.4) | 0.117 | -7.2 (4.4) | 0.222 | 196.9 (454.0) | 0.774 | 5.1 (5.2) | 0.482 | 2.0 (4.4) | 0.719 | -96.0 (142.5) | 0.578 | -0.9 (1.8) | 0.601 | -2.9 (2.0) | 0.202 |

| | | Baseline | | | | | | Year 1 | | | | | | Baseline→Year 1 | | | | | |
|---|---|---|---|---|---|---|---|---|---|---|---|---|---|---|---|---|---|---|---|
| | | E | | T | | P | | E | | T | | P | | E | | T | | P | |
| **Stage** | **Region** | b (SE) | p | b (SE) | p | b (SE) | p | b (SE) | p | b (SE) | p | b (SE) | p | b (SE) | p | b (SE) | p | b (SE) | p |
| | Left Lobule V | -25.9 (107.7) | 0.811 | 1.4 (1.1) | 0.498 | -0.2 (1.5) | 0.871 | -178.6 (120.7) | 0.592 | -2.8 (1.4) | 0.176 | -2.2 (1.2) | 0.271 | -51.9 (44.6) | 0.678 | -0.1 (0.6) | 0.833 | -0.5 (0.6) | 0.465 |
| | Right Lobule V | 111.3 (94.8) | 0.491 | 1.1 (1.0) | 0.498 | 0.4 (1.3) | 0.871 | -32.2 (122.9) | 0.968 | -1.0 (1.4) | 0.531 | -1.1 (1.2) | 0.684 | -43.3 (44.7) | 0.678 | -0.8 (0.5) | 0.555 | -1.4 (0.6) | 0.069 |
| | Left Lobule VI | -206.5 (241.2) | 0.528 | 0.6 (2.6) | 0.805 | -0.5 (3.3) | 0.871 | 12.0 (295.9) | 0.968 | -3.1 (3.3) | 0.531 | 1.8 (2.8) | 0.684 | 33.1 (53.4) | 0.719 | -0.4 (0.7) | 0.785 | -0.8 (0.7) | 0.393 |
| | Right Lobule VI | -338.4 (233.5) | 0.491 | 2.2 (2.5) | 0.498 | -2.9 (3.2) | 0.871 | -201.9 (281.6) | 0.956 | -2.1 (3.3) | 0.531 | -1.1 (2.8) | 0.684 | 0.7 (79.5) | 0.993 | -1.1 (1.0) | 0.555 | -1.2 (1.1) | 0.393 |
| | Left Hippocampus | -215.8 (90.3) | 0.122 | 0.3 (1.0) | 0.989 | -1.5 (1.3) | 0.545 | -59.6 (126.6) | 0.980 | -1.3 (1.4) | 0.645 | -0.9 (1.2) | 0.686 | 30.7 (26.2) | 0.748 | 0.1 (0.3) | 0.839 | -0.1 (0.4) | 0.829 |
| | Right Hippocampus | -178.2 (97.1) | 0.216 | 0.0 (1.1) | 0.989 | -0.3 (1.4) | 0.946 | -43.7 (133.2) | 0.980 | -2.7 (1.4) | 0.373 | -0.6 (1.2) | 0.686 | -13.2 (30.2) | 0.882 | -0.1 (0.4) | 0.839 | -0.6 (0.4) | 0.801 |
| | Left Amygdala | -52.4 (37.7) | 0.256 | 0.2 (0.4) | 0.989 | -0.5 (0.5) | 0.545 | -45.5 (50.2) | 0.980 | -0.4 (0.6) | 0.645 | -0.6 (0.5) | 0.686 | -28.6 (21.7) | 0.748 | 0.1 (0.3) | 0.839 | -0.1 (0.3) | 0.829 |
| | Right Amygdala | -61.1 (41.5) | 0.256 | -0.2 (0.4) | 0.989 | -0.8 (0.6) | 0.545 | -52.7 (54.1) | 0.980 | -0.2 (0.6) | 0.794 | -0.5 (0.5) | 0.686 | 14.6 (18.9) | 0.882 | 0.2 (0.2) | 0.839 | 0.2 (0.3) | 0.829 |
| | Left Thalamus | -153.8 (147.4) | 0.362 | 0.3 (1.6) | 0.989 | -1.9 (2.0) | 0.545 | 4.9 (192.1) | 0.980 | -1.4 (2.2) | 0.645 | 1.0 (1.9) | 0.686 | -17.6 (51.6) | 0.882 | -0.9 (0.6) | 0.839 | -0.8 (0.7) | 0.801 |
| | Right Thalamus | -48.9 (145.8) | 0.739 | 1.1 (1.5) | 0.989 | -0.1 (2.0) | 0.946 | -37.3 (184.3) | 0.980 | -1.7 (2.1) | 0.645 | 0.7 (1.8) | 0.686 | -2.5 (65.9) | 0.970 | 0.6 (0.8) | 0.839 | 0.1 (0.9) | 0.924 |
| | Left Lobule IX | 133.6 (117.9) | 0.821 | 0.1 (1.3) | 0.933 | -0.5 (1.6) | 0.865 | 158.9 (142.6) | 0.746 | 1.6 (1.6) | 0.730 | 1.1 (1.4) | 0.673 | -49.2 (48.1) | 0.573 | -0.7 (0.6) | 0.616 | -1.1 (0.6) | 0.484 |
| | Right Lobule IX | 139.9 (129.5) | 0.821 | 1.0 (1.4) | 0.933 | -0.5 (1.8) | 0.865 | 197.6 (149.0) | 0.746 | 2.2 (1.7) | 0.681 | 2.3 (1.4) | 0.673 | -4.8 (40.6) | 0.907 | -0.7 (0.5) | 0.616 | -0.6 (0.5) | 0.522 |
| | Vermis VI | 16.2 (67.0) | 0.993 | 0.7 (0.7) | 0.933 | -0.3 (0.9) | 0.865 | 67.9 (81.8) | 0.746 | 0.1 (0.9) | 0.872 | 0.6 (0.8) | 0.673 | -30.6 (21.0) | 0.573 | -0.2 (0.3) | 0.616 | -0.3 (0.3) | 0.512 |
| | Vermis VII | -2.9 (33.5) | 0.993 | 0.2 (0.4) | 0.933 | -0.2 (0.5) | 0.865 | -18.0 (41.8) | 0.746 | -0.7 (0.5) | 0.623 | -0.6 (0.4) | 0.673 | 12.5 (13.0) | 0.573 | -0.1 (0.2) | 0.623 | 0.1 (0.2) | 0.678 |
| | Vermis VIII | -66.4 (67.3) | 0.821 | 0.8 (0.7) | 0.933 | -1.4 (0.9) | 0.715 | 32.5 (88.3) | 0.746 | 0.6 (1.0) | 0.786 | 0.4 (0.8) | 0.677 | -40.8 (25.0) | 0.573 | -0.2 (0.3) | 0.638 | -0.7 (0.3) | 0.372 |
| | Vermis IX | 3.7 (27.5) | 0.993 | 0.3 (0.3) | 0.933 | -0.5 (0.4) | 0.715 | -15.4 (36.6) | 0.746 | -0.2 (0.4) | 0.786 | -0.2 (0.4) | 0.677 | -1.8 (13.6) | 0.907 | -0.0 (0.2) | 0.883 | -0.2 (0.2) | 0.512 |
| | Vermis X | -1.4 (15.1) | 0.993 | 0.1 (0.2) | 0.933 | -0.2 (0.2) | 0.715 | 18.7 (20.2) | 0.746 | -0.1 (0.2) | 0.786 | 0.2 (0.2) | 0.673 | -7.1 (5.1) | 0.573 | -0.1 (0.1) | 0.616 | -0.1 (0.1) | 0.512 |
| | Total Gray Matter | -613.5 (8011.5) | 0.993 | 30.4 (84.6) | 0.933 | -33.0 (110.0) | 0.865 | -8421.4 (10348.4) | 0.746 | -109.6 (119.3) | 0.730 | 42.9 (102.0) | 0.677 | 1067.3 (2522.4) | 0.844 | 16.7 (31.2) | 0.663 | -19.9 (35.6) | 0.678 |
| | Cortical Volume | 65.4 (6948.6) | 0.993 | -21.5 (73.4) | 0.933 | 10.9 (95.3) | 0.909 | -5587.1 (8973.5) | 0.746 | -75.9 (103.6) | 0.782 | 64.0 (87.8) | 0.673 | 2756.9 (2404.2) | 0.573 | 28.3 (30.4) | 0.616 | 14.3 (34.3) | 0.678 |
| | Cerebral White Matter | -11641.4 (7838.3) | 0.821 | 10.8 (84.5) | 0.933 | -116.0 (107.6) | 0.715 | -3403.3 (10426.4) | 0.746 | -238.7 (110.5) | 0.381 | -95.9 (98.5) | 0.673 | 1543.8 (1828.8) | 0.578 | 21.5 (22.3) | 0.616 | -11.7 (25.7) | 0.678 |

*Note.* Reproductive-stage stratified analyses across all region families. Outliers removed (11 subjects; 3*SD* within sex and timepoint). Reproductive stage classified for female participants; hysterectomy (*n* = 1) and undetermined stage (*n* = 3) excluded owing to insufficient subgroup size; males shown separately. E = 17β-estradiol; T = testosterone; P = progesterone; *b* = unstandardized coefficient; *SE* = standard error; *n* = analytic sample per hormone within stage. Post-hoc: all regions FDR-corrected within family (p_FDR reported throughout). Prediction (Baseline→Year 1) controls for baseline volume. *$p$_FDR < .05, **$p$_FDR < .01, ***$p$_FDR < .001.

**Table S18.** Reproductive stage-stratified hormone-volume change-score associations for all regions, with outliers.

| | | E | | T | | P | |
|---|---|---|---|---|---|---|---|
| **Stage** | **Region** | *b (SE)* | *p* | *b (SE)* | *p* | *b (SE)* | *p* |
| | | *n* = 10 | | *n* = 9 | | *n* = 9 | |
| **Reproductive** | Left Crus I | -214.3 (118.1) | 0.258 | -4.1 (4.2) | 0.750 | 0.7 (1.2) | 0.700 |
| | Right Crus I | -171.8 (187.6) | 0.450 | -2.1 (6.1) | 0.750 | 1.9 (1.3) | 0.470 |
| | Left Crus II | -270.1 (329.6) | 0.450 | -4.7 (10.6) | 0.750 | 3.5 (2.3) | 0.470 |
| | Right Crus II | -663.9 (287.0) | 0.258 | -17.7 (10.6) | 0.678 | 1.5 (3.6) | 0.700 |
| | Left Lobule V | -59.9 (116.6) | 0.839 | -0.9 (3.5) | 0.805 | 0.6 (0.9) | 0.636 |
| | Right Lobule V | -113.1 (72.3) | 0.527 | -2.2 (2.6) | 0.659 | 0.4 (0.7) | 0.636 |
| | Left Lobule VI | -12.0 (75.0) | 0.880 | 2.2 (2.1) | 0.659 | 0.9 (0.4) | 0.445 |
| | Right Lobule VI | -158.4 (125.8) | 0.527 | -3.2 (4.3) | 0.659 | 1.2 (1.0) | 0.594 |
| | Left Hippocampus | 2.2 (47.6) | 0.984 | 0.7 (1.3) | 0.619 | 0.0 (0.4) | 0.927 |
| | Right Hippocampus | -41.0 (42.9) | 0.932 | -1.1 (1.1) | 0.619 | 0.1 (0.3) | 0.927 |
| | Left Amygdala | 37.8 (12.6) | 0.183 | 1.1 (0.4) | 0.288 | 0.0 (0.2) | 0.927 |
| | Right Amygdala | 0.5 (24.3) | 0.984 | 0.3 (0.6) | 0.619 | 0.0 (0.2) | 0.927 |
| | Left Thalamus | 71.7 (90.9) | 0.932 | 2.1 (2.4) | 0.619 | -0.7 (0.6) | 0.927 |
| | Right Thalamus | 46.6 (90.0) | 0.940 | 2.3 (2.9) | 0.619 | -0.3 (0.8) | 0.927 |
| | Left Lobule IX | -112.2 (43.8) | 0.253 | -2.2 (1.7) | 0.802 | 0.6 (0.4) | 0.376 |
| | Right Lobule IX | -117.2 (85.1) | 0.614 | -1.3 (2.9) | 0.802 | 1.2 (0.5) | 0.376 |
| | Vermis VI | -34.3 (9.5) | 0.152 | -0.8 (0.5) | 0.802 | 0.0 (0.2) | 0.883 |
| | Vermis VII | -12.5 (23.4) | 0.684 | -0.1 (0.8) | 0.939 | 0.2 (0.2) | 0.376 |
| | Vermis VIII | -26.4 (30.8) | 0.614 | -0.4 (0.9) | 0.802 | 0.2 (0.2) | 0.631 |
| | Vermis IX | -0.8 (21.9) | 0.971 | 0.5 (0.6) | 0.802 | 0.2 (0.1) | 0.376 |
| | Vermis X | 8.3 (8.6) | 0.614 | 0.3 (0.3) | 0.802 | 0.0 (0.1) | 0.883 |
| | Total Gray Matter | 2418.5 (3823.7) | 0.684 | 57.1 (129.3) | 0.802 | -41.9 (28.1) | 0.376 |
| | Cortical Volume | 4452.3 (4631.0) | 0.614 | 72.2 (164.1) | 0.802 | -61.9 (32.0) | 0.376 |
| | Cerebral White Matter | 5113.8 (4668.0) | 0.614 | 72.7 (148.3) | 0.802 | -62.4 (25.7) | 0.376 |
| | | *n* = 10 | | *n* = 10 | | *n* = 10 | |
| **Early Postmenopausal** | Left Crus I | -9.0 (282.4) | 0.976 | -10.9 (4.4) | 0.225 | -3.6 (4.4) | 0.855 |
| | Right Crus I | -334.5 (249.1) | 0.745 | -7.1 (6.0) | 0.515 | -6.6 (3.8) | 0.561 |
| | Left Crus II | 327.4 (523.3) | 0.745 | 1.2 (12.6) | 0.929 | 1.7 (9.0) | 0.855 |
| | Right Crus II | 275.7 (346.2) | 0.745 | 7.4 (7.9) | 0.515 | 2.3 (6.0) | 0.855 |
| | Left Lobule V | -26.0 (90.0) | 0.784 | -3.0 (1.6) | 0.256 | -0.9 (1.4) | 0.558 |
| | Right Lobule V | -53.8 (142.9) | 0.784 | -5.4 (2.3) | 0.256 | -1.4 (2.3) | 0.558 |
| | Left Lobule VI | -70.5 (98.0) | 0.784 | -0.8 (2.4) | 0.738 | -1.5 (1.6) | 0.558 |
| | Right Lobule VI | -177.6 (101.5) | 0.563 | 1.1 (3.0) | 0.738 | -2.1 (1.9) | 0.558 |
| | Left Hippocampus | 37.9 (52.6) | 0.524 | 0.4 (1.3) | 0.882 | 0.8 (0.8) | 0.445 |
| | Right Hippocampus | 20.3 (29.7) | 0.524 | 0.5 (0.7) | 0.882 | 0.7 (0.4) | 0.419 |
| | Left Amygdala | -72.3 (21.5) | 0.121 | -1.3 (0.7) | 0.607 | **-1.3 (0.2)** | **0.011*** |
| | Right Amygdala | -40.5 (21.8) | 0.365 | -0.3 (0.6) | 0.882 | -0.4 (0.4) | 0.445 |
| | Left Thalamus | 47.9 (66.2) | 0.524 | 0.3 (1.6) | 0.882 | 1.1 (1.0) | 0.445 |
| | Right Thalamus | 96.4 (71.7) | 0.474 | 0.7 (1.9) | 0.882 | 1.7 (1.2) | 0.419 |
| | Left Lobule IX | -29.3 (81.5) | 0.816 | -2.5 (1.5) | 0.406 | -0.7 (1.3) | 0.695 |
| | Right Lobule IX | -64.2 (53.0) | 0.400 | -1.7 (1.2) | 0.406 | -1.3 (0.8) | 0.255 |
| | Vermis VI | 40.0 (10.7) | 0.108 | 0.7 (0.4) | 0.406 | 0.6 (0.2) | 0.191 |
| | Vermis VII | 52.2 (19.9) | 0.158 | 0.7 (0.6) | 0.417 | 0.8 (0.4) | 0.217 |
| | Vermis VIII | -39.9 (12.1) | 0.108 | -0.8 (0.4) | 0.406 | -0.7 (0.2) | 0.191 |
| | Vermis IX | -1.1 (32.8) | 0.974 | -0.2 (0.8) | 0.848 | -0.3 (0.5) | 0.695 |

| Stage | Region | E b (SE) | E p | T b (SE) | T p | P b (SE) | P p |
|---|---|---|---|---|---|---|---|
| | Vermis X | 18.9 (12.4) | 0.360 | 0.3 (0.3) | 0.417 | 0.3 (0.2) | 0.217 |
| | Total Gray Matter | -5366.6 (3791.8) | 0.360 | -118.5 (89.7) | 0.406 | -113.9 (53.9) | 0.217 |
| | Cortical Volume | -5456.7 (3247.3) | 0.360 | -121.0 (77.2) | 0.406 | -98.4 (50.7) | 0.217 |
| | Cerebral White Matter | -2681.6 (4376.8) | 0.709 | 84.6 (98.3) | 0.476 | 0.8 (75.0) | 0.992 |
| | | *n* = 27 | | *n* = 27 | | *n* = 25 | |
| **Late Postmenopausal** | Left Crus I | -111.2 (64.1) | 0.194 | -3.2 (2.1) | 0.625 | -2.8 (1.1) | 0.083 |
| | Right Crus I | -101.6 (78.5) | 0.278 | -1.4 (2.7) | 0.939 | -2.3 (1.4) | 0.161 |
| | Left Crus II | -61.5 (67.4) | 0.371 | 0.1 (2.3) | 0.978 | 1.2 (1.2) | 0.327 |
| | Right Crus II | -115.2 (59.3) | 0.194 | 0.8 (2.1) | 0.939 | 1.6 (1.0) | 0.161 |
| | Left Lobule V | -3.4 (17.4) | 0.848 | -0.9 (0.5) | 0.213 | -0.5 (0.3) | 0.335 |
| | Right Lobule V | -21.9 (17.4) | 0.323 | **-1.4 (0.5)** | **0.037*** | -0.7 (0.3) | 0.122 |
| | Left Lobule VI | -45.9 (38.2) | 0.323 | 0.2 (1.3) | 0.906 | -0.2 (0.7) | 0.742 |
| | Right Lobule VI | -49.9 (38.8) | 0.323 | 0.7 (1.3) | 0.798 | 0.6 (0.6) | 0.442 |
| | Left Hippocampus | 8.5 (16.7) | 0.965 | -0.1 (0.6) | 0.925 | 0.0 (0.3) | 0.941 |
| | Right Hippocampus | -5.5 (13.9) | 0.965 | 0.1 (0.5) | 0.924 | 0.2 (0.3) | 0.941 |
| | Left Amygdala | -1.1 (13.7) | 0.965 | -0.2 (0.4) | 0.924 | 0.0 (0.3) | 0.941 |
| | Right Amygdala | -14.5 (11.1) | 0.965 | -0.1 (0.4) | 0.924 | -0.2 (0.2) | 0.941 |
| | Left Thalamus | 3.8 (38.8) | 0.965 | -0.5 (1.3) | 0.924 | -0.1 (0.7) | 0.941 |
| | Right Thalamus | 1.9 (42.6) | 0.965 | 0.6 (1.4) | 0.924 | 0.2 (0.8) | 0.941 |
| | Left Lobule IX | 2.4 (19.8) | 0.995 | -0.4 (0.6) | 0.692 | -0.6 (0.4) | 0.284 |
| | Right Lobule IX | 17.8 (23.4) | 0.995 | -1.0 (0.8) | 0.692 | -0.2 (0.5) | 0.989 |
| | Vermis VI | -2.5 (12.1) | 0.995 | -0.4 (0.4) | 0.692 | -0.4 (0.2) | 0.141 |
| | Vermis VII | -3.8 (6.8) | 0.995 | 0.4 (0.2) | 0.329 | 0.3 (0.1) | 0.103 |
| | Vermis VIII | -3.8 (12.1) | 0.995 | -0.7 (0.4) | 0.329 | -0.6 (0.2) | 0.103 |
| | Vermis IX | -3.0 (6.5) | 0.995 | -0.1 (0.2) | 0.692 | 0.0 (0.1) | 0.989 |
| | Vermis X | 0.0 (3.6) | 0.995 | 0.0 (0.1) | 0.692 | -0.1 (0.1) | 0.271 |
| | Total Gray Matter | 383.6 (2081.1) | 0.995 | 33.2 (68.3) | 0.692 | 0.5 (39.6) | 0.989 |
| | Cortical Volume | 475.0 (1765.1) | 0.995 | 32.4 (57.8) | 0.692 | 4.2 (33.7) | 0.989 |
| | Cerebral White Matter | 792.0 (1695.6) | 0.995 | -47.1 (55.2) | 0.692 | 0.5 (33.3) | 0.989 |
| | | *n* = 44 | | *n* = 43 | | *n* = 43 | |
| **Males** | Left Crus I | 189.9 (78.5) | 0.081 | 1.2 (1.0) | 0.600 | 1.5 (1.1) | 0.638 |
| | Right Crus I | 147.8 (73.9) | 0.105 | 0.9 (0.9) | 0.600 | 1.1 (1.0) | 0.638 |
| | Left Crus II | 48.9 (57.5) | 0.401 | 0.4 (0.7) | 0.600 | 0.4 (0.8) | 0.659 |
| | Right Crus II | 129.3 (86.9) | 0.193 | 0.6 (1.1) | 0.600 | 0.5 (1.2) | 0.659 |
| | Left Lobule V | 23.4 (28.1) | 0.430 | 0.1 (0.3) | 0.724 | 0.3 (0.4) | 0.497 |
| | Right Lobule V | 23.5 (29.5) | 0.430 | 0.4 (0.4) | 0.629 | 0.6 (0.4) | 0.273 |
| | Left Lobule VI | 71.5 (36.1) | 0.110 | 0.2 (0.5) | 0.724 | 0.7 (0.5) | 0.281 |
| | Right Lobule VI | **146.1 (48.2)** | **0.017*** | 1.2 (0.6) | 0.261 | 1.4 (0.7) | 0.184 |
| | Left Hippocampus | 10.2 (16.7) | 0.816 | -0.2 (0.2) | 0.819 | -0.1 (0.2) | 0.909 |
| | Right Hippocampus | 19.5 (17.8) | 0.816 | 0.0 (0.2) | 0.954 | 0.0 (0.2) | 0.909 |
| | Left Amygdala | 10.2 (13.6) | 0.816 | 0.2 (0.2) | 0.819 | 0.0 (0.2) | 0.909 |
| | Right Amygdala | 2.4 (11.5) | 0.958 | 0.0 (0.1) | 0.954 | 0.0 (0.2) | 0.909 |
| | Left Thalamus | 23.7 (32.8) | 0.816 | 0.1 (0.4) | 0.954 | 0.4 (0.4) | 0.909 |
| | Right Thalamus | 2.3 (42.8) | 0.958 | -0.4 (0.5) | 0.819 | 0.4 (0.5) | 0.909 |
| | Left Lobule IX | 48.9 (29.7) | 0.451 | 0.3 (0.4) | 0.895 | 0.1 (0.4) | 0.844 |
| | Right Lobule IX | 45.2 (26.8) | 0.451 | 0.3 (0.3) | 0.895 | 0.1 (0.4) | 0.844 |
| | Vermis VI | 12.3 (13.8) | 0.542 | 0.0 (0.2) | 0.973 | 0.1 (0.2) | 0.844 |

| Stage | Region | E b (SE) | E p | T b (SE) | T p | P b (SE) | P p |
|---|---|---|---|---|---|---|---|
| | Vermis VII | 1.1 (8.3) | 0.891 | 0.0 (0.1) | 0.973 | 0.0 (0.1) | 0.844 |
| | Vermis VIII | 15.4 (16.1) | 0.542 | 0.0 (0.2) | 0.985 | -0.1 (0.2) | 0.844 |
| | Vermis IX | 14.2 (9.3) | 0.451 | 0.1 (0.1) | 0.895 | 0.0 (0.1) | 0.844 |
| | Vermis X | 3.3 (3.7) | 0.542 | 0.1 (0.0) | 0.895 | 0.0 (0.1) | 0.844 |
| | Total Gray Matter | 546.2 (1818.5) | 0.851 | 17.6 (20.8) | 0.895 | 29.9 (23.7) | 0.844 |
| | Cortical Volume | -1286.1 (1668.0) | 0.557 | 8.3 (19.5) | 0.973 | 22.9 (22.1) | 0.844 |
| | Cerebral White Matter | 1044.0 (1158.8) | 0.542 | -2.3 (13.7) | 0.973 | 7.3 (15.7) | 0.844 |

*Note.* Reproductive-stage stratified change-score analyses (ΔHormone predicting ΔVolume) across all region families. With outliers retained. Hysterectomy (*n* = 1) and undetermined stage (*n* = 3) excluded; males shown separately. Post-hoc: all regions FDR-corrected within family. **p*_FDR < .05.

**Table S19.** Reproductive stage-stratified hormone-volume change-score associations for all regions, without outliers.

| | | E | | T | | P | |
|---|---|---|---|---|---|---|---|
| **Stage** | **Region** | *b (SE)* | *p* | *b (SE)* | *p* | *b (SE)* | *p* |
| | | *n* = 7 | | *n* = 6 | | *n* = 6 | |
| | Left Crus I | 119.5 (65.4) | 0.837 | -5.8 (10.7) | 0.912 | -0.5 (0.2) | 0.566 |
| | Right Crus I | -271.8 (303.5) | 0.863 | -0.4 (17.8) | 0.986 | -0.5 (0.6) | 0.719 |
| | Left Crus II | -314.0 (589.1) | 0.863 | -35.0 (23.8) | 0.760 | -0.5 (1.8) | 0.827 |
| | Right Crus II | -65.6 (485.9) | 0.905 | -44.7 (11.8) | 0.654 | -1.9 (0.9) | 0.566 |
| | Left Lobule V | -44.3 (212.8) | 0.854 | -19.4 (10.7) | 0.640 | -0.4 (0.9) | 0.812 |
| | Right Lobule V | -65.9 (137.8) | 0.854 | -1.1 (17.2) | 0.960 | -0.5 (0.6) | 0.812 |
| | Left Lobule VI | -203.8 (85.7) | 0.562 | 26.8 (8.8) | 0.640 | 0.7 (1.1) | 0.812 |
| | Right Lobule VI | -132.8 (269.5) | 0.854 | -13.8 (32.1) | 0.960 | 0.5 (1.5) | 0.812 |
| | Left Hippocampus | -59.7 (19.4) | 0.449 | 3.9 (3.1) | 0.616 | 0.2 (0.0) | 0.065 |
| | Right Hippocampus | -60.8 (94.5) | 0.703 | -4.4 (1.6) | 0.616 | -0.1 (0.2) | 0.933 |
| | Left Amygdala | 48.5 (21.2) | 0.449 | -4.7 (2.7) | 0.616 | -0.1 (0.2) | 0.933 |
| **Reproductive** | Right Amygdala | -46.2 (45.4) | 0.624 | 4.9 (7.1) | 0.616 | 0.0 (0.4) | 0.937 |
| | Left Thalamus | 255.4 (164.8) | 0.523 | -7.0 (5.8) | 0.616 | **-0.4 (0.0)** | **0.038*** |
| | Right Thalamus | 25.1 (136.2) | 0.871 | 9.6 (10.4) | 0.616 | 0.6 (0.1) | 0.157 |
| | Left Lobule IX | -155.2 (77.7) | 0.411 | 6.7 (7.4) | 0.744 | 0.4 (0.1) | 0.641 |
| | Right Lobule IX | -218.7 (103.7) | 0.411 | 10.3 (4.0) | 0.744 | 0.5 (0.2) | 0.641 |
| | Vermis VI | -25.8 (3.2) | 0.150 | 2.3 (0.2) | 0.542 | 0.1 (0.1) | 0.785 |
| | Vermis VII | -37.8 (29.5) | 0.469 | 2.2 (3.3) | 0.744 | 0.0 (0.2) | 0.931 |
| | Vermis VIII | -34.5 (51.6) | 0.715 | -1.4 (2.8) | 0.744 | -0.1 (0.1) | 0.641 |
| | Vermis IX | -32.4 (24.8) | 0.469 | 1.7 (2.5) | 0.744 | 0.0 (0.1) | 0.931 |
| | Vermis X | 25.1 (10.3) | 0.411 | -2.4 (1.3) | 0.744 | 0.0 (0.1) | 0.931 |
| | Total Gray Matter | 539.7 (4881.7) | 0.922 | 379.7 (664.1) | 0.744 | -6.2 (33.5) | 0.931 |
| | Cortical Volume | 2955.6 (6197.6) | 0.756 | 321.2 (755.6) | 0.744 | -10.8 (34.9) | 0.931 |
| | Cerebral White Matter | 12546.0 (6777.8) | 0.411 | -498.0 (225.1) | 0.744 | -23.4 (6.7) | 0.641 |
| | | *n* = 10 | | *n* = 10 | | *n* = 10 | |
| | Left Crus I | -9.0 (282.4) | 0.976 | -10.9 (4.4) | 0.225 | -3.6 (4.4) | 0.855 |
| | Right Crus I | -334.5 (249.1) | 0.745 | -7.1 (6.0) | 0.515 | -6.6 (3.8) | 0.561 |
| | Left Crus II | 327.4 (523.3) | 0.745 | 1.2 (12.6) | 0.929 | 1.7 (9.0) | 0.855 |
| | Right Crus II | 275.7 (346.2) | 0.745 | 7.4 (7.9) | 0.515 | 2.3 (6.0) | 0.855 |
| | Left Lobule V | -26.0 (90.0) | 0.784 | -3.0 (1.6) | 0.256 | -0.9 (1.4) | 0.558 |
| | Right Lobule V | -53.8 (142.9) | 0.784 | -5.4 (2.3) | 0.256 | -1.4 (2.3) | 0.558 |
| | Left Lobule VI | -70.5 (98.0) | 0.784 | -0.8 (2.4) | 0.738 | -1.5 (1.6) | 0.558 |
| | Right Lobule VI | -177.6 (101.5) | 0.563 | 1.1 (3.0) | 0.738 | -2.1 (1.9) | 0.558 |
| | Left Hippocampus | 37.9 (52.6) | 0.524 | 0.4 (1.3) | 0.882 | 0.8 (0.8) | 0.445 |
| **Early Postmenopausal** | Right Hippocampus | 20.3 (29.7) | 0.524 | 0.5 (0.7) | 0.882 | 0.7 (0.4) | 0.419 |
| | Left Amygdala | -72.3 (21.5) | 0.121 | -1.3 (0.7) | 0.607 | **-1.3 (0.2)** | **0.011*** |
| | Right Amygdala | -40.5 (21.8) | 0.365 | -0.3 (0.6) | 0.882 | -0.4 (0.4) | 0.445 |
| | Left Thalamus | 47.9 (66.2) | 0.524 | 0.3 (1.6) | 0.882 | 1.1 (1.0) | 0.445 |
| | Right Thalamus | 96.4 (71.7) | 0.474 | 0.7 (1.9) | 0.882 | 1.7 (1.2) | 0.419 |
| | Left Lobule IX | -29.3 (81.5) | 0.816 | -2.5 (1.5) | 0.406 | -0.7 (1.3) | 0.695 |
| | Right Lobule IX | -64.2 (53.0) | 0.400 | -1.7 (1.2) | 0.406 | -1.3 (0.8) | 0.255 |
| | Vermis VI | 40.0 (10.7) | 0.108 | 0.7 (0.4) | 0.406 | 0.6 (0.2) | 0.191 |
| | Vermis VII | 52.2 (19.9) | 0.158 | 0.7 (0.6) | 0.417 | 0.8 (0.4) | 0.217 |
| | Vermis VIII | -39.9 (12.1) | 0.108 | -0.8 (0.4) | 0.406 | -0.7 (0.2) | 0.191 |
| | Vermis IX | -1.1 (32.8) | 0.974 | -0.2 (0.8) | 0.848 | -0.3 (0.5) | 0.695 |

| Stage | Region | E | | T | | P | |
|---|---|---|---|---|---|---|---|
| | | b (SE) | p | b (SE) | p | b (SE) | p |
| | Vermis X | 18.9 (12.4) | 0.360 | 0.3 (0.3) | 0.417 | 0.3 (0.2) | 0.217 |
| | Total Gray Matter | -5366.6 (3791.8) | 0.360 | -118.5 (89.7) | 0.406 | -113.9 (53.9) | 0.217 |
| | Cortical Volume | -5456.7 (3247.3) | 0.360 | -121.0 (77.2) | 0.406 | -98.4 (50.7) | 0.217 |
| | Cerebral White Matter | -2681.6 (4376.8) | 0.709 | 84.6 (98.3) | 0.476 | 0.8 (75.0) | 0.992 |
| | | *n* = 26 | | *n* = 26 | | *n* = 25 | |
| Late Postmenopausal | Left Crus I | -142.9 (146.4) | 0.965 | -2.9 (2.1) | 0.739 | -2.8 (1.1) | 0.083 |
| | Right Crus I | 7.8 (177.4) | 0.965 | -1.1 (2.6) | 0.887 | -2.3 (1.4) | 0.161 |
| | Left Crus II | 88.8 (149.6) | 0.965 | 0.3 (2.2) | 0.887 | 1.2 (1.2) | 0.327 |
| | Right Crus II | -23.7 (133.7) | 0.965 | 1.2 (2.0) | 0.887 | 1.6 (1.0) | 0.161 |
| | Left Lobule V | 17.4 (39.5) | 0.664 | -0.9 (0.6) | 0.243 | -0.5 (0.3) | 0.335 |
| | Right Lobule V | -94.0 (35.6) | 0.061 | **-1.4 (0.5)** | **0.045*** | -0.7 (0.3) | 0.122 |
| | Left Lobule VI | -89.4 (86.6) | 0.418 | 0.2 (1.3) | 0.852 | -0.2 (0.7) | 0.742 |
| | Right Lobule VI | -92.4 (88.0) | 0.418 | 0.8 (1.3) | 0.730 | 0.6 (0.6) | 0.442 |
| | Left Hippocampus | 50.9 (36.7) | 0.270 | 0.0 (0.6) | 0.934 | 0.0 (0.3) | 0.941 |
| | Right Hippocampus | 48.6 (28.9) | 0.215 | 0.2 (0.5) | 0.930 | 0.2 (0.3) | 0.941 |
| | Left Amygdala | -0.1 (31.3) | 0.997 | -0.2 (0.5) | 0.930 | 0.0 (0.3) | 0.941 |
| | Right Amygdala | -17.7 (25.3) | 0.591 | -0.1 (0.4) | 0.930 | -0.2 (0.2) | 0.941 |
| | Left Thalamus | 170.8 (78.9) | 0.215 | -0.4 (1.3) | 0.930 | -0.1 (0.7) | 0.941 |
| | Right Thalamus | 162.6 (89.2) | 0.215 | 0.8 (1.4) | 0.930 | 0.2 (0.8) | 0.941 |
| | Left Lobule IX | 18.5 (45.2) | 0.980 | -0.4 (0.7) | 0.740 | -0.6 (0.4) | 0.284 |
| | Right Lobule IX | 80.0 (51.3) | 0.735 | -1.0 (0.8) | 0.732 | -0.2 (0.5) | 0.989 |
| | Vermis VI | 5.3 (27.6) | 0.984 | -0.4 (0.4) | 0.740 | -0.4 (0.2) | 0.141 |
| | Vermis VII | -9.2 (15.4) | 0.932 | 0.4 (0.2) | 0.380 | 0.3 (0.1) | 0.103 |
| | Vermis VIII | 0.6 (27.6) | 0.984 | -0.7 (0.4) | 0.380 | -0.6 (0.2) | 0.103 |
| | Vermis IX | 1.9 (14.8) | 0.984 | -0.1 (0.2) | 0.740 | 0.0 (0.1) | 0.989 |
| | Vermis X | 11.6 (7.8) | 0.735 | 0.0 (0.1) | 0.740 | -0.1 (0.1) | 0.271 |
| | Total Gray Matter | 5184.8 (4611.0) | 0.735 | 35.6 (69.9) | 0.740 | 0.5 (39.6) | 0.989 |
| | Cortical Volume | 4229.5 (3929.8) | 0.735 | 33.8 (59.3) | 0.740 | 4.2 (33.7) | 0.989 |
| | Cerebral White Matter | 3513.3 (3818.9) | 0.736 | -47.7 (56.7) | 0.740 | 0.5 (33.3) | 0.989 |
| | | *n* = 39 | | *n* = 38 | | *n* = 38 | |
| Males | Left Crus I | 185.5 (106.2) | 0.120 | 0.2 (1.5) | 0.905 | 0.1 (1.2) | 0.975 |
| | Right Crus I | 232.3 (94.9) | 0.079 | 1.9 (1.3) | 0.628 | 1.0 (1.1) | 0.975 |
| | Left Crus II | 54.6 (84.5) | 0.523 | -0.2 (1.1) | 0.905 | 0.0 (1.0) | 0.975 |
| | Right Crus II | 242.3 (125.1) | 0.120 | 0.5 (1.7) | 0.905 | 0.0 (1.5) | 0.975 |
| | Left Lobule V | 17.3 (41.4) | 0.679 | -0.1 (0.5) | 0.803 | 0.0 (0.5) | 0.934 |
| | Right Lobule V | 48.2 (39.6) | 0.309 | 0.3 (0.5) | 0.708 | 0.4 (0.4) | 0.523 |
| | Left Lobule VI | 87.5 (46.9) | 0.141 | 0.8 (0.6) | 0.607 | 1.0 (0.5) | 0.152 |
| | Right Lobule VI | 144.2 (66.8) | 0.141 | 1.0 (0.9) | 0.607 | 1.4 (0.8) | 0.152 |
| | Left Hippocampus | 20.8 (23.2) | 0.445 | 0.1 (0.3) | 0.821 | 0.0 (0.3) | 0.902 |
| | Right Hippocampus | 33.5 (25.7) | 0.445 | 0.1 (0.3) | 0.821 | 0.1 (0.3) | 0.902 |
| | Left Amygdala | 16.5 (19.2) | 0.445 | 0.2 (0.3) | 0.821 | -0.1 (0.2) | 0.902 |
| | Right Amygdala | 13.1 (15.7) | 0.445 | 0.1 (0.2) | 0.821 | 0.1 (0.2) | 0.902 |
| | Left Thalamus | 35.6 (46.1) | 0.445 | 0.6 (0.6) | 0.821 | 0.6 (0.5) | 0.666 |
| | Right Thalamus | 45.6 (59.1) | 0.445 | -0.3 (0.7) | 0.821 | 0.8 (0.6) | 0.666 |
| | Left Lobule IX | 74.0 (42.3) | 0.326 | 0.3 (0.6) | 0.869 | 0.2 (0.5) | 0.948 |
| | Right Lobule IX | 57.1 (35.9) | 0.326 | 0.1 (0.5) | 0.917 | 0.1 (0.4) | 0.948 |
| | Vermis VI | 40.6 (17.5) | 0.261 | 0.0 (0.2) | 0.917 | 0.1 (0.2) | 0.948 |

| Stage | Region | E b (SE) | E p | T b (SE) | T p | P b (SE) | P p |
|---|---|---|---|---|---|---|---|
| | Vermis VII | 6.8 (12.1) | 0.688 | 0.0 (0.2) | 0.917 | 0.0 (0.1) | 0.948 |
| | Vermis VIII | 24.2 (22.9) | 0.465 | -0.2 (0.3) | 0.869 | 0.0 (0.3) | 0.948 |
| | Vermis IX | 17.3 (12.7) | 0.361 | 0.2 (0.2) | 0.838 | 0.0 (0.1) | 0.948 |
| | Vermis X | 4.6 (4.6) | 0.465 | 0.0 (0.1) | 0.869 | 0.0 (0.1) | 0.948 |
| | Total Gray Matter | 3472.8 (2240.7) | 0.326 | 41.8 (28.3) | 0.838 | 43.0 (23.7) | 0.627 |
| | Cortical Volume | 919.4 (2271.2) | 0.688 | 33.1 (28.4) | 0.838 | 37.4 (23.8) | 0.627 |
| | Cerebral White Matter | 751.0 (1678.5) | 0.688 | 11.2 (21.6) | 0.869 | 20.0 (18.2) | 0.931 |

*Note.* Reproductive-stage stratified change-score analyses (ΔHormone predicting ΔVolume) across all region families. Outliers removed (11 subjects; 3*SD* within sex and timepoint). Hysterectomy (*n* = 1) and undetermined stage (*n* = 3) excluded; males shown separately. Post-hoc: all regions FDR-corrected within family. *$p$_FDR < .05.
